# LetA defines a structurally distinct transporter family involved in lipid trafficking

**DOI:** 10.1101/2025.03.21.644421

**Authors:** Cristina C. Santarossa, Yupeng Li, Sara Yousef, Hale S. Hasdemir, Carlos C. Rodriguez, Max B. Haase, Minkyung Baek, Nicolas Coudray, John G. Pavek, Kimber N. Focke, Annika L. Silverberg, Carmelita Bautista, Johannes Yeh, Michael T. Marty, David Baker, Emad Tajkhorshid, Damian C. Ekiert, Gira Bhabha

## Abstract

Membrane transport proteins translocate diverse cargos, ranging from small sugars to entire proteins, across cellular membranes. A few structurally distinct protein families have been described that account for most of the known membrane transport processes. However, many membrane proteins with predicted transporter functions remain uncharacterized. We determined the structure of *E. coli* LetAB, a phospholipid transporter involved in outer membrane integrity, and found that LetA adopts a distinct architecture that is structurally and evolutionarily unrelated to known transporter families. LetA functions as a pump at one end of a ~225 Å long tunnel formed by its binding partner, MCE protein LetB, creating a pathway for lipid transport between the inner and outer membranes. Unexpectedly, the LetA transmembrane domains adopt a fold that is evolutionarily related to the eukaryotic tetraspanin family of membrane proteins, including TARPs and claudins. LetA has no detectable homology to known transport proteins, and defines a new class of membrane transporters. Through a combination of deep mutational scanning, molecular dynamics simulations, AlphaFold-predicted alternative states, and functional studies, we present a model for how the LetA-like family of membrane transporters may use energy from the proton-motive force to drive the transport of lipids across the bacterial cell envelope.

## INTRODUCTION

Membrane transport is critical for fundamental cellular processes, including cell growth, division and homeostasis. Transporters can mediate either the active or passive transport of a wide variety of substrates across cellular membranes, such as nutrients, ions, and drugs^1–3^. To date, several transporter folds have been identified^4–8^, including members of the ATP-binding cassette (ABC) and solute carrier families, among others. However, many membrane proteins hypothesized to be transporters remain uncharacterized, even in well-studied model organisms such as *Escherichia coli*. Some of these hypothetical transporters may be evolutionarily related to known transporter families, but have diverged beyond recognition at the sequence level. Alternatively, these unstudied protein families may represent new kinds of transporters that await experimental characterization.

The Mammalian Cell Entry (MCE) family of proteins has been implicated in lipid transport across the cell envelope in double-membraned bacteria^9–11^ and between the ER and chloroplasts in plants^12,13^. Members of this family play an important role in maintaining the cell envelope of Gram-negative bacteria^10,14^, and in the scavenging of host lipids in *Mycobacteria*^*11*^. To carry out their cellular function, MCE proteins adopt a range of architectures that form pathways between membranes^9,15^. MCE proteins are usually associated with other membrane proteins, which are thought to drive lipid translocation through pores formed at the center of hexameric rings formed by MCE proteins. The best characterized MCE systems interact with ABC transporters to drive substrate translocation, and include the *E. coli* <u>M</u>aintenance of outer membrane <u>L</u>ipid <u>A</u>symmetry (Mla)^9,10,16–23^, the plant TRIGALACTOSYLDIACYLGLYCEROL (TGD) pathways^12,13,24^, and the Mycobacterial Mce1^15,25–28^ and Mce4 systems^11,25,29,30^. However, many other MCE gene clusters do not encode components of an ABC transporter, and it is unknown if and how lipids are translocated, or how transport is energized.

LetB (<u>L</u>ipophilic <u>E</u>nvelope-spanning <u>T</u>unnel B) is a large MCE protein that forms a hydrophobic tunnel long enough to span the periplasm between the inner membrane (IM) and outer membrane (OM) in *E. coli*^*31*^ (Fig. 1a). The prevailing model in the field is that LetB transports lipids between membranes through a central hydrophobic tunnel^9,14,31–33^. The mechanism by which lipids enter the LetB tunnel remains unknown. LetB is encoded in an operon together with LetA (Fig. 1a), a multipass transmembrane (TM) protein with no detectable homology to transporter families at the primary sequence level. In Gram-negative bacteria, proteins homologous to LetA are encoded adjacent to some classes of MCE proteins, including the <u>P</u>ara<u>q</u>uat Inducible (Pqi) system in *E. coli*, suggesting that these proteins may have evolved to function together in lipid transport. LetA is poised to act as a pump for substrate translocation through the LetB tunnel and has the potential to define a new class of membrane transport proteins.

**Fig. 1.**
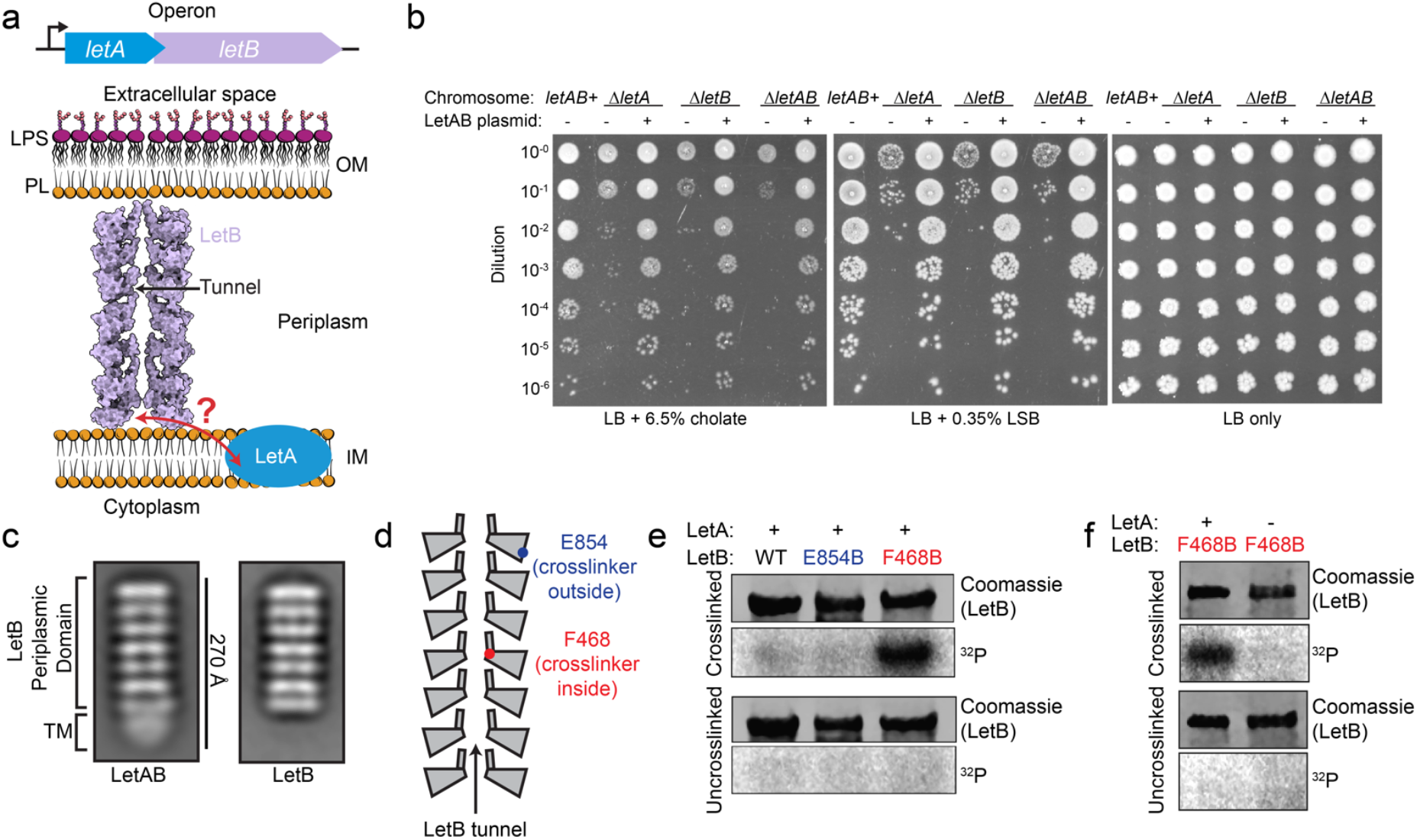
LetA and LetB are functionally linked and physically interact. **a**, Schematic of the *letAB* operon, and model of LetA and LetB in the cell envelope. A cross-section of LetB (PDB 6V0C) is shown, oriented in the context of the IM and OM. **b**, Cellular assay to assess the function of *letA, letB*, or *letAB* deletion mutants. 10-fold serial dilutions of strains were spotted on LB agar with or without the indicated detergent. All strains are constructed in a Δ*pqiAB* background. **c**, 2D class averages from negative stain EM data for LetAB and the periplasmic domain of LetB alone. **d**, Schematic of LetB, with spheres showing positions of residues targeted for incorporation of the photo crosslinking amino acid, BPA. F468 is located inside the LetB tunnel, and E854 is located outside the tunnel. **e**, SDS-PAGE analysis of purified LetAB without BPA incorporation (WT) or with BPA incorporated at position 468 or 854. Samples were either UV crosslinked *in vivo* or uncrosslinked, and the SDS-PAGE gel was stained with Coomassie (LetB) or phosphor-imaged (^32^P signal). **f**, SDS-PAGE analysis of purified LetB with BPA incorporated at position 468, with or without co-expression of LetA, prepared as in (**e**).

Here, we show that LetA is functionally linked to LetB, and that the two proteins physically interact. We report the structure of the LetAB complex and find that LetA is distantly related to the eukaryotic tetraspanin superfamily of membrane proteins, which are not known to have intrinsic transporter activity, including the Transmembrane AMPA receptor Regulatory Proteins (TARPs), claudins, and vitamin K epoxide reductase (VKOR). Our structure, together with deep mutational scanning, molecular dynamics (MD) simulations, and AlphaFold predictions of alternative states coupled with experimental validation, lead to a model for how LetA may drive phospholipid transport to maintain OM integrity in *E. coli*, providing insights into a previously uncharacterized family of transporters in bacteria.

## RESULTS

### LetA and LetB are functionally linked and form a complex

Deletion of *letA* and *letB* together (*ΔletAB*) in *E. coli* was previously shown to cause mild sensitivity to the bile salt, cholate, and the zwitterionic surfactant, lauryl sulfobetaine (LSB)^31,34^. Both phenotypes are exacerbated when *pqiAB*, a second *E. coli* MCE system, is also deleted (*ΔpqiAB ΔletAB*)^31,34^. To assess the relative contributions of *letA* and *letB* to cholate and LSB sensitivity, we deleted *letA* and *letB* individually in a *ΔpqiAB* background. Strains lacking *letA* or *letB* exhibit similar growth defects to each other and to *ΔletAB* mutants, which can be rescued by complementation with a plasmid carrying WT *letAB* (Fig. 1b). These results suggest that LetA and LetB likely function in the same pathway.

To examine if LetA and LetB physically interact, we co-expressed both proteins in *E. coli*, pulled down His-tagged LetA, and found that untagged LetB is pulled down in a LetA-dependent manner (Extended Data Feig. 1a, Supplementary Feigure 1a) and can form a large complex with an apparent molecular weight of ~670 kDa (Extended Data Feigs. 1b-c, Supplementary Feigure 1b). Negative stain EM of purified LetAB shows particles with seven characteristic bands of density resembling LetB^31^, and an additional globular density at one end (Fig. 1c), which we hypothesized corresponds to LetA and the TM helices of LetB surrounded by a detergent micelle. Overall, these data show that LetA and LetB interact to form a stable complex.

### LetA is required for lipid entry into LetB tunnel

Previous studies have shown that the soluble, periplasmic domain of LetB binds phospholipids^9^, and crosslinking experiments in *E. coli* lysates suggest that the binding sites for phospholipid substrates are in the LetB central tunnel^31^. It is unclear, however, if phospholipids spontaneously enter the tunnel of the full-length, membrane-embedded LetAB complex *in vivo*. To address this question, we used an *in vivo* crosslinking assay. We grew *E. coli* in the presence of ^32^P orthophosphate, to label bulk phospholipids and other phosphate-containing molecules, and over-expressed LetAB with the unnatural photocrosslinking amino acid *p*-benzoyl-L-phenylalanine (BPA) incorporated at specific sites in LetB. We then irradiated live cells with UV light to allow *in vivo* crosslinking of molecules in proximity to the site of the BPA probe. Following purification of LetAB complexes, we analyzed the crosslinking of ^32^P-labeled molecules to LetB by electrophoresis and phosphorimaging. BPA was positioned inside the LetB tunnel (F468) or on the periplasm-facing exterior surface (E854, negative control) (Fig. 1d), locations validated in previous work^31^. We detect strong ^32^P incorporation into LetB when BPA was positioned inside the tunnel, but minimal ^32^P incorporation with BPA positioned outside the tunnel, as expected (Fig. 1e, Supplementary Feigure 1c). To test if lipid access to the LetB tunnel is dependent on LetA, we assessed lipid crosslinking in the LetB tunnel with or without co-expression of LetA. Efficient crosslinking was observed when LetA and LetB were co-expressed, but crosslinking was greatly reduced when LetA was not co-expressed (Fig. 1f, Supplementary Feigure 1d). LetB is localized to the membrane whether LetA is co-expressed or not (Extended Data Feig. 1d, Supplementary Feigure 1e), suggesting that its correct cellular localization is independent of LetA. Taken together, these results suggest that LetA is necessary for phospholipid entry into the tunnel of full-length LetB *in vivo*. The simplest interpretation of this result is that LetA is an exporter that loads lipids from the IM into LetB.

### Overall structure of LetAB complex

To understand how LetA and LetB interact, we determined the structure of the LetAB complex using single particle cryo-EM. We determined two structures of the LetAB complex, one in the presence of a crosslinker, glutaraldehyde, and the other without crosslinker. The crosslinked and uncrosslinked datasets yielded maps with similar average resolutions across the LetAB complex (2.5-4.6 Å: Fig. 2a, Extended Data Feig. 2, Supplementary Table 1), and the overall backbone conformation of LetA is similar in both structures (root-mean square displacement (RMSD) of 0.787 Å across 3123 atoms). However, as the TM region is better resolved in the presence of the crosslinker (Extended Data Feigs. 2C, 2H), we will primarily focus our discussion on the crosslinked LetAB structure, except where noted.

**Fig. 2.**
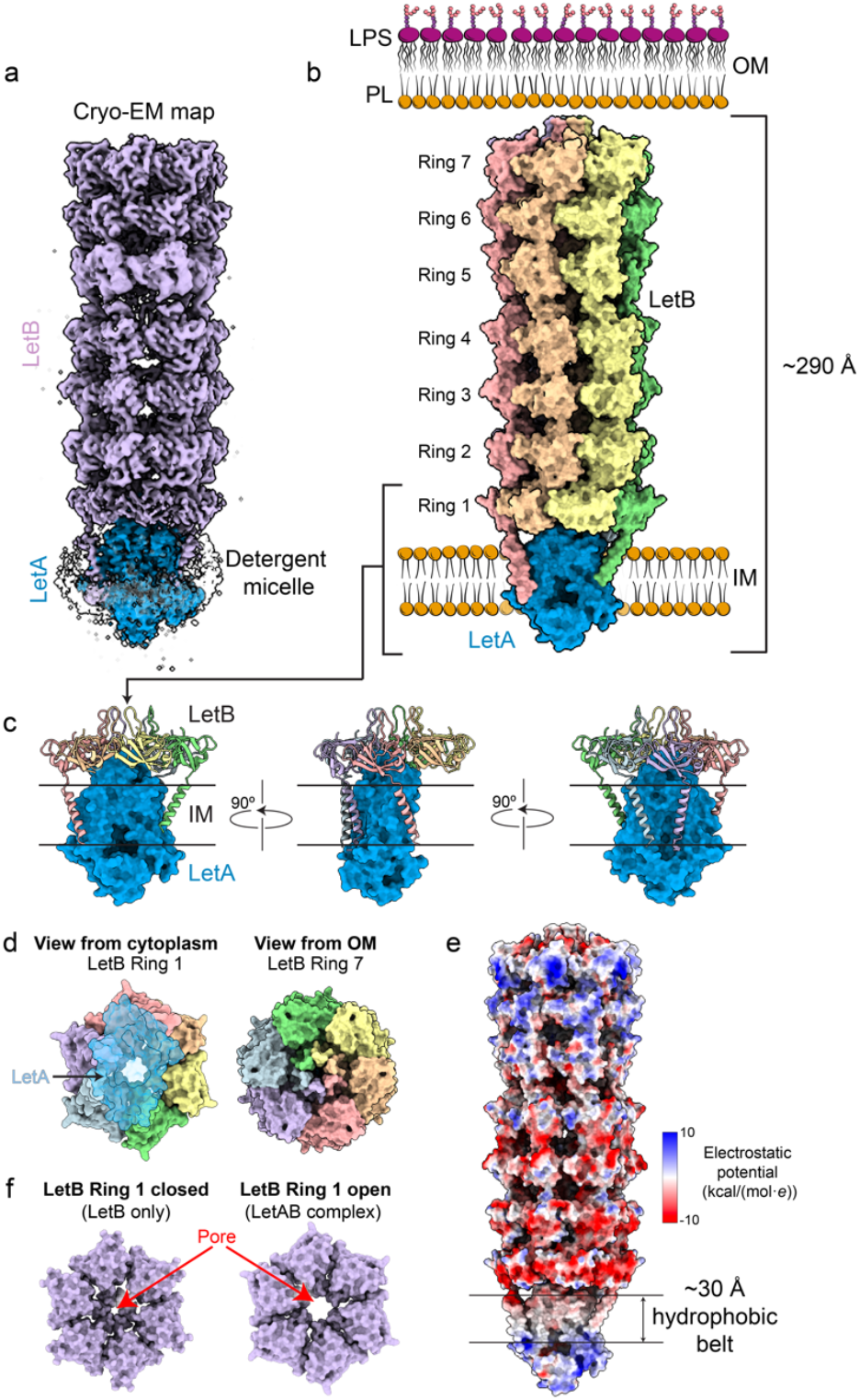
Cryo-EM structure of the LetAB complex. **a**, Composite map of crosslinked LetAB (Map 1). LetA (blue), LetB (purple) and the detergent micelle (white) are indicated. The contour level of the map is set to 2.37. **b**, Surface representation of the LetAB structure oriented in the context of the IM and OM. LetB monomers are depicted in different colors. The LetAB complex is ~290 Å long, and LetA is embedded in the IM. Membrane phospholipids (PL) and lipopolysaccharides (LPS) are shown. **c**, Structure of LetA (blue surface) in complex with MCE Ring 1 and LetB transmembrane helices, (cartoon). LetB monomers are colored as in (**b**). **d**, Views of the LetAB complex from the cytoplasm and from the OM, shown as surface representations. The LetA surface is partially transparent (blue), to illustrate its proximity to the LetB tunnel entrance. **e**, Molecular surface of the LetAB complex (Map 1) colored by electrostatic potential. **f**, Surface representation of MCE Ring 1 (purple) in the closed (PDB 6V0C) or open (Map 1) state. The view from the cytoplasm is shown.

The LetAB complex is an elongated assembly (Figs. 2a-b), ~290 Å long and ~90 Å wide, and consists of six copies of LetB and one copy of LetA. The periplasmic region of LetB hexamerizes to form the tunnel, consistent with previous structures^31,34^, and dominates the assembly, accounting for ~225 Å of the total length and 92% of the mass. The seven LetB MCE domains associate with neighboring protomers to form seven MCE rings, which stack to create a central hydrophobic tunnel that runs through the entire length of the LetB periplasmic domain (Extended Data Feigs. 1e-f). Each LetB protomer contains a single N-terminal TM helix, resulting in a total of six TM helices that anchor the assembly in the IM. A single copy of LetA associates with the LetB hexamer in the IM, making interactions with MCE Ring 1 and the TM helices of LetB (Figs. 2c-d). Consistent with its membrane localization, LetA contains a ~30 Å hydrophobic belt around its circumference, which we would expect to be embedded in the IM (Fig. 2e). Notably, density for only four of the six LetB TM helices are apparent in the EM map, and these four helices interact with LetA. The remaining two TM helices are not resolved (Fig. 2c) and may not stably interact with LetA, resulting in pronounced asymmetry in the TM region of the complex.

The wall of the LetB central tunnel is formed by pore-lining loops that emerge from each MCE domain^31^. Previous cryo-EM structures have shown that the pore-lining loops from MCE Rings 1, 5, 6 and 7 can adopt open and closed conformations, which control the diameter of the central tunnel, thereby potentially regulating the passage of substrates. LetA is positioned directly underneath the pore of MCE Ring 1. In the absence of LetA, MCE Ring 1 of LetB is predominantly in the closed state, in which the pore through the ring is not wide enough to allow passage of a phospholipid (Fig. 2f)^31^. Interestingly, in our structure of the LetAB complex, MCE Ring 1 of LetB adopts an open state, suggesting that binding to LetA modulates the conformation of the LetB tunnel (Fig. 2f).

### Overall structure of LetA

*E. coli* LetA is a single polypeptide that consists of two related modules, which we term “LetA modules”. Each LetA module consists of a cytoplasmic Zinc Ribbon (ZnR) domain followed by a transmembrane domain (TMD) (Figs. 3a-c). These modules are widespread across Proteobacteria^14^, and are either found in a single gene, as in *E. coli*, or encoded by two separate genes (e.g. *Pseudomonas aeruginosa*). Interestingly, the *E. coli* LetA modules can form a functional heterodimer when co-expressed from separate genes (split-LetA) (Extended Data Feigs. 3a-b, Supplementary Feigure 1f). The TMD of each LetA module contains four TM helices, one interfacial helix at the membrane-periplasm boundary, and a three-stranded β-sheet extending into the periplasm (Figs. 3a-d). The two LetA modules, which share only ~25% sequence identity, associate in a head-to-head manner in the structure, resulting in an intramolecular dimer with 2-fold pseudo-symmetry (Fig. 3e). The two TMDs form an inverted V-shape, creating a large, hydrophilic cleft that faces the cytoplasm (Fig. 3f). In this cleft, we observed the presence of a ^361^GRWSM-Ѱ-D-Ѱ-F^369^ motif (where Ѱ denotes an aliphatic amino acid: L, I, V, or M) that is well conserved in the C-terminal LetA module across a diverse set of LetA-like proteins (Extended Data Feigs. 3c-e). A similar motif is also present in the N-terminal LetA module, but is less strongly conserved. In addition to this cleft, LetA contains a periplasmic pocket 174 Å^3^ in volume (Fig. 3f), which is amphipathic and formed primarily by residues of TMD^C^, along with TM3 of TMD^N^. This periplasmic pocket sits directly below the entrance to the LetB tunnel, with the LetA periplasmic β-sheets creating a hydrophobic bridge that connects the pocket to the pore lining loops of MCE Ring 1 (Fig. 3g). In contrast, an equivalent pocket is not present in TMD^N^. The cleft and periplasmic pocket could potentially serve as substrate binding sites, and function as part of the substrate translocation pathway.

**Fig. 3.**
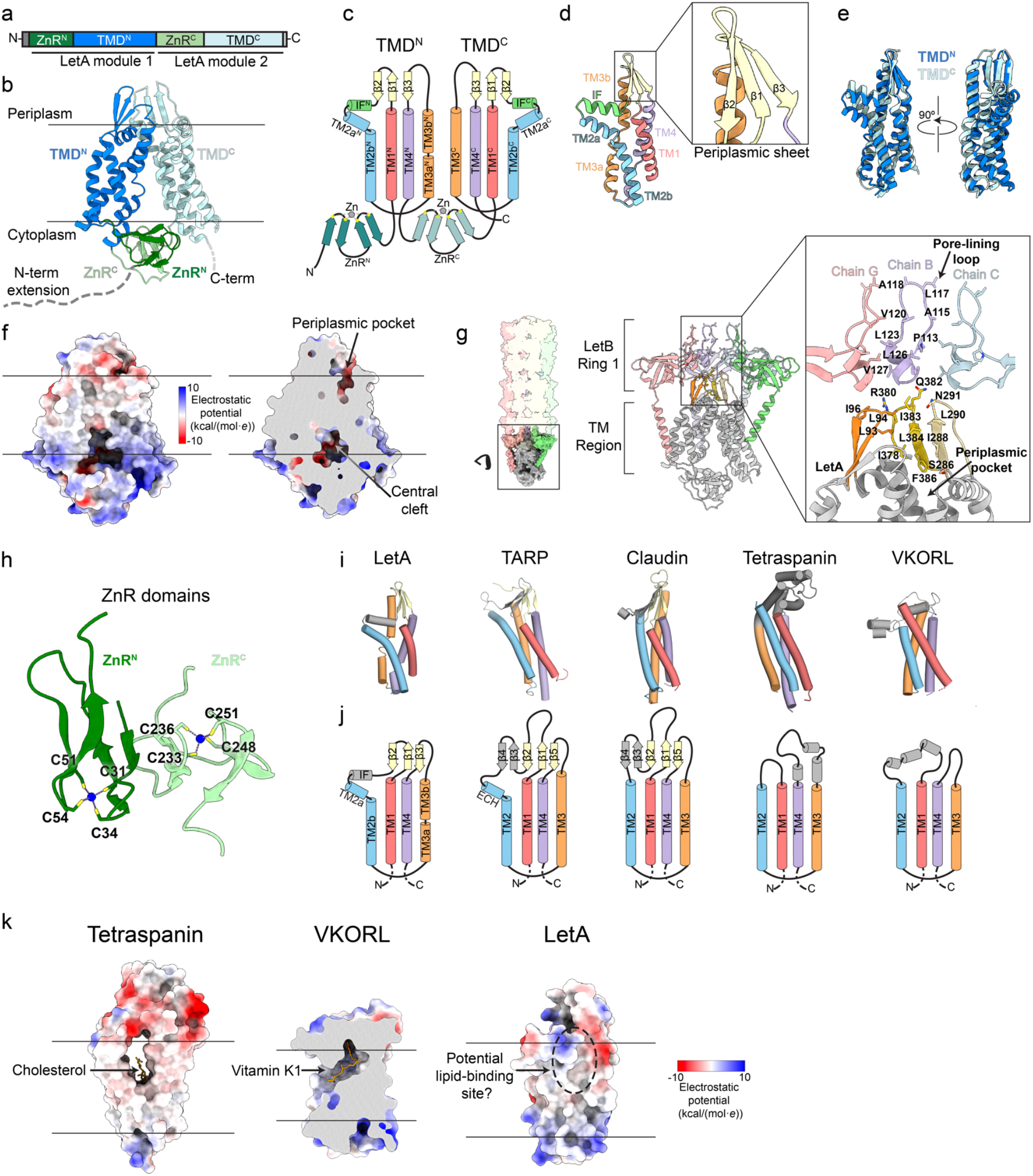
Structural overview of LetA. **a**, Schematic representation of the LetA protein domain organization. **b**, Cartoon representation of the LetA structure, colored as in (**a**). N-terminal and C-terminal extensions were not resolved in our density, and are shown as dashed lines drawn approximately to scale. Membrane boundaries are indicated by black lines. **c**, Topology diagram of LetA. In addition to secondary structure, the Zn-coordinating cysteines (yellow circles) and Zn atoms (gray circles) are shown. **d**, Cartoon representation of the LetA TMD^N^. Colors and labels same as in (**c**); periplasmic β-strands are shown in the inset. **e**, Superposition of TMD^N^ and TMD^C^, showing structural conservation between the two domains. TMD^N^ is rotated ~160°, which results in the superposition of the two domains. **f**, Electrostatic potential surface of LetA shown in full (left) and in cross-section (right), highlighting the periplasmic pocket and central cleft. **g**, Surface representation of LetAB, highlighting LetA (gray) and MCE Ring 1 (colored by chain). Cartoon representation of LetA (gray, with the periplasmic sheet in yellow and orange) and MCE Ring 1 of LetB (colored by chain). Inset highlights the periplasmic sheet of LetA and pore-lining loops of LetB. Residues involved in the putative lipid transport pathway are shown as sticks and labeled. **h**, Cartoon representation of the ZnR domains. Metal-coordinating cysteines are labeled and the Zn atoms are shown as blue spheres. **i**,**j**, Cartoon representations with helices are shown as cylinders (**i**) and corresponding topology diagrams (**j**) of LetA TMD^N^, TARPγ2 (PDB 6DLZ), claudin-4 (PDB 7KP4), tetraspanin^40^ (CD81, PDB 5TCX), and vitamin K epoxide reductase-like^41^ (VKORL, PDB 6WV8). The secondary structural elements of LetA TMD^N^ conserved with structurally related proteins are colored as in (**c**). **k**, Surface representations of tetraspanin, VKORL and LetA TMD^N^, colored by electrostatic potential. Cholesterol and vitamin K1 binding pockets of tetraspanin and VKORL, respectively, are shown, and the corresponding region on LetA is indicated with a dotted circle (periplasmic pocket).

On the cytoplasmic side, ZnR^N^ and ZnR^C^ interact to form a structural unit. Each ZnR domain consists of two stacked β-hairpins (Fig. 3h) and contains a tetracysteine motif involved in metal binding (CXXC-X_n_-CXXC, where n ranges from 11 to 18, Extended Data Feig. 3c). ZnR^C^ forms part of the connection between TMD^N^ and TMD^C^, and interacts non-covalently with ZnR^N^ to form a ZnR dimer on the cytoplasmic side of LetA, perhaps stabilizing the association between the N-terminal and C-terminal halves of LetA. In prokaryotes, members of the ZnR family can bind various transition metals depending on the ZnR domain’s functional role^35^, most commonly zinc or iron. To assess if LetA is a metal-binding protein and to profile its metal-binding specificity, we performed inductively coupled plasma mass spectrometry (ICP-MS) on purified LetA protein, which shows enrichment of zinc atoms specifically (Extended Data Feig. 3f). Calibration using a standard curve (see Methods) suggests that LetA specifically binds ~2 zinc atoms per protein molecule (n=2, range: 1.7 - 2), indicating that both ZnR domains preferentially coordinate zinc under our experimental conditions.

### LetA defines a new family of transporters that is distantly related to the eukaryotic tetraspanin superfamily

LetA is thought to be a transporter, but is not related to known protein families at the sequence level. To further assess its evolutionary relationship to known transporter families, we performed a structure-based search of the PDB using FoldSeek^36^. We were unable to identify structural similarity to known transporter folds, suggesting that LetA represents a new type of membrane transport protein. However, this search revealed that an individual LetA TMD is structurally related to the tetraspanin superfamily of integral membrane proteins in eukaryotes. The LetA TMD most closely resembles TARPs and claudins, and is more distantly related to vitamin K epoxide reductase and tetraspanin itself (Figs. 3i-j). All these proteins share a common topology in the TM helices, but only LetA contains ZnR domains, and is arranged as a pseudodimer, containing two consecutive tetraspanin-like domains. Functionally, the eukaryotic proteins are highly divergent, and none are known to exhibit transporter activity. Tetraspanins are involved in membrane organization via the formation of microdomains that serve to recruit binding partners, often involved in signal transduction^37^. TARPs play a role in regulation of ion channel function in neurons^38^, claudins function in cell-cell adhesion^39^ and VKOR is involved in the recycling of oxidized vitamin K1. Interestingly, both tetraspanin and vitamin K epoxide reductase contain lipid-binding sites, for cholesterol^40^ and vitamin K1^41^, roughly in regions corresponding to the periplasmic pocket in LetA, which is a possible substrate binding site (Fig. 3k). LetA is most closely related to TARPs and claudins, which have structurally-equivalent β-sheets in their extracytoplasmic regions with 3-5 β-strands.

To explore evolutionary relationships between LetA and proteins whose structures have yet to be experimentally characterized, we carried out a Foldseek search of the AlphaFold database of predicted protein structures. This search revealed potential uncharacterized structural homologs of full-length LetA that are present in some parasites and marine protists (Extended Data Feig. 3g). The AlphaFold predictions resemble LetA, but lack ZnR domains. MCE proteins are generally restricted to double-membraned bacteria and photosynthetic eukaryotes, raising the question of if and how LetA-like proteins may mediate lipid transport in parasites and marine protists. However, LetA-like proteins identified in kinetoplastids and dinoflagellates appear to be fused to an extracytoplasmic β-jellyroll domain (Extended Data Feig. 3g) reminiscent of the bridge-like lipid transport domains of VPS13^42^, YhdP^43^, and the LPS exporter^44^. Thus, these distantly related relatives of LetA may potentially mediate the transport of lipids or other hydrophobic molecules in some eukaryotes via these bridge-like proteins instead of MCE tunnels. Taken together, these analyses place LetA and LetA-like proteins in the tetraspanin superfamily, which was previously thought to be a eukaryotic innovation^37^, but we show to be present in prokaryotes as well.

### Deep mutational scanning reveals functionally important regions in LetA

To gain unbiased insight into functionally important residues in LetA, we used deep mutational scanning (DMS), in which each position in LetA was mutated to all possible amino acids. The impact of each mutation on LetA function in cells was assessed in the presence of LSB or cholate (Extended Data Feig. 4). Heatmaps illustrating the impact of each mutation on LetA fitness show similar patterns with cholate and LSB (Extended Data Feigs. 5a-b). As expected, mutation of the start codon, or introducing a stop codon at most positions resulted in reduced fitness. To identify positions likely to be important for LetA function, we calculated a tolerance score for each residue (See Methods), ranging from 0 to 1, where 0 means no mutations are tolerated and 1 means all mutations are tolerated (Fig. 4a, Extended Data Feigs. 5a-b). The majority of residues tolerate mutations well (~90% have tolerance scores ≥0.7, Extended Data Feig. 4d), including a ~25 residue cytoplasmic extension at the N-terminus of LetA (Extended Data Feigs. 5a-b). LetA constructs truncating this region are expressed and largely rescue growth of the *ΔpqiAB ΔletAB* strain in the presence of cholate or LSB (Extended Data Feigs. 4e-f, Supplementary Feigure 1g). However, a subset of positions in LetA were less tolerant of mutation (TS < 0.7, Extended Data Feig. 4d), including 53 positions for cholate and 37 positions for LSB (Fig. 4a, Extended Data Feigs. 5a-b). The majority of these putative functionally important residues cluster in three discrete regions of the LetA structure: 1) the periplasmic pocket in TMD^C^, 2) a polar network in TMD^C^, and 3) the ZnR domains (Fig. 4b).

**Fig. 4.**
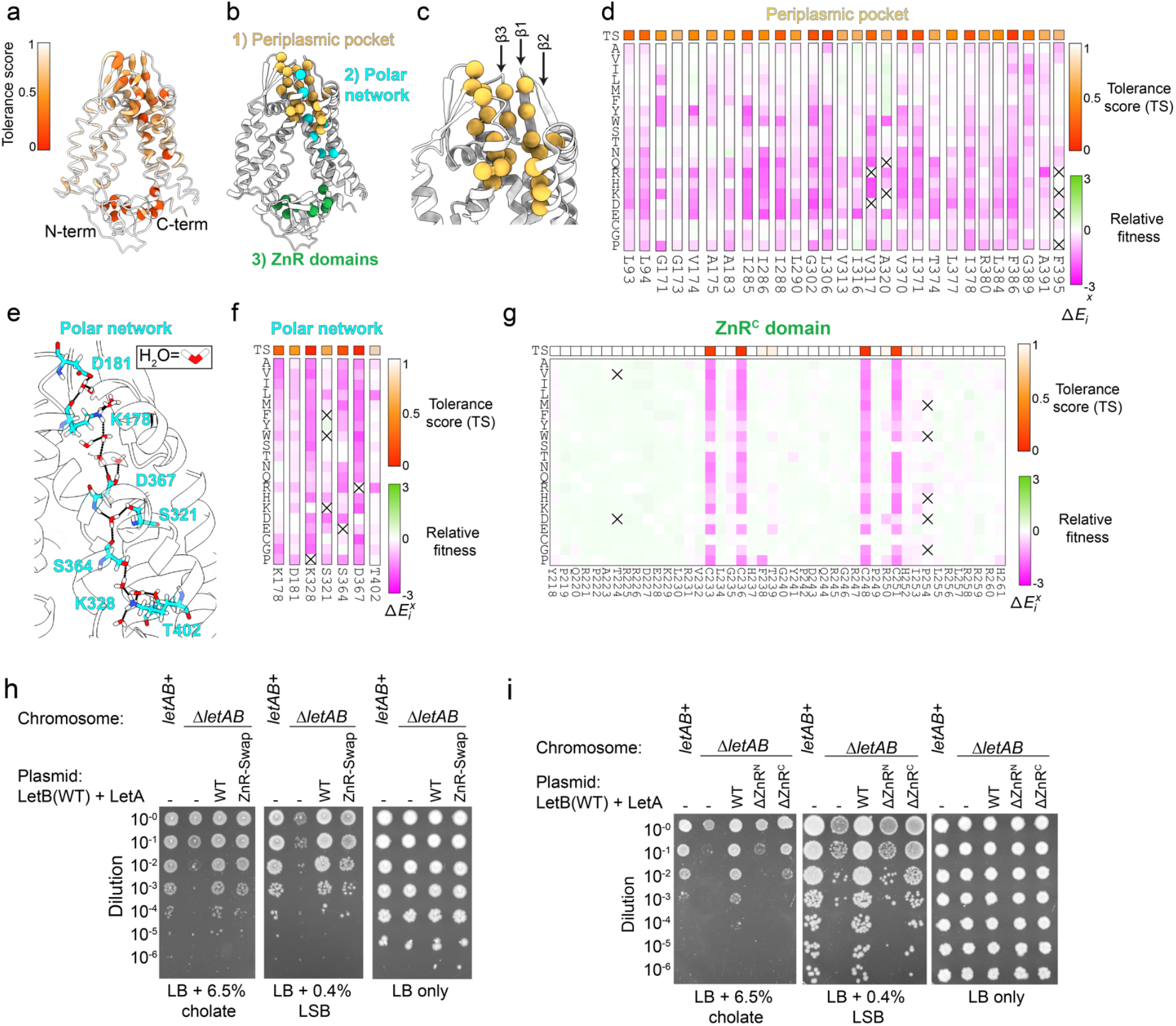
Functional regions of LetA revealed by deep mutational scanning and cellular assays. **a**, LetA structure colored by mutational tolerance scores. Residues most sensitive to mutation appear as the deepest shade of red and thicker backbone trace. **b**, Cartoon representation of LetA displaying residues identified as functionally important by DMS (spheres marking each Cα position). Three functional regions were identified: Periplasmic pocket (yellow), polar network (cyan), and ZnR domains (green). **c**, Enlargement of LetA periplasmic pocket, highlighting the position of the periplasmic β-sheet and functionally important residues. **d**, DMS data corresponding to residues in the periplasmic pocket. Vertical strips for individual LetA residues from the heat map shown in Extended Data Feig. 5a are reproduced here. X-axis: residue positions in WT LetA; Y-axis: all possible amino acid substitutions. Each square represents the average fitness cost of an individual mutation relative to the WT sequence (two replicates). Mutations that decrease fitness relative to the WT are shown in shades of magenta, while mutations that increase fitness are in shades of green. Squares containing an “X” indicate incomplete coverage. The colored square above each strip indicates the tolerance score, calculated as described in the Methods. **e**, A snapshot of the LetA coordinates from equilibrium MD simulations, highlighting the region of the LetA structure corresponding to the polar network. Residues in the polar network with low tolerance scores from DMS experiments are shown as sticks. Water molecules from MD simulations are shown, and hydrogen bonds between the residues and water molecules are illustrated as black dotted lines. **f**,**g**, DMS data corresponding to residues in the polar network **f** and ZnR^C^ domain **g**. Individual strips are shown for each residue in (F) or the whole region corresponding to ZnR^C^ **g**, with colors and annotations as in (**d**). **h**,**i**, Cellular assay to assess the function of LetA mutants, in which the ZnR domains of LetA are replaced with the ZnR domains from *E. coli* PqiA (**h**) or are deleted (**i**). WT LetB is co-expressed with each ZnR mutant. 10-fold serial dilutions of strains were spotted on LB agar with or without the indicated detergent. All strains are constructed in a *ΔpqiAB* background.

#### Periplasmic pocket

The periplasmic pocket (Fig. 4c) is open to the periplasm, and lies just below the pore through MCE Ring 1 of LetB. Thus, this pocket is well positioned to serve as a binding site for lipids moving between the IM and the LetB tunnel, and is analogous to the lipid binding sites observed in tetraspanin and VKOR. From the cholate and LSB datasets combined, approximately half of all residues with low tolerance scores clustered to this region (28 residues), suggesting that the periplasmic pocket is of functional importance (Fig. 4d). Of the 28 residues, 19 have a TS <0.7 in both the cholate and LSB datasets, while the remainder follow similar fitness patterns in LSB and cholate, albeit with slightly higher tolerance scores in the cholate dataset. The majority of these residues are hydrophobic, and are less tolerant of mutations to polar residues, suggesting that maintaining the hydrophobic character of this pocket is critical, consistent with a role in binding to lipids or other hydrophobic molecules. Most of the 28 residues are buried within the pocket or cluster to TMD^C^ strands β1 and β3, which may allow β1 and β3 to act as a hydrophobic “slide” for lipid translocation between the periplasmic pocket of LetA and the pore of MCE Ring 1 (Figs. 3g,4c). Taken together, DMS and the LetA structure support an important role for the periplasmic pocket in LetA function, potentially as a substrate-binding site involved in lipid translocation between LetA and LetB.

#### Polar network in TMD^C^

A cluster of residues with low tolerance scores lies in the TM region of TMD^C^ (Figs. 4e-f), and these residues have polar or charged side chains. The presence of polar and charged side chains in TM regions is unusual, as residues in TM regions are usually hydrophobic. This cluster includes residues K178, D181, S321, K328, S364, D367, and T402 and forms a polar network across the membrane, from the periplasmic pocket to the cytoplasm via the central cleft (Fig. 4e). Two of these residues (S364, D367) belong to the ^361^GRWSM-Ѱ-D-Ѱ-F^369^ motif near the central cleft of LetA, and other residues in the motif are also moderately sensitive to mutation (Extended Data Feigs. 5a-b). Multiple sequence alignment of a panel of LetA-like proteins show that they are well conserved (Extended Data Feig. 3c). Mutating each of the polar network residues to alanine resulted in stable protein complexes capable of binding to LetB, suggesting that these mutations do not drastically disrupt LetA folding or stability, but rather likely have a specific role in the transport mechanism (Extended Data Feig. 5c, Supplementary Feigure 1h). Studies of similar polar networks in other TM proteins have shown that these residues can be important for energy transduction by forming proton transfer pathways^1,45^, often via a combination of ionizable residues and intervening, bridging water molecules. Consistent with a potential role in proton shuttling, several of these residues are predicted to have significantly perturbed pKa’s: K178 and K328 have predicted pKa’s of 8.72 and 8.35, respectively, which are lower than the normal pKa of lysine (≥10) while D367 has a predicted pKa (6.71) that is higher than the normal pKa of aspartate (4.0). To assess whether the residues in the polar network are capable of shuttling protons, we performed equilibrium MD simulations to examine their solvent accessibility. Waters were clearly observed within the polar network, hydrogen bonding with the polar side chains and forming a network of interactions bridging the periplasmic and cytoplasmic spaces (Fig. 4e, Extended Data Feigs. 5d-e). Notably, only the core of TMD^C^ is accessible to water, whereas the core of TMD^N^, which lacks a polar network, remains inaccessible (Extended Data Feig. 5d). Proton shuttling through the polar network could potentially be harnessed to energize conformational changes of LetA for lipid transport.

#### ZnR domains

The cysteines of ZnR domains are expected to be important for function, as they typically maintain the structural fold through tetrahedral coordination of the metal ion^46^. Consistent with this expectation, we found that the metal-coordinating cysteines cannot tolerate mutations (Fig. 4g, Extended Data Feigs. 5a-b). Individual cysteine-to-alanine mutants were indistinguishable from a complete deletion of *letAB* when tested for growth in the presence of cholate or LSB (Extended Data Feig. 5f).

Surprisingly, no other residues in the ZnR domains were sensitive to mutation (TS ≥0.7) (Fig. 4a). This led us to hypothesize that the overall ZnR fold, rather than the sequence, is important for LetA function. To test this, we replaced ZnR^N^ and ZnR^C^ of LetA with those of *E. coli* PqiA, which share 21% and 48% sequence identity, respectively (Extended Data Feig. 5g). The ZnR-swap mutant showed cholate and LSB resistance similar to the WT, suggesting that substantial sequence divergence can be tolerated outside of the Zn coordinating cysteines (Fig. 4h, Extended Data Feig. 5h, Supplementary Feigure 1i). Given the surprisingly high tolerance for sequence divergence, we hypothesized that the low fitness of metal-coordinating cysteines could be explained by the effect of these mutations on protein folding. Western blotting of the Cys mutants shows reduced levels of LetA protein, consistent with the hypothesis that the mutants fold poorly or are less stable (Extended Data Feig. 5i, Supplementary Feigure 1j).

To examine whether the ZnRs are essential for function, we generated two mutants, in which either ZnR^N^ or ZnR^C^ were deleted, and tested these in complementation assays. LetAΔZnR^N^ fails to rescue growth, while LetAΔZnR^C^ partially rescues growth (Fig. 4i) despite reduced expression in cells compared to the WT (Extended Data Feig. 5j, Supplementary Feigure 1k). The LetAΔZnR^C^ mutant binds LetB in a pull down assay, suggesting that the protein is folded and at least partially functional (Extended Data Feig. 5k). Consistent with the idea that the ZnR^C^ domain may not play a key role in LetA function, some LetA homologs lack ZnR^C^ entirely (e.g., *Shewanella sp*. SNU WT4, Uniprot A0A4Y6I6U8). In contrast, the LetAΔZnR^N^ protein is unable to pull down LetB, suggesting that this mutation may interfere with LetA folding (Extended Data Feig. 5k, Supplementary Feigure 1l). Taken together, our results lead to a model in which the ZnR domains likely play a role in modulating the structure or conformational ensemble sampled by LetA, rather than playing a direct role in transport.

### Lipid binding and specificity

In the central cleft in LetA, additional density is apparent in the uncrosslinked EM map, consistent with the size and shape of a diacyl lipid, which we call “Lipid 1” (Fig. 5a, Extended Data Feig. 6a).

**Fig. 5.**
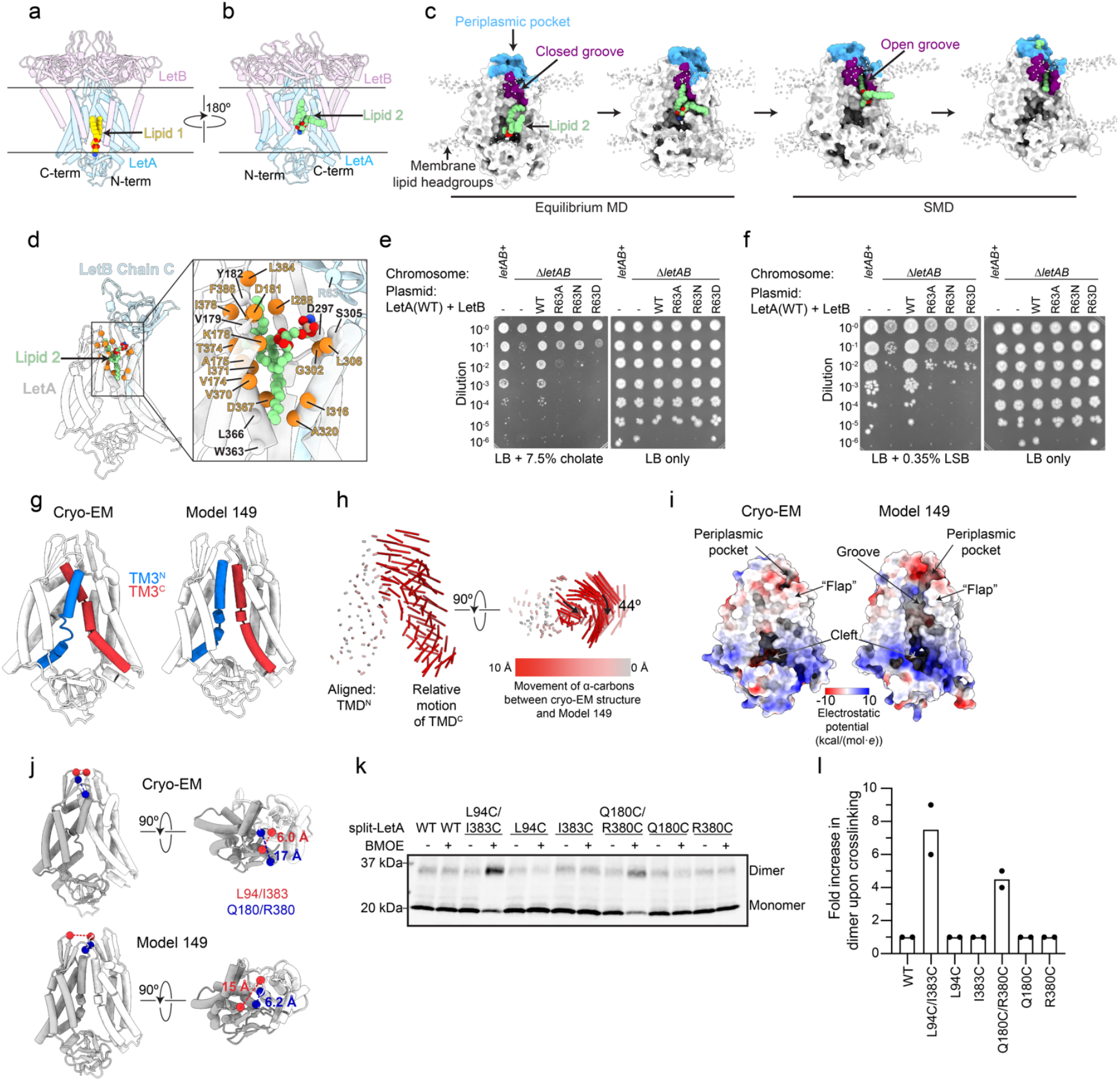
LetA conformational states and interactions with lipids. **a**, Cartoon representation of the uncrosslinked cryo-EM structure of LetA (blue) and MCE Ring 1 (purple) with Lipid 1 (yellow spheres) bound in the central cleft. Density corresponding to Lipid 1 is shown in Extended Data Feig. 6a. **b**, LetA coordinates from an equilibrium MD simulation (Replica 3, −Lipid 1) rotated 180° relative to (**a**) with Lipid 2 (green spheres) shown. **c**, Four snapshots of LetA coordinates from an MD equilibrium simulation (Replica 3, −Lipid 1) followed by an SMD simulation in which the lipid is pulled by one tail. LetA (white surface) is in the same orientation as (**b**), and Lipid 2 (PMPE, green), amphipathic groove (purple), and periplasmic pocket (blue) are indicated. Phosphorus atoms of the bulk membrane lipids are represented as white spheres to indicate the position of the membrane. **d**, Snapshot of LetAB coordinates from the end of a 300-ns equilibrium simulation after SMD simulations. The inset highlights the potential interaction of Lipid 2 (PMPE, carbon: green, nitrogen: blue, oxygen: red, phosphorus: beige) with surrounding residues, shown as spheres marking each Cα position. Interacting residues that are sensitive to mutation are shown as orange spheres, and those that are tolerant to mutations are shown as white spheres. **e**,**f**, Cellular assays to assess the function of LetB mutants. WT LetA is co-expressed with each LetB mutant. 10-fold serial dilutions of strains were spotted on LB agar with or without cholate (**e**) or LSB (**f**). All strains are constructed in a Δ*pqiAB* background. **g**, Cryo-EM structure of LetA and AlphaFold Model 149 with TM3 helices colored to highlight potential changes in conformation of these regions. **h**, Visualization of changes in Cα positions between the cryo-EM structure and AlphaFold Model 149. The model on the left is oriented as in (**g**), but omitted for clarity; vectors indicate the displacement of Cα positions between the cryo-EM structure and AlphaFold Model 149. Models are aligned on TMD^N^, emphasizing the relative motions of TMD^C^. Black arrows indicate direction of domain movement in Model 149 relative to the cryo-EM structure. **i**, Molecular surfaces of the LetA cryo-EM structure and AlphaFold Model 149, colored by electrostatic potential. **j**, Cryo-EM LetA structure and AlphaFold Model 149, with residues L94, I383, Q180, and R380 shown as spheres. The Cβ distances are shown for the pairs of residues that are mutated to cysteine in each case. **k**, Representative Western blot of a crosslinking experiment to probe the LetA conformations observed in the cryo-EM structure and AlphaFold Model 149, as shown in (**j**). Samples were treated with either DMSO (3%, control) or BMOE (1 mM) and subjected to Western blotting using α-LetA antibody (clone 72). Successful crosslinking results in a heterodimeric product with an apparent molecular weight of ~35 kDa. **l**, Quantification of Western blots from LetA Cys crosslinking assay, including the blot from (**k**) as well as one additional biological replicate (n=2 total). Bar graph shows the fold increase in the crosslinked heterodimeric product upon treatment with the BMOE crosslinker relative to the corresponding uncrosslinked control.

Site-specific crosslinking suggests that the Lipid 1 site is also occupied by a phospholipid *in vivo* (Extended Data Feigs. 6a-b, Supplementary Feigure 1m), and MD simulations show that a lipid remains stably bound at this site, whether or not it is included in the starting model (Extended Data Feigs. 6c-e). LetA residues contacting this lipid, however, are generally insensitive to mutation, as seen in our DMS data (Extended Data Feigs. 5a-b, 6a). Collectively, these findings suggest that Lipid 1 may bind stably to LetA, but may not represent a substrate.

Intriguingly, in our MD simulations, we observed spontaneous upward movement of a lipid into the central cleft, which is on the side of LetA opposite the Lipid 1 site (Fig. 5b). The identity of this lipid, which we refer to as “Lipid 2”, is different among replicas, and its position varies slightly between replicas (see Methods, Extended Data Feig. 6f). Lipid 2 originates from the cytoplasmic leaflet of the IM, and moves along the cleft to a position approximately halfway across the membrane towards the periplasmic pocket. The lipid then remains stably bound at this position (Fig. 5c, Extended Data Feig. 6f, Supplementary Video 1), without any further movement towards the periplasm. We hypothesize that Lipid 2 may represent a substrate and suggest a transport mechanism, involving 1) spontaneous movement of a lipid from the cytoplasmic leaflet to the site observed near the middle of the membrane, 2) translocation to the periplasmic pocket, perhaps following a conformational change in LetA, and 3) transfer from the periplasmic pocket of LetA to the LetB tunnel. To explore the possible trajectory of Lipid 2 between its stable position in the LetA central cleft to the periplasmic pocket, we performed steered MD (SMD) simulations. We found that to accommodate Lipid 2, the IF^C^ and TM2a^C^ helices open laterally as a unit, analogous to a flap, revealing an amphipathic groove (Fig. 5c, Supplementary Movie 2). Pulling on one tail required the least amount of work (Extended Data Feig. 6g) compared to pulling both tails or the head group, and resulted in the tails being splayed apart, shielded from the solvent by hydrophobic residues, while the headgroup is inside the pocket interacting with polar residues (Figs. 5c-d, Supplementary Movie 2). The MD simulations suggest a plausible pathway for Lipid 2 movement from the inner leaflet to the periplasmic pocket, facilitated by conformational changes.

To observe if and how Lipid 2 relaxes into the periplasmic pocket after translocation from the Lipid 2 binding site in the central cleft, we conducted 300-ns equilibrium simulations starting from the final state reached during each SMD simulation (Extended Data Feig. 6c). We found that Lipid 2 is highly flexible in the periplasmic pocket, and samples an ensemble of configurations (Extended Data Feigs. 6h-i). However, we observed two common themes. First, the negatively charged phosphate headgroup of the lipid tended to dock against R63 from LetB, which is on the bottom of MCE Ring 1, and faces towards the periplasmic pocket of LetA (Fig. 5d, Supplementary Movie 3). LetB R63 is highly conserved across LetB-like proteins, and is important for LetAB function, as mutations result in sensitivity to both cholate and LSB despite similar expression levels (Figs. 5e-f, Extended Data Feigs. 6j-k, Supplementary Feigure 1n). Second, the fatty acyl tails sample the surrounding hydrophobic surfaces of LetA in different orientations, including with both tails oriented downwards into the periplasmic pocket, upwards towards the pore of LetB, or in a splayed conformation (Fig. 5c, Extended Data Feig. 6i, Supplementary Movie 3). The interactions between Lipid 2 and LetAB are reminiscent of the protein-lipid interactions described in the cryo-EM structure of *E. coli* MlaFEDB^16^. While MlaFEDB is energized by an ABC transporter, MlaFE, which is unrelated to LetA, both transporter systems interact with an MCE protein: MlaD or LetB. In MlaFEDB, one of the bound phospholipids adopts a splayed conformation, with the headgroup docked against a nearby Arg residue, analogous to LetB R63. This binding mode was proposed to facilitate lipid reorientation and transfer between the MCE subunit and the ABC transporter. A similar process may occur during lipid transfer between LetA and LetB, consistent with our hypothesis that the periplasmic β-strands of LetA provide a pathway for lipid movement from LetA to LetB (Figs. 3g, 4c), and the splayed conformation may maximize lipid contact with hydrophobic surfaces during the transfer.

Our MD simulations do not speak to lipid specificity for the substrate. All lipids used in the simulations (PE, PG and cardiolipin) are observed to occupy the Lipid 2 position in the central cleft at least once in the different simulations. To further explore the lipid binding specificity of LetAB, we performed lipidomics experiments on purified LetAB. We found that PE was enriched and cardiolipin was depleted in purified LetAB relative to the *E. coli* membrane as a whole, while PG levels were not significantly different (Extended Data Feigs. 6l-m). Whether this preference reflects LetA substrate specificity, or membrane lipids bound in the periphery of the TMDs remains unclear. Taken together, our data suggest the presence of two lipids, Lipid 1, which likely remains stably bound in the LetA cleft, and Lipid 2, which likely represents the substrate.

### Conformational changes in LetA that may facilitate transport

If LetA is indeed a transporter, it must cycle through a series of conformational states to enable lipid translocation across the bilayer. To explore potential additional states of LetA, we turned to protein structure prediction. We used AlphaFold2^47^ with reduced multiple sequence alignment depth, which was recently pioneered as an approach to predict alternative states of transporters^48,49^. We generated 160 predictions, which can be grouped into five major clusters based upon the state of the LetA TMDs (see Methods, Fig. 5g, Extended Data Feigs. 7a-b, Supplementary Data 1). The models in Cluster 1 are similar to our cryo-EM structure. In the remaining four clusters, we observe two major types of motion, which occur to varying degrees and in different combinations. First, TMD^C^ rotates relative to TMD^N^, up to ~45 degrees, around an axis roughly perpendicular to the membrane membrane plane (Fig. 5h). Second, in Clusters 4 and 5, one TM3 segment slides past the other at the interface between the TMDs (Fig. 5g, Supplementary Movie 4). The combination of TMD rotation and TM3 sliding motions result in the opening of an amphipathic groove between the two TM3 helices on one side and the flap formed by the IF^C^ and TM2a^C^ helices on the other, which connects the Lipid 2 site in the central cleft from our MD simulations to the periplasmic pocket (Fig. 5i). The groove is composed of polar and hydrophobic residues that are sensitive to mutations. This predicted state suggests a possible pathway for lipid movement that is consistent with our SMD simulations.

To assess whether LetA indeed samples alternative states similar to those predicted by AlphaFold2, we focused on Model 149, which represents some of the largest conformational changes in our predicted states relative to our cryo-EM structure. Based on the difference in conformations between our cryo-EM structure and Model 149, we designed an *in vivo* cysteine-based crosslinking assay. We introduced pairs of cysteine mutations at positions in LetA, for which the distance between the cysteine pair is expected to change between the cryo-EM state and Model 149. In the presence of a bi-functional maleimide reagent with a short linker between reactive groups (BMOE; ~8 Å), the mutations we engineered are predicted to be selectively crosslinked in either Model 149 (Q180C/R380C) or the cryo-EM (L94C/I383C) structure, but not in both (Fig. 5j). We used our split-LetA construct with non-essential cysteines removed as the background for these mutations (Cys-light LetA; see Methods), as successful crosslinking is expected to covalently link the two LetA modules together to form a crosslinked dimer, resulting in a large mobility shift when analyzed by SDS-PAGE. The Cys-light LetA, as well as the L94C/I383C and Q180C/R380C derivatives, are all functional in cells (Extended Data Feig. 7c).

BMOE treatment of the L94C/I383C and Q180C/R380C mutants led to a ~7-and ~5-fold, respectively, increase in the dimer band relative to that of the DMSO control (Figs. 5k-l, Supplementary Feigure 1o), suggesting that the cryo-EM and Model 149 states are indeed sampled in cells. The L94C, I383C, Q180C, and R380C single mutant controls exhibit background levels of crosslinking, suggesting that the crosslinking in the double mutants is specific. Together, these results support the existence of an “open-groove” LetA state that is consistent with AlphaFold2 predictions, which may play a role in facilitating transport.

## Discussion

LetB forms a tunnel across the bacterial cell envelope, creating a hydrophobic tunnel for lipid transport between the IM and OM. How lipids are loaded into LetB was unknown, and LetA was hypothesized to play a role in the lipid transport mechanism. We propose a model for LetAB-mediated lipid transport (Fig. 6). 1) In the absence of LetA, LetB MCE Ring 1 is predominantly in the closed conformation, preventing the passage of lipids through the tunnel. LetA binding to LetB results in the opening of the tunnel. In the cell, it is possible that LetA is constitutively bound to LetB or it may be that LetA binding regulates the opening of the LetB tunnel. 2) LetA may cause local distortion of the membrane, enabling the spontaneous movement of phospholipids from the inner leaflet of the IM into the LetA central cleft at the middle of the membrane. 3) To translocate the lipid, LetA may undergo a conformational change, likely driven by proton shuttling, revealing an amphipathic groove. The conformational change in LetA was predicted by AlphaFold2, and is consistent with the results of *in vivo* crosslinking experiments. 4) The lipid traverses the amphipathic groove with the tails in a splayed configuration, in which each fatty acyl tail interacts with functionally important hydrophobic and polar residues. Residues involved in proton-shuttling are also predicted to interact with the lipid headgroup during the translocation process, as has been described for proton-coupled transporters^1,50^. 5) The lipid arrives in the LetA periplasmic pocket, where it is largely flexible, but constrained by interactions of the polar head group with R63 of LetB MCE Ring 1. The lipid tails interact with surrounding hydrophobic residues from LetA, and ultimately the lipid slides across the periplasmic β1 and β3 strands of LetA to enter the LetB tunnel. 6) The lipid is extruded from the periplasmic pocket into the LetB tunnel, and LetA returns to the resting state. This mechanism may be similar to extractors such as the LPS exporter, which extracts LPS from the outer leaflet of the IM via the central cavity of the LptFG dimer^51^. ATP-dependent conformational changes result in collapse of the central pocket, and extrusion of LPS into a periplasm-spanning bridge formed by LptC proteins, to mediate LPS transport to the OM. A similar open-to-collapsed transition is observed in tetraspanin, where cholesterol and other lipids have been shown to bind the central cavity and modulate protein conformation^40^ (Extended Data Feig. 7d). An attractive but speculative possibility is that an ancestral lipid binding protein reminiscent of tetraspanin, already undergoing lipid-dependent conformational changes, evolved to couple the proton motive force to lipid binding and unbinding, leading to proton-driven, active transporters such as LetA.

**Fig. 6.**
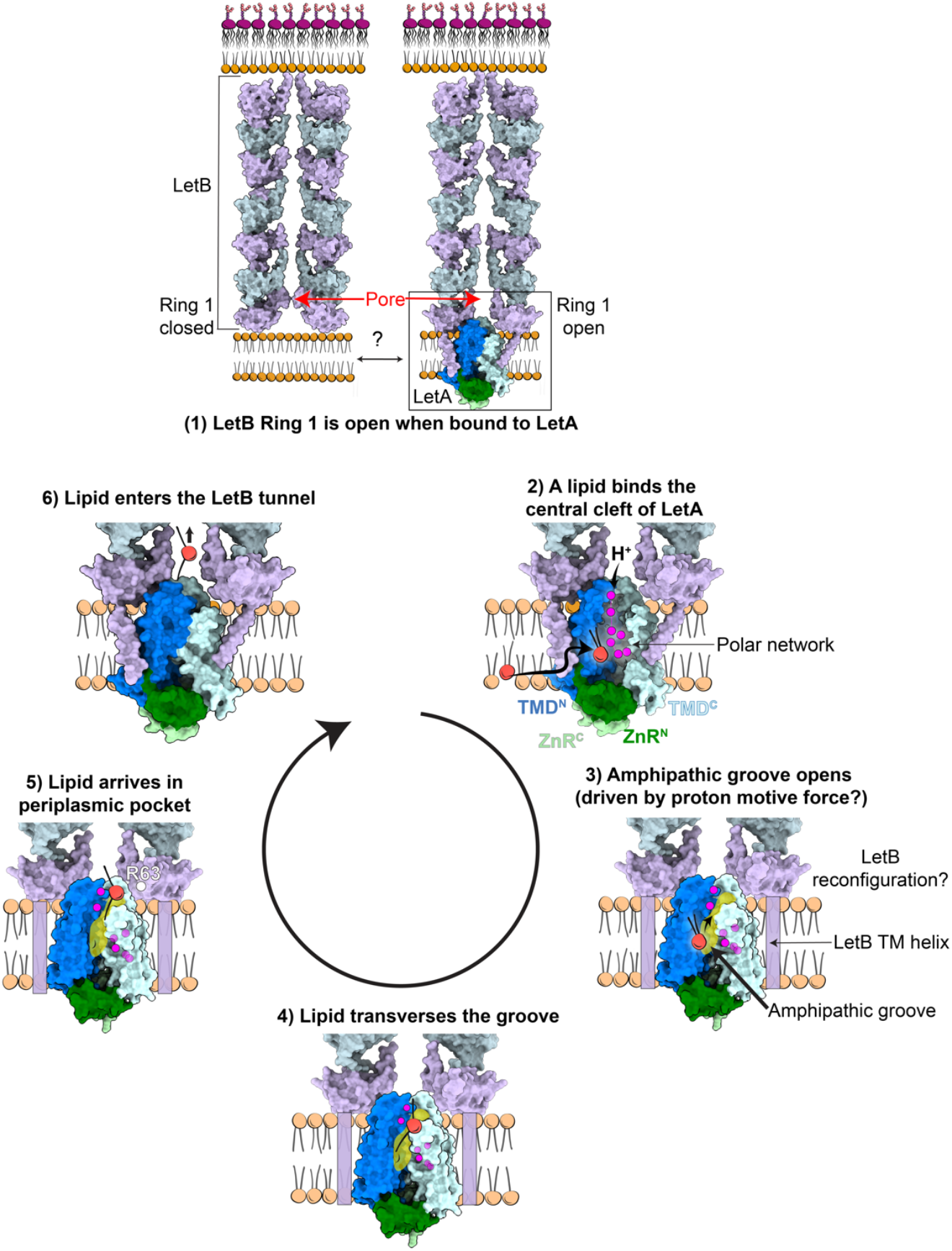
Model for lipid translocation by LetAB. Proposed model for lipid translocation and LetA conformational changes. (1) MCE Ring 1 is open when bound to LetA. Closed (left, PDB 6V0C) and open LetB states (right) are shown. (2) A phospholipid binds to the central cleft of LetA, representing Lipid 2 from MD simulations. (3) LetA adopts an alternative state, possibly driven by the proton motive force, revealing an amphipathic groove. The alternative state may resemble Alphafold Model 149. This conformation results in a steric clash with LetB, making it likely that some conformational rearrangements simultaneously occur in LetB. (4) Lipid transverses the groove, as observed in SMD simulations and (5) arrives in the periplasmic pocket, where R63 of LetB dictates lipid orientation. (6) LetA likely reverts to the cryo-EM state, driving the phospholipid into the LetB tunnel.

## Supporting information

Composite model related to Map 1

Composite model related to Map 2

## Acknowledgements

We thank James Chen, Margot Di Cesare, Yongbo Ding, Sabrina Giacometti, Alice Herneisen, Kacie McCarty, and Edward Twomey for critical reading and feedback on our manuscript. We gratefully acknowledge the following funding sources: NIH R35GM128777, NIH 1K99GM157496-01, NIH R24-GM145965, NIH R01-DK128315, Charles H. Revson Foundation, and the Burroughs Wellcome Fund. We thank Yang Zhang for generating the LetA plasmid (pBEL2214) and purifying LetA protein. Cryo-EM grids were prepared and screened at NYU Langone Health’s Cryo-Electron Microscopy Laboratory, which is partly supported by grants NIH/NCI P30CA016087 and R01NS108151-07, and we thank Alice Paquette and William Rice for assistance with cryo-EM grid screening and microscope operation. A portion of this research was supported by NIH grant R24GM154185, as cryo-EM data collection was performed at the Pacific Northwest Center for Cryo-EM (PNCC) with assistance from Harry Scott and Rose Marie Haynes. Cryo-EM data collection was also performed at the National Center for CryoEM Access and Training (NCCAT) and the Simons Electron Microscopy Center located at the New York Structural Biology Center, supported by the NIH Common Fund Transformative High Resolution Cryo-Electron Microscopy program (U24 GM129539, and NIGMS R24 GM154192) and by grants from the Simons Foundation (SF349247) and NY State Assembly. For data processing, we used computing resources maintained by the High Performance Computing (HPC) Facility at NYU and the Advanced Research Computing at Hopkins (ARCH) core facility (rockfish.jhu.edu), which is supported by the National Science Foundation (NSF) grant number OAC1920103. ICP-MS data were obtained by Stephen J. Eyles at the University of Massachusetts Mass Spectrometry Core Facility, RRID:SCR_019063. Illumina sequencing was performed by NYU Langone’s Genome Technology Center (RRID: SCR_017929). This shared resource is partially supported by the Cancer Center Support Grant P30CA016087 at the Laura and Isaac Perlmutter Cancer Center. The MD simulations have been performed using the computational resources provided by the NSF Supercomputing Centers (ACCESS grant number MCA06N060), and Delta advanced computing and data resource which is supported by NSF (award OAC 2005572) and the State of Illinois. Monoclonal antibody generation was supported in part by the CSHL Antibody and Phage Display Shared Resource funded by the Cancer Center Support Grant 5P30CA045508.

## Data Availability

The cryo-EM maps have been deposited at the Electron Microscopy Data Bank under the following accession codes: Map 1 (EMD-49148), Map 1a (EMD-49145), Map 1b (EMD-49146), Map 1c (EMD-49147), Map 2 (EMD-49152), Map 2a (EMD-49149), Map 2b (EMD-49150), Map 2c (EMD-49151). The coordinates of the atomic models have been deposited at the PDB under the following accession codes: 9N8W (Map 1), 9N8X (Map 2). Raw sequencing reads were deposited to the Sequence Read Archive under BioProject ID: PRJNA1221345 (accessions SAMN46739294-301). To analyze reads resulting from deep mutational scanning, scripts from the following Github repository were used, https://github.com/MaxabHaase/LetA. Processed variant counts and fitness scores were deposited to MaveDB under the experiment X. Files related to molecular dynamic simulations have been deposited at Zenodo under dataset identifier X. The lipidomics mass spectrometry files are available at MassIVE under dataset identifier MSV000096297. A list of plasmids that have been deposited at Addgene is provided in Supplementary Table 2 with their identifiers. Uncropped images from Western blots and Coomassie gels shown are provided in Supplementary Feigure 1.

## Methods

### Expression and purification of LetAB

Plasmid pBEL1284, which encodes for N-terminal 6xHis2xQH-TEV tagged LetA and untagged LetB, was transformed into OverExpress C43 (DE3) cells (Lucigen, 60446-1) (Supplementary Table 2). For protein expression, overnight cultures (LB + 100 µg/mL carbenicillin + 1% glucose) were diluted in LB (Difco) supplemented with carbenicillin (100 µg/mL), grown at 37°C with shaking to an OD600 of ~0.9, and then induced by addition of arabinose to a final concentration of 0.2%.

Cultures were further incubated at 37°C with shaking for 4 hours, and then harvested by centrifugation. The pellets were resuspended in lysis buffer (50 mM Tris pH 8.0, 300 mM NaCl, 10% glycerol), flash frozen in liquid nitrogen and stored at −80°C. Cells were lysed by two passes through an Emulsiflex-C3 cell disruptor (Avestin), then centrifuged at 15,000g for 30 minutes at 4°C to pellet cell debris. The clarified lysate was subjected to ultracentrifugation at 37,000 rpm (182,460 g) for 45 minutes at 4°C in a Fiberlite F37L-8 × 100 Fixed-Angle Rotor (Thermo Fisher Scientific, 096-087056). The supernatant was discarded and the membrane fraction was solubilized in 50 mM Tris pH 8.0, 300 mM NaCl, 10% glycerol, 25 mM n-Dodecyl-B-D-maltoside (DDM) by rocking overnight at 4°C. Insoluble debris were pelleted by ultracentrifugation at 37,000 rpm for 45 minutes at 4°C. Solubilized membranes were then passed twice through a column containing Ni Sepharose Excel resin (Cytiva). Eluted proteins were concentrated using an Amicon Ultra-0.5 Centrifugal Filter Unit concentrator (MWCO 100 kDa, UFC510096) before separation on the Superdex 200 Increase 10/300 column (Cytiva) equilibrated with either Tris (20 mM Tris-HCl pH 8.0, 150 mM NaCl, 0.5 mM DDM and 10% glycerol) or HEPES (20 mM HEPES pH 7.4, 150 mM NaCl, 0.5 mM DDM and 10% glycerol) gel filtration buffer. Fractions containing LetAB were pooled, concentrated and applied to grids for negative stain EM or cryo-EM. One liter of culture typically yields 30-40 µg of the LetAB complex.

### Negative stain EM

To prepare grids for negative-stain EM analysis, a fresh sample of LetAB was applied to a carbon-coated 400 mesh copper grid (Ted Pella, 01754-F), freshly glow discharged for 30 seconds. and blotted off. Immediately after blotting, 3 µL of a 2% uranyl formate solution was applied for staining and blotted off on filter paper (Whatman 1) from the opposite side. Application and blotting of stain was repeated four times. The sample was allowed to air dry before imaging. A negative stain grid of LetB^ΔTM^ was prepared previously^31^ using a similar procedure and stored. New images from this sample were acquired for this study. Data were collected on the Talos L120C TEM (FEI) equipped with the 4k x 4k OneView camera (Gatan) at a nominal magnification of 73,000x corresponding to a pixel size of 2.03 Å /px on the sample and a defocus range of −1 to −2 μm defocus. Negative-stain dataset size was determined to be sufficient by the ability to see features in the 2D classes of picked particles. For both LetAB and LetB^ΔTM^ datasets, micrographs were imported into cryoSPARC^57^ (v.3.3.1) and approximately 200 particles were picked manually, followed by automated template-based picking. Particles were extracted with a 320 pixel box size. Several rounds of 2D classification were performed using default parameters, except that “Force max over poses/shifts” and “Do CTF correction” were both set to False.

### Cryo-EM sample preparation and data collection

To generate the crosslinked LetAB sample, 1% glutaraldehyde was added to purified LetAB (HEPES gel filtration buffer) at a final concentration of 0.025%. The sample was incubated on ice for one hour and then quenched by the addition of 75 mM Tris-HCl pH 8.0. The sample was incubated for 15 minutes on ice before filtering using an Ultrafree centrifugal filter (catalog #UFC30GVNB) and loading onto a Superdex 200 Increase 10/300 column (Cytiva), equilibrated with buffer containing 20 mM Tris-HCl pH 8.0, 150 mM NaCl, 0.5 mM DDM, to remove aggregated LetAB. Fractions containing the LetAB complex were concentrated to 1 mg/mL using the Amicon Ultra-0.5 centrifugal filter unit concentrator (MWCO 100 kDa, UFC510096). Continuous carbon grids (Quantifoil R 2/2 on Cu 300 mesh grids + 2 nm Carbon, Quantifoil Micro Tools, C2-C16nCu30-01) were glow-discharged for 5 s in an easiGlow Glow Discharge Cleaning System (Ted Pella). 3 μL of freshly prepared sample was added to the glow-discharged grid. Grids were prepared using a Vitrobot Mark IV (Thermo Fisher Scientific). Grids were blotted with a blot force of 0 for 3 s at 4°C with 100% chamber humidity and then plunge-frozen into liquid ethane. Grids were clipped for data acquisition.

Grids containing crosslinked LetAB were screened at the NYU cryo-EM core facility on the Talos Arctica (Thermo Fisher Scientific) equipped with a K3 camera (Gatan). The grids were selected for data collection on the basis of ice quality and particle distribution. The selected cryo-EM grid was imaged on two separate sessions at the Pacific Northwest Center for Cryo-EM (PNCC) on Krios-1, a Titan Krios G3 electron microscope (Thermo Fisher Scientific) equipped with a K3 direct electron detector with a BioContinuum energy filter (Gatan). Super-resolution movies were collected at 300 kV using SerialEM^58^ at a nominal magnification of 81,000x, corresponding to a super-resolution pixel size of 0.5144 Å (or a nominal pixel size of 1.029 Å after binning by 2). 12,029 movies were collected over a defocus range of −0.8 to −2.1 µm and each movie consisted of 50 frames with a total dose of 50 e-/Å^2^. Further data collection parameters are shown in Supplementary Table 1.

The uncrosslinked LetAB complex was prepared as described in “Expression and purification of LetAB”, except the Superdex 200 Increase 10/300 column was equilibrated in buffer containing 20 mM Tris-HCl pH 8.0, 150 mM NaCl, 0.5 mM DDM. Continuous carbon grids (Quantifoil R 2/2 on Cu 300 mesh grids + 2 nm Carbon, Quantifoil Micro Tools, C2-C16nCu30-01) were glow-discharged for 5 s in an easiGlow Glow Discharge Cleaning System (Ted Pella). 3 μL of freshly prepared sample (1 mg/mL) in Tris gel filtration buffer was added to the glow-discharged grid. Grids were prepared using a Vitrobot Mark IV (Thermo Fisher Scientific). Grids were blotted with a blot force of 0 for 3 s at 4°C with 100% chamber humidity and then plunge-frozen into liquid ethane. Grids were clipped for data acquisition. Grids were screened at the NYU cryo-EM laboratory on the Talos Arctica (Thermo Fisher Scientific) system equipped with a K3 camera (Gatan). The grid with the best ice quality and particle distribution was imaged at the New York Structural Biology Center (NYSBC) on Krios-1, a Titan Krios G3 electron microscope (Thermo Fisher Scientific) equipped with K3 direct electron detector with a BioContinuum energy filter (Gatan). Super-resolution movies were collected at 300 kV using Leginon^59^ at a nominal magnification of 81,000x, corresponding to a super-resolution pixel size of 0.535 Å (or a nominal pixel size of 1.083 Å after binning by 2). Movies were collected over a defocus range of −2 to −5 µm and each movie consisted of 40 frames with a total dose of 51 e-/Å^2^. A total of 12,464 movies were collected, consisting of 5,372 movies at 0° tilt and 7,083 movies at −30° tilt. Further data collection parameters are shown in Supplementary Table 1.

### Cryo-EM data processing and model building for crosslinked LetAB

Data processing workflow for the crosslinked LetAB sample is shown in Supplementary Table 1. A combination of cryoSPARC (versions 3.2.0-4.3.0) and RELION^60^ (version 3.1.0) were used for data processing. Dose-fractionated movies were gain-normalized, drift-corrected, summed and dose-weighted, and binned by 2 using the cryoSPARC Patch Motion module. CTF estimation for each summed image was carried out using cryoSPARC Patch CTF estimation. To generate 2D templates for auto-picking, 1003 particles were manually picked, extracted (box size = 576 px) and subjected to 2D classification. The classes with top, tilted and side views of LetAB were selected as templates for auto-picking, which yielded 3,582,925 particles after extraction (box size = 576 px). The particles were subjected to 2D classification (200 classes) with Force Max over poses/shifts set to False. Well-aligned 2D classes were selected (1,793,362 particles) and a 3D reconstruction was generated using ab-initio reconstruction. The 3D reconstruction was used as a template for 3D refinement in RELION, which revealed well-resolved density for LetB Rings 1-4, poor and noisy density for LetB Rings 5-7, and no density for LetA, likely due to Rings 1-4 dominating the particle alignment. To improve resolution for LetA, local refinement was performed using a mask around LetB Rings 1-4 and the TM region, followed by particle subtraction in RELION where the signal for the TM region and Rings 1-2 were kept and re-centered to the middle of the box (256 px). The subtracted particles were imported into cryoSPARC, where reference-free 3D classification was performed using the ab-initio module (2 classes) to remove misaligned and ‘junk’ particles. This resulted in one class with 1,171,725 particles with high-resolution features. The particles from this class were further cleaned using 2D classification and then subjected to non-uniform refinement (942,263 particles). The aligned particles were imported into RELION and sorted using 3D classification without alignment (8 classes), which revealed one class containing density for the TM region with high resolution features. The particles (158,666) were then subjected to local refinement in RELION, yielding a map (Map 1a) with a nominal resolution of 3.4 Å (Extended Data Feig. 2c).

To obtain high resolution maps of Rings 2-4 and Rings 5-7, the 1,793,362 particles from the initial 2D classification step were Fourier cropped to a box size of 128 pixels. The particles were sorted using heterogeneous refinement (5 classes) in cryoSPARC, which revealed only one class where LetB is straight rather than curved. The particles from this class (448,403) were re-extracted (box size = 512 px), aligned using non-uniform refinement, and imported into RELION. The aligned particles underwent local refinement, followed by particle subtraction to yield signal for either Rings 2-4 or Rings 5-7. During particle subtraction, the subtracted images were re-centered to the middle of the box, which was cropped to either 360 px (Rings 2-4) or 256 px (Rings 5-7). The two particle subtracted stacks were imported into cryoSPARC, where the particles were subjected to reference-free 3D classification using ab-initio reconstruction (3 classes) to remove misaligned and ‘junk’ particles. The particles from the selected class were aligned using non-uniform refinement, and then imported into RELION for 3D classification without alignment (8 classes). The classes with the highest resolution features were selected, their particles combined before being imported into cryoSPARC for non-uniform refinement to improve the densities for both the MCE core domains and the pore-lining loops^31^, resulting in Map 1b (Rings 2-4) and Map 1c (Rings 5-7). All refinement steps were performed without symmetry applied (C1).

During the model building process, initial reports describing AlphaFold2^47^ and RoseTTAFold^61^ became public. To accelerate model building for LetAB, RoseTTAFold^61^ was used to predict the 3D structure of LetA. The model was fit as a rigid body into the LetA density in Map 1a, followed by rigid body fitting of TMD^N^ (aa 66-218), TMD^C^ (aa 261-418), ZnR^N^ (aa 24-65) and ZnR^C^ (aa 219-261). Residues 1-26 and 419-427 were deleted due to the absence of density for them. As our ICP-MS data suggests LetA binds zinc, we used Coot (v.0.8.9.2)^62^ to add zinc ligands to the densities found in between the predicted metal-coordinating cysteines. To build the model for LetB Rings 1-2, PDB 6V0J was used as a starting model, as it best matches the density in Map 1a. The model was first fit as a rigid body into the density corresponding to LetB, followed by rigid-body fitting of each MCE domain. The pore-lining loops of MCE Ring 1 exhibit C3 symmetry. Densities for four out of six TM helices of LetB were observed, and those helices were manually built using Coot. Residues (25-45) of a LetB TM helix (Chain B) were stubbed due to the lack of side chain density.

For Maps 1b and 1c, PDB 6V0F and PDB 6V0E, respectively, best fit into the density, respectively, after rigid body docking. Each MCE domain was rigid-body fit into the density. The pore-lining loops of Rings 5 and 6 exhibit C3 symmetry. Extra densities that do not correspond to the protein are present near the pore-lining loops between Rings 5-6 and Rings 6-7, but the resolution is too low to determine the identity of the ligand. Therefore, ligands were not modeled into these densities. It is possible that these densities represent non-native ligands, such as DDM from the sample buffer. Each model was real-space-refined into its respective map using PHENIX^63^ with global minimization, Ramachandran, secondary structure and ligand restraints. Using UCSF Chimera^64^, Maps 1a and 1c were fit into Map 1b and resampled such that the maps overlaid with one another. These maps were then used to stitch together the models. The MCE Ring 2 model from Map 1b (instead of the one from Map 1a) was used to generate the composite model since this map had complete density for MCE Ring 2. The resulting composite model was used as a template to generate a composite density map (Map 1) using the PHENIX Combine Focused Maps module. The model was real-space-refined into Map 1 using PHENIX with global minimization, Ramachandran, secondary structure and ligand restraints. The composite model reveals that LetB Rings 3, 5 and 6 are in a single, closed conformation, whereas the conformation of Rings 2, 4, and 7 could not be reliably assessed due to weak density for the pore-lining loops (Extended Data Feig. 1g).

For validation, statistics regarding the final models (Supplementary Table 1) were derived from the real_space_refine algorithm of PHENIX and MolProbity^65^, EMRINGER^66^, and CaBLAM^67^ from the PHENIX package^63^ were used for model validation. Model correlations to our EM maps were estimated with CC calculations and map-model FSC plot from the PHENIX package.

### Cryo-EM data processing and model building for uncrosslinked LetAB

Data processing workflow for the uncrosslinked LetAB sample is shown in Extended Data Feig. 2. A combination of cryoSPARC (versions 3.3.1-4.3.0) and RELION (version 4.0-beta) were used for data processing. Particle picking was performed in RELION on the motion-corrected micrographs generated by NYSBC using MotionCor2^68^. 2D templates were generated for auto-picking on manually picked particles. The particles (3,014,365 at 0° tilt; 3,960,481 at −30° tilt) were imported into cryoSPARC and re-extracted (600 px, Fourier cropped to 100 px) from Patch CTF corrected micrographs generated within CryoSPARC (gain-normalized, drift-corrected, summed, binned 2x and dose-weighted using the cryoSPARC Patch Motion module). The particles underwent several rounds of 2D classification (200 classes) with Force Max over poses/shifts set to False. Well-aligned 2D classes were selected, resulting in 1,582,691 particles at 0° tilt and 1,918,023 particles at 30° tilt. The particles were combined, sorted by 2D classification, and the selected particles were re-extracted (512 px). The particles (2,658,362) were then aligned by non-uniform refinement (C6 symmetry applied), and imported into RELION, where they were subjected to local refinement with C6 symmetry relaxation applied. The signal for LetB Rings 2-7 was removed using the Particle Subtraction module. The subtracted images were re-centered so that the signal for LetA+Rings 1-2 was in the middle of the box, which was cropped to 256 px. The subtracted particles were imported into cryoSPARC and sorted by several rounds of 2D classification (200 classes) to remove misaligned and ‘junk” particles. The particles were further sorted using the ab-initio reconstruction (5 classes, 3 rounds) to yield 1,131,012 “clean” particles, which were then aligned using non-uniform refinement. The particles were imported into RELION for 3D classification without alignment and with a mask around LetA, which revealed a class showing high-resolution features. The particles were imported into CryoSPARC for non-uniform refinement. To continue filtering out low-resolution particles, the particles were sorted by 3D classification without alignment in RELION, followed by non-uniform refinement in cryoSPARC, two additional times. After non-uniform refinement, the particles underwent local refinement in cryoSPARC to yield a map with a nominal resolution of 3.4 Å (Map 2a, Extended Data Feig. 2H).

To obtain high resolution maps of Rings 2-4 and Rings 5-7, the 2,658,362 particles from the initial 2D classification step were sorted using heterogeneous refinement (5 classes), which revealed only one class where LetB is straight rather than curved. The particles from this class (738,470) were aligned using non-uniform refinement with symmetry applied, and imported into RELION. The aligned particles underwent local refinement, followed by particle subtraction to yield signal for either Rings 2-4 or Rings 5-7. During particle subtraction, the subtracted images were re-centered to the middle of the box, which was cropped to either 360 px (Rings 2-4) or 256 px (Rings 5-7). The two particle subtracted stacks were imported into cryoSPARC, where the particles were sorted using ab-initio reconstruction (3 classes) to remove misaligned and ‘junk’ particles. The classes with high-resolution features were selected and the particles were aligned using non-uniform refinement, and then imported into RELION for 3D classification without alignment (8 classes). The classes with the highest resolution features were selected, their particles combined and imported into cryoSPARC for non-uniform refinement to improve the densities for both the MCE core domains and the pore-lining loops^31^, resulting in Map 2b (Rings 2-4) and Map 2c (Rings 5-7).

The crosslinked LetAB model was used to build the model for uncrosslinked LetAB. LetA was rigid-body fit into the LetA density in Map 2a. An additional “wishbone”-shaped density was observed in the central cavity of LetA that is consistent with the size and shape of a diacyl phospholipid. Because the density was insufficient to unambiguously assign the head group structure and fatty acid chain lengths, we modeled this density as 1,2-dipalmitoyl-sn-glycero-3-phosphoethanolamine (PDB PEF, 16:0/16:0), which is an abundant phospholipid species in the *E. coli* IM. Atoms not well accommodated in the observed density were pruned from the ligand. The MCE Ring 1 model from Map 1a was rigid-body docked into Map 2a. The individual MCE domains were then rigid-body fit into the map. The model was real-space-refined into Map 2a map using PHENIX with global minimization, Ramachandran, secondary structure and ligand restraints.

To build a model for Rings 2-4 and Rings 5-7, the models from Map 1b and Map 1c were rigid-body fit into Map 2b and Map 2c, respectively. Each MCE domain was rigid-body fit into the map. LetB residues 614-619 were deleted from the model due to poor density in the map. Similar to the crosslinked LetAB model, the pore-lining loops of Rings 1, 5 and 6 exhibit C3 symmetry. Extra densities are present near the pore-lining loops between Rings 5-6 and Rings 6-7. Since the resolution is too low to determine the identity of the ligand, the extra densities were left unmodeled. Each model was real-space-refined into its respective map using PHENIX with global minimization, Ramachandran, secondary structure and ligand restraints. Using UCSF Chimera, Maps 2a and 2c were fit into Map 2b and resampled such that the maps overlaid with one another. These maps were then used to stitch together the models. The MCE Ring 2 model from Map 2b was used to generate the composite model since this map had complete density for MCE Ring 2. The resulting composite model was used as a template to generate a composite density map (Map 2) using the vop command in ChimeraX. The model was refined using Phenix.real_space_refine into Map 2 with global minimization, Ramachandran, secondary structure and ligand restraints.

For validation, statistics regarding the final models (Supplementary Table 1) were derived from the real_space_refine algorithm of PHENIX and MolProbity^65^, EMRINGER^66^, and CaBLAM^67^ from the PHENIX package^63^ were used for model validation. Model correlations to our EM maps were estimated with CC calculations and map-model FSC plot from the PHENIX package.

We used this model to assess the conformation of the LetB Rings via CHAP^52^. LetB Rings 3, 5 and 6 are in a single, closed conformation, whereas the conformation of Rings 2, 4, and 7 could not be reliably assessed due to weak density for the pore-lining loops (Extended Data Feig. 1G). However, since the different segments of LetB were processed separately (Extended Data Feig. 2I), it is unclear if the conformations of Rings 1, 3, 5, and 6 are correlated.

### Sequence alignment

LetA and PqiA proteins are widespread across Proteobacteria. Using the *E. coli* LetA sequence, we performed a protein BLAST to search for LetA and PqiA proteins across the orders within each of the five classes of Proteobacteria. Only sequences that contained both LetA modules were considered. We then performed a tblastn search using the core nucleotide database and the specified organism to determine if the gene is in an operon with *letB* or *pqiB*, which would indicate if the query sequence is a *letA* or *pqiA* gene, respectively. LetB and PqiB can be identified based on sequence length and AlphaFold2 prediction; LetB has six, seven or eight MCE domains while pqiB has three. Through this method, we identified 20 sequences, where nine are LetA and 11 are PqiA proteins. The LetA sequences are from Gammaproteobacteria, whereas PqiA are from Alpha-and Betaproteobacteria. The sequences were aligned using MUSCLE^53^ and annotated using Jalview (version 2.11.3.3)^69^.

To generate a sequence alignment for LetB proteins, we performed a protein BLAST to search for sequences across the orders of Gammaproteobacteria. We then performed a tblastn search using the core nucleotide database and the specified organism to determine if the gene is in an operon with *letA*. The resulting 20 sequences correspond to structures containing 6, 7, or 8 MCE Rings and were aligned using MUSCLE^53^. This alignment was used to generate a sequence logo using WebLogo3. The Uniprot IDs used to generate the sequence alignment are: P76272 (*E. coli*), A0A4Y5YG70 (*S. polaris*), A0A1N7PAF3 (*N. antarctica*), A0A5C6QK63 (*C. hornerae*), A0A4P6P630 (*L. sediminis*), A0A7W4Z577 (*L. lipolytica*), A0A4P7JQ50 (*Thalassotalea sp. HSM 43*), A0A1Q2M9J7 (*M. agarilyticus*), A0A2R3ITY9 (P. aeruginosa), A0A2Z3I1J2 (*Gammaproteobacteria bacterium ESL0073*), A0A090IGL8 (*M. viscosa*), A0A0×1KWU0 (*V. cholerae*), Q6LQU6 (*P. profundum*), Y1672 (*H. influenzae*), A0A2U8I7C7 (*Candidatus Fukatsuia symbiotica*), A0A085GCR5 (*B. agrestis*), A0A8E7UPQ1 (*S. enterica*), A0A8H8Z9P5 (*S. flexneri*), D4GG77 (*P. ananatis*), and B2VJ84 (*E. tasmaniensis*).

### Deep Mutational Scanning

A library containing all the possible single amino acid mutants in LetA (*n* = 8,540) was synthesized by Twist Bioscience (San Francisco, CA). Apart from the engineered mutations, these plasmid variants were identical to pBEL2071, which refactored LetA and LetB as two non-overlapping open reading frames, as they overlap by 32 bp in their native genomic context. Our LetA mutant library was divided into four ~325 bp sub-libraries (codons 1–104, 105–208, 209-320 or 321-427) due to Illumina sequencing length limitations. The sub-libraries were then handled independently. Two independent biological replicates of the deep mutational scanning experiments described below were performed starting from these sub-libraries. A *ΔletAB ΔpqiAB* strain, bBEL384, was transformed by electroporation with each sub-library and grown overnight at 37°C in LB containing 200 µg/mL carbenicillin. We obtained ~2 × 10^6 CFU for each sub-library. For the replicate experiment, ~1 × 10^7 CFU was obtained. The cultures were diluted 1:20 into fresh LB media containing 100 µg/mL carbenicillin and 50 µg/mL kanamycin and shaken (200 rpm) at 37°C until OD_600_= ~1. The cultures were plated on LB (BD Difco, catalog #DF0445–07-6) + 100 µg/mL carbenicillin (“no selection”), LB + 100 µg/mL carbenicillin + 0.105% LSB (“selection”), or LB + 100 µg/mL carbenicillin + 8% cholate (“selection”). After overnight incubation on the no selection and selection plates, colonies from each condition were separately scraped and pooled, plasmids were extracted, and amplicons were generated by PCR. The amplicons from each sub-library were then pooled in equimolar amounts to generate the no selection and selection samples. The NEBNext Ultra II Library Prep kit (New England Biologs #E7645) was used to generate the library for Illumina MiSeq 2 × 250 paired-end sequencing.

Paired-end sequencing data were mapped to a reference WT LetA sequence using the bowtie2^70^ algorithm (v2.4.1), filtered with samtools^71^ (v1.9; flags -f 2 -q 42), and overlapping paired ends were merged into a single sequence with PANDAseq^72^ (v2.11). Lastly, primer sequences used for amplicon amplification were removed using cutadapt^73^ (v1.9.1). Processed and merged reads were then analyzed using custom Python scripts to count the frequency of the LetA variants^74^. Briefly, DNA sequences were filtered by length, removing any sequence larger or smaller than the length of the expected library. Next, sequences were correctly oriented to the proper reading frame and translated to the corresponding protein sequence. Finally, the frequency of each amino acid variant at every position was counted and the counts were normalized to the sequencing depth as read counts per million. Normalized LetA variant counts were then used for calculation of the relative fitness value (*ΔE*_*i*_^*X*^), which is defined as the log frequency of observing each amino acid x at each position i in the selected versus the non-selected population, relative to the wild-type amino acid (30). The equation for this calculation is as follows:

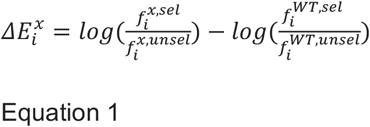

For cholate selection, we found that the square of the Pearson correlation coefficient (r^2^) between two biological replicates to be r^2^ = 0.897 (Extended Data Feig. 4B). For LSB selection, the square of the Pearson correlation coefficient is r^2^ = 0.786. These coefficients indicate replicates are in a good agreement with one another. We were able to extract meaningful fitness information for 8,478/8,540 variants for cholate and 8,504/8,540 variants for LSB. Meaningful fitness information for a mutation could not be extracted if counts were not present in either the unselected or selected dataset. For example, many mutations at position 51 had no sequence coverage due to poor representation of residue 51 mutations in the synthesized library. The relative fitness values exhibited a bimodal distribution, where the two modes represent the neutral and deleterious mutant groups (Extended Data Feig. 4C). We established a cutoff to identify mutations with relative fitness values that are substantially different from the median (0) by calculating the modified Z-score (M_i_)^75^ for each mutation using Equations 2 and 3, where x_i_ is a single data value, 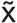 is the median of the dataset, and MAD is the median absolute deviation of the dataset:

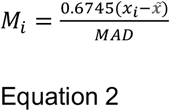

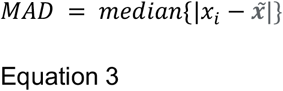

Since modified Z-scores with an absolute value of greater than 3.5 are potential outliers^75^, mutations with a Z-score of <-3.5 and >3.5 were considered to be deleterious or advantageous, respectively, to LetA function. For each residue, we calculated a tolerance score based on the number and types of amino acid substitutions that are tolerated. The tolerance scores were calculated by using a modified version of the Zvelebil similarity score^76,77^, which is based on counting key differences between amino acids.

Each mutation is given a starting score of 0.1. For each key difference (i.e. ‘small’, ‘aliphatic’, ‘proline’, ‘negative’, ‘positive’, ‘polar’, ‘hydrophobic’ and ‘aromatic), a score of 0.1 is given, such that mutations to dissimilar amino acids (e.g. alanine to arginine) contribute more to the score. If the mutation is tolerated based on our modified Z-score cut-off, the score for that particular mutation is added to a starting score of 0.1. For each sequence position, the scores for tolerant mutations were summed, then divided by the maximum score possible. A score of 1.0 therefore indicates full tolerance in that position, a score of 0 denotes no tolerance, and in-between scores suggest different levels of tolerance for that amino acid type.

In the cholate dataset, three of the 53 residues with tolerance scores <0.7 do not cluster in the three main groups (ZnRs, polar network, and outward-open pocket). The residues are A128, F141, and A272. In the LSB dataset, two residues with tolerance scores <0.7 out of 37 are located outside of the three main clusters: I131 and F141. F141 is considered functionally important in both datasets, but it is unclear what its role is, as this residue appears isolated in the membrane. Residues A272 and I131 interact with residues in the LetB TM helices, potentially stabilizing interactions between LetA and LetB in the membrane. As residue A128 precedes the helical “break” of TM3 in TMD^N^, this position may only tolerate small hydrophobic residues to maintain the structural integrity of LetA.

### Complementation Assays

*letA* and *letB* knockout strains were constructed in the *E. coli* K-12 BW25113 *ΔpqiA* background by P1 transduction from corresponding strains of the Keio collection^78^, followed by excision of the antibiotic resistance cassettes using pCP20^79^. To test the impact of *letA* (bBEL620), *letB* (bBEL621), and *letAB* (bBEL609) deletion mutants on cell viability, overnight cultures grown in LB were diluted 1:50 into fresh LB without antibiotics. The knockout strains carrying pET17b-*letAB* (Addgene #175804) or its mutants were grown in the presence of 100 µg/mL carbenicillin. Cultures were grown for ≈1.5 hours at 200 rpm and 37°C until reaching an OD600 of ~1.0, then normalized to a final OD600 of 1.0 with fresh LB. From these normalized cultures, 10-fold serial dilutions in LB were prepared in a 96-well plate, and 1 µl of each dilution was spotted onto plates containing LB agar, or LB agar supplemented with either LSB or sodium cholate. The source of LB agar used influenced the LSB and cholate phenotypes, and our agar was prepared from the following components: 10g Tryptone (Gibco, catalog #211705), 10g NaCl (Sigma-Aldrich, catalog #S3014), 5g Yeast extract (Gibco, catalog #212750), 15g Agar (BD Difco, catalog #214530) per one liter of deionized water. Plates were incubated ~18-20 hours at 37°C and then imaged using a ChemiDoc XRS+ System (Bio-Rad). Stock solutions of LSB (5% w/v) and sodium cholate (40% w/v; Thermo Fisher, catalog #A17074.18) were prepared in deionized water and stored at −80°C. At least three independent transformants were used to perform replicates for each phenotypic assay.

### Small-scale pull-down assays

Plasmid pBEL1284 was transformed into OverExpress C43 (DE3) cells (Lucigen). The *letA* or *letB* regions were mutated using Gibson assembly. Whole plasmid sequencing was performed by Plasmidsaurus using Oxford Nanopore Technology with custom analysis and annotation. Overnight cultures (LB + 100 µg/mL carbenicillin + 1% glucose) were diluted into 20 mL LB (Difco) supplemented with carbenicillin (100 µg/mL), grown at 37°C with shaking to an OD600 of ~0.9, and then induced by addition of arabinose to a final concentration of 0.2%. Cultures were further incubated at 37°C with shaking for 4 hours, and then harvested by centrifugation. The pellets were resuspended in 1 mL of lysozyme resuspension buffer (50 mM Tris pH 8.0, 300 mM NaCl, 1 mg/mL lysozyme, 25 U/mL benzonase, 1 mM TCEP) and were incubated for 1 hr at 4°C. The cells were lysed with eight cycles of a freeze-thaw method, where samples are immersed in liquid nitrogen until fully frozen and then thawed in a 37°C heat block. The lysate containing crude membrane fractions was pelleted at 20,000 g for 15 min, and resuspended in 250 µL of membrane resuspension buffer (50 mM Tris pH 8.0, 300 mM NaCl, 10% glycerol and 25 mM DDM, 1 mM TCEP), and shaken for 1 hr. The sample volume was then increased to 1 mL with 10 mM imidazole wash buffer (50 mM Tris pH 8.0, 300 mM NaCl, 10 mM imidazole) and insoluble material was pelleted at 20,000 g for 15 min. Each supernatant was then mixed with 25 µL of nickel Ni Sepharose Excel resin (Cytiva) for 30 min. The beads were pelleted at 500 g for 1 min and the supernatant removed. The beads were then washed four times with 40 mM imidazole wash buffer (50 mM Tris pH 8.0, 300 mM NaCl, 40 mM imidazole, 10% glycerol, 0.5 mM DDM, 1 mM TCEP) and finally resuspended in 50 µL of elution buffer (50 mM Tris pH 8.0, 300 mM NaCl, 300 mM imidazole, 10% glycerol, 0.5 mM DDM, 1 mM TCEP). The beads were removed by passing through an Ultrafree centrifugal filter (10,000 g for 1 min) at 4°C. The samples were then mixed with 5x SDS-PAGE loading buffer, and analyzed by SDS-PAGE and stained using InstantBlue Protein Stain (Abcam, Catalog #ab119211). At least two replicates of the experiment were performed, from independently purified proteins.

To obtain membrane fractions without cell debris, lysed samples were centrifuged at 16,000 g for 10 minutes at 4°C. The supernatant was collected and the membrane fraction was isolated by ultracentrifugation in a TLA 120.2 rotor (100,000 rpm, 15 min at 4°C). The supernatant was removed, the pellet resuspended in 500 µL of membrane resuspension buffer, and shaken for 1 hour at 4°C. Added 500 µL of 20 mM imidazole wash buffer (50 mM Tris pH 8.0, 300 mM NaCl, 20 mM imidazole, 10% glycerol, 0.5 mM DDM, 1 mM TCEP), and centrifuged at 20,000 x g for 15 min at 4°C to remove insoluble debris. The samples were then purified using affinity resin as described above.

### Generation of LetA monoclonal antibodies

Plasmid pBEL2214 was transformed into Rosetta (DE3) cells (Novagen) for protein expression and overnight cultures were grown in LB supplemented with carbenicillin (100 µg/mL), chloramphenicol (38 µg/mL) and 1% glucose at 37°C. The overnight cultures were diluted 1:50 in fresh LB media supplemented with carbenicillin (100 µg/mL) and chloramphenicol (38 µg/mL). Upon reaching an OD600 of ~0.9, protein expression was induced with the addition of L-arabinose to a final concentration of 0.2%. Cells were cultured for an additional 4 hours at 37°C with shaking, and then harvested by centrifugation. The pellets were resuspended in lysis buffer (50 mM Tris pH 8.0, 300 mM NaCl, 10% glycerol) flash frozen in liquid nitrogen and stored at −80°C. Cells were lysed by three passes through an Emulsiflex-C3 cell disruptor (Avestin), then centrifuged at 15,000g for 30 minutes at 4°C to pellet cell debris. The clarified lysate was subjected to ultracentrifugation at 37,000 rpm (182,460 g) for 45 minutes at 4°C in a Fiberlite F37L-8 × 100 Fixed-Angle Rotor (Thermo Fisher Scientific, 096-087056), and the membrane fraction was solubilized in 50 mM Tris pH 8.0, 300 mM NaCl, 10% glycerol, 25 mM DDM by rocking overnight at 4°C. Insoluble debris were pelleted by ultracentrifugation at 37,000 rpm for 45 minutes at 4°C. Solubilized membranes were then passed twice through a column containing Ni Sepharose resin (Cytiva). Eluted proteins were exchanged into low salt buffer (20 mM HEPES, pH 7.0, 25 mM NaCl, 0.5 mM DDM, 10% glycerol) using an Amicon Ultra-0.5 Centrifugal Filter Unit concentrator (MWCO 30 kDa, UFC503008) before injection into a Mono S 5/50 GL column (Cytiva). The column was eluted using a salt gradient from 25 mM to 1.5 M NaCl over 40 column volumes. The eluted proteins containing LetA were concentrated using an Amicon Ultra-0.5 Centrifugal Filter Unit concentrator (MWCO 30 kDa, UFC503008) before separation in a Superdex 200 Increase 10/300 column (Cytiva) equilibrated in gel filtration buffer (50 mM Tris-HCl pH 8.0, 150 mM NaCl, 0.5 mM DDM, 10% glycerol).

To generate LetA rat monoclonal antibodies, three six-week old Sprague Dawley rats (Taconics) were immunized with purified LetA protein (100 µg per animal per boost for 5 boosts). Immune response was monitored by ELISA to measure the serum anti-LetA IgG titer from blood samples. After a 60-day immunization course, the rat with the strongest anti-LetA immune response was terminated and 10^8^ splenocytes were collected for making hybridomas by fusing with rat myeloma cell line YB2/0, following the standard method^80^. All procedures were approved by the Cold Spring Harbor Laboratory Institutional Animal Care and Use Committee (IACUC).

To select monoclonal antibodies, supernatants collected from individual hybridoma culture media were screened by ELISA to identify hybridoma clones positive for LetA. Positive hybridoma colonies were then isolated and seeded to establish pure hybridoma clones from single cell colonies. For ELISA, purified LetA or negative control protein streptavidin diluted in LetA storage buffer (20 mM Tris, pH8, 150 mM NaCl, 10% glycerol, 0.5 mM DDM) were coated (50 ng/well) on ELISA plates (Thermo Fisher Scientific, 464718) following the manufacturer’s instructions. Prior to adding the hybridoma supernatant, the coated plate was blocked with LetA storage buffer (20 mM Tris, pH8, 150 mM NaCl, 10% glycerol, 0.5 mM DDM) containing 0.5% bovine serum albumin (BSA) at 4 °C for six hours. After blocking, hybridoma supernatants diluted in LetA buffer (1:1 dilution) were added to the ELISA plate and incubated at room temperature for 1 hour, followed by three times of extensive wash with LetA buffer. The secondary antibody (112-035-003, Jackson ImmunoResearch for anti-rat IgG HRP) was then added and incubated at room temperature for 30 min, followed by three times of extensive wash with LetA buffer. Chromogenic binding signal was developed by using 3,3’,5,5’-Tetramethylbenzidine (TMB) ultra as the HRP substrate (Thermo Fisher Scientific, 34028) following the manufacturer’s instructions. Data was collected by measuring the absorbance at 450 nm with a plate reader (Cytation 5, Agilent). The ELISA assay revealed 23 antibody clones to be strong binders of LetA, and two were found to detect LetA in cell lysates via Western blotting (clones 45 and 72). Clone 45 recognizes an epitope in the ZnR domains, and can also recognize PqiA. The epitope recognized by Clone 72 is in the N-terminal extension of LetA, and does not appear to cross-react with PqiA.

### Western blotting

To test for protein expression in the strains used for the complementation assays, 5 mL cultures of *E. coli* strains *ΔpqiAB* (bBEL385), *ΔletAB* (bBEL466), and *ΔpqiAB* Δ*letAB* (bBEL609), containing each complementation plasmid were grown to an OD600 of ~1. For plasmid containing strains, the cultures were supplemented with carbenicillin (100 µg/mL). The cells were pelleted at 4500 g for 10 mins and resuspended in 1 mL of freeze thaw lysis buffer (PBS pH 7.4, 1 mg/ml lysozyme and 1 μL/ml of benzonase (Millipore), and incubated on ice for 1 hour. The cells were lysed with eight cycles of a freeze-thaw method, where samples are immersed in liquid nitrogen until fully frozen and then thawed in a 37°C heat block. After lysis, the cells were pelleted at ~20,000 g for 15 minutes and the pellets were resuspended in 100 μl of SDS-PAGE loading buffer. Each sample (10 μL) was separated on an SDS-PAGE gel and transferred to a nitrocellulose membrane using the Trans-Blot Turbo Transfer System (Bio-Rad Laboratories). The membranes were blocked in Phosphate Buffered Saline Tween20 (PBST, 1X PBS + 0.1% Tween20) containing 5% milk for 1 hour at room temperature. To probe LetA, the membranes were incubated with primary antibody in PBST + 5% BSA, either rat monoclonal anti-LetA clone 45 or clone 72 at a final concentration of 0.5 µg/mL or 2 µg/mL, respectively. To probe LetB or BamA, membranes were incubated with rabbit polyclonal anti-LetB (1:10,000 dilution) in PBST + 5% BSA or rabbit polyclonal anti-BamA^31^ (1:2000 dilution) in PBST + 5% BSA, respectively. Membranes were incubated in primary antibody solution for either 1 hour at room temperature or overnight at 4°C with agitation. The membranes were then washed 3 times with PBST and incubated with Goat anti-rat IgG IRDye® 680RD (LI-COR Biosciences, Catalog #925-68071), Goat anti-rabbit IgG IRDye® 680CW (LI-COR Biosciences, Catalog #926-68071) or Goat anti-rabbit IgG IRDye® 800CW (LI-COR Biosciences, Catalog #925-32211) secondary antibodies in Intercept (TBS) Blocking Buffer (LI-COR Biosciences, Catalog #927-60003) for 1 hour at room temperature with agitation. The membranes were then washed 3 times with PBST and imaged on a LI-COR Odyssey Classic.

### ICP-MS

Plasmid pBEL1284 was modified to encode only LetA with a C-terminal 2xQH-7xHis tag to yield pBEL2214. The plasmid was transformed into Rosetta (DE3) cells (Novagen) for protein expression and overnight cultures were grown in LB supplemented with carbenicillin (100 µg/mL), chloramphenicol (38 µg/mL) and 1% glucose at 37°C. The overnight cultures were diluted 1:50 in fresh LB media supplemented with carbenicillin (100 µg/mL) and chloramphenicol (38 µg/mL). Upon reaching an OD600 of ~0.6, the media was supplemented with 1X metals (50 µM FeCl_3_, 20 µM CaCl_2_, 10 µM MnCl_2_, 10 µM ZnSO_2_, 2 µM CoCl_2_, 2 µM CuCl_2,_ 2 µM NiCl_2_, 2 µM Na_2_MoO_4_, 2 µM Na_2_SeO_3_, 2 µM H_3_BO_3_) and protein expression was induced with the addition of L-arabinose to a final concentration of 0.2%. Cells were cultured for an additional 4 hours at 37°C with shaking, and then harvested by centrifugation. The pellets were resuspended in lysis buffer (50 mM Tris pH 8.0, 300 mM NaCl, 10% glycerol, 1 mM TCEP), flash frozen in liquid nitrogen and stored at −80°C. Cells were lysed by three passes through an Emulsiflex-C3 cell disruptor (Avestin), then centrifuged at 15,000g for 30 minutes at 4°C to pellet cell debris. The clarified lysate was subjected to ultracentrifugation at 37,000 rpm (182,460 g) for 45 minutes at 4°C in a Fiberlite F37L-8 × 100 Fixed-Angle Rotor (Thermo Fisher Scientific, 096-087056), and the membrane fraction was solubilized in 50 mM Tris pH 8.0, 300 mM NaCl, 10% glycerol, 25 mM DDM, 1 mM TCEP by rocking overnight at 4°C. Insoluble debris were pelleted by ultracentrifugation at 37,000 rpm for 45 minutes at 4°C. Solubilized membranes were then passed twice through a column containing Ni Sepharose Excel resin (Cytiva). Eluted proteins were concentrated using the Amicon Ultra-0.5 Centrifugal Filter Unit concentrator (MWCO 30 kDa, UFC503008) before separation on the Superdex 200 Increase 10/300 column (Cytiva) equilibrated in gel filtration buffer (50 mM Tris-HCl pH 8.0, 150 mM NaCl, 0.5 mM DDM, 10% glycerol, 1 mM TCEP). LetA concentrations were quantified by gel densitometry using bovine serum albumin standards.

Samples for ICP-MS analysis were prepared by adding 0.1 mL trace-metal grade nitric acid to 1.9 mL of protein sample as provided. Samples were analyzed using a Perkin Elmer NexION 350D inductively coupled plasma mass spectrometer. All liquid samples were infused into the nebulizer via peristaltic pump at 0.3 mL/min. For full scan elemental analysis, the instrument “TotalQuant” method was employed, using factory response factors. For quantitative analysis of zinc, calibrators were prepared by dilution of certified single element standard (Perkin Elmer) with 5% nitric acid, and these were used to generate a standard response curve. Except for zinc, none of the 79 other elements tested, such as iron, nickel and cobalt, were enriched in the protein sample relative to the buffer control. These results suggest that the ZnR domains of LetA bind zinc, though we note that the metal binding properties of similar ZnR proteins can be sensitive to the experimental conditions^81^.

### BPA cross-linking assays

OverExpress C43 (DE3) cells were transformed with plasmids to express LetAB (either WT or mutant forms derived from pBEL1284) or LetB (either WT or mutant forms derived from pBEL2782). The cells were co-transformed with pEVOL-pBpF (Addgene #31190) to encode a tRNA synthetase/tRNA pair for the *in vivo* incorporation of p-benzoyl-l-phenylalanine (BPA, Bachem, Catalog #F-2800.0005) in *E. coli* proteins at the amber stop codon, TAG^82^. Bacterial colonies were inoculated in LB broth supplemented with carbenicillin (100 μg/mL) and chloramphenicol (38 μg/mL) and grown overnight at 37°C. The following day, bacteria were pelleted and resuspended in ^32^P Labeling Medium (a low phosphate minimal media we optimized starting from LS-5052^83^: 1 mM Na_2_HPO_4_, 1 mM KH_2_PO_4_, 50 mM NH_4_Cl, 5 mM Na_2_SO_4_, 2 mM MgSO_4_, 20 mM Na_2_-Succinate, 0.2x trace metals and 0.2% glucose) supplemented with carbenicillin (100 μg/mL) and chloramphenicol (38 μg/mL) and inoculated 1:33 in 20 mL of the same medium. Bacteria were grown to OD600 = ~0.6-0.7 and a final concentration of 0.2% L-arabinose, 0.5 mM BPA, and 500 μCi ^32^P orthophosphoric acid (PerkinElmer, Catalog #NEX053010MC) were added and left to induce overnight at room temperature.

The following day, the cells were harvested by centrifugation (4,500 g for 10 min) and resuspended in 1 mL of phosphate buffer saline (PBS, pH 7.4), and the ‘crosslinked’ samples underwent crosslinking by treatment with 365 nM UV in a Spectrolinker for 30 min. Both the crosslinked and uncrosslinked cells were pelleted (6,000 g for 2 min) and resuspended in 1 mL of lysozyme resuspension buffer (50 mM Tris pH 8.0, 300 mM NaCl, 1 mg/mL lysozyme, 25 U/mL benzonase) and were incubated for 1 hr at 4°C. The cells then underwent eight cycles of freeze-thaw lysis by alternating between liquid nitrogen and a 37°C heat block. The lysate was pelleted at 20,000 g for 15 min, and the pellets were resuspended in 250 µL of membrane resuspension buffer (50 mM Tris pH 8.0, 300 mM NaCl, 10% glycerol and 25 mM DDM), and shaken for 1 hr. The sample volume was then increased to 1 mL with 10 mM imidazole wash buffer (50 mM Tris pH 8.0, 300 mM NaCl, 10 mM imidazole) and insoluble material was pelleted at 20,000 g for 15 min. Each supernatant was then mixed with 25 µL of nickel Ni Sepharose Excel resin (Cytiva) for 30 min. The beads were pelleted at 500 g for 1 min and the supernatant collected. The beads were then washed four times with 40 mM wash buffer (50 mM Tris pH 8.0, 300 mM NaCl, 40 mM imidazole, 10% glycerol, 0.5 mM DDM) and finally resuspended in 50 µL of elution buffer (50 mM Tris pH 8.0, 300 mM NaCl, 300 mM imidazole, 10% glycerol, 0.5 mM DDM). The samples were then mixed with 5x SDS-PAGE loading buffer, and the beads spun down at 12,000 g for 2 min. Eluted protein was analyzed by SDS-PAGE and stained using InstantBlue Protein Stain (Abcam, Catalog #ab119211). Relative loading of the LetA or LetB monomer band on the gel was estimated integrating the density of the corresponding bands in the InstantBlue-stained gel in ImageJ (Rueden et al., 2017), and this was used to normalize the amount of protein loaded on a second gel, to enable more accurate comparisons between samples. The normalized gel was stained with InstantBlue, and ^32^P signal was detected using a phosphor screen and scanned on a Typhoon scanner (Amersham). At least two biological replicates of the experiment were performed, starting with an independent protein expression culture grown on a different day.

### Disulfide-crosslinking assays

To perform these assays, we generated variants of our split-LetA construct. As one cysteine from each pair is within TMD^N^ and the second Cys from each pair is within TMD^C^, we introduced these cysteine pairs into a variant of our split-LetA construct with non-essential cysteines removed (C124S, C266S, C343S; ΔCysSplitLetA), to facilitate the detection of the crosslink of interest. A crosslinking event is predicted to lead to covalent linkage between TMD^N^ and TMD^C^, resulting in a dimer with a large molecular weight shift on SDS-PAGE relative to either domain alone. The metal-coordinating cysteines of the ZnR domains cannot be mutated without affecting LetA function, but are likely protected from maleimide crosslinkers by the bound zinc ion. Given that our DMS data suggests that Q180 and R380 can tolerate mutations to cysteines, we selected these residues to probe the alternative conformation.

OverExpress C43 (DE3) cells (Lucigen) containing pBEL2802 or its mutants were grown overnight at 37°C in LB medium supplemented with carbenicillin (100 µg/mL) and 1% glucose. Overnight cultures were diluted 1:50 to 20 mL of fresh LB media containing carbenicillin (100 µg/mL). Cells were grown to an OD600 of ~0.8, pelleted by centrifugation (4,500 g for 10 mins at 4°C), and resuspended in 1.5 mL PBS pH 7.4. From this stock, 500 µL cell suspension was pipetted into two separate Eppendorf tubes, and either treated with 3% dimethyl sulfoxide (DMSO; solvent used for dissolving cross-linkers) or with 1 mM BMOE (Thermo Scientific Pierce, Catalog #PI22323). To cap unreacted cysteines, both the DMSO- and BMOE-treated samples were incubated with 2 mM N-ethylmaleimide (NEM, Thermo Scientific Pierce, Catalog #PI23030) and incubated for 10 min at RT while rotating in the dark. To quench unreacted BMOE, the cells were incubated with 10 mM L-cysteine (Sigma, Catalog #168149) for 10 min at RT while rotating. The cells were pelleted by centrifugation (6,000 g for 2 min), flash frozen in liquid nitrogen, and stored −80°C. To lyse the cells, the pellets were resuspended in 1 mL of lysozyme resuspension buffer (50 mM Tris pH 8.0, 300 mM NaCl, 1 mg/mL lysozyme, 25 U/mL benzonase, 1 mM DTT) and incubated for 1 hr at 4°C. The cells then underwent eight cycles of freeze-thaw lysis by alternating between liquid nitrogen and a 37°C heat block. The lysate was pelleted at 20,000 g for 15 min, and resuspended in 250 µL of membrane resuspension buffer (50 mM Tris pH 8.0, 300 mM NaCl, 10% glycerol and 25 mM DDM, 1 mM DTT), and shaken for 1 hr. Insoluble debris was removed by centrifugation at 20,000 ag for 15 min at 4°C. For each sample, 20 µL of the supernatant was mixed with 20 µL 2X SDS-PAGE loading dye supplemented with fresh 50 mM DTT. The samples were heated to 50°C for 15 min, and then 10 µL of the sample was loaded to an SDS-PAGE gel. The LetA bands were probed by Western blotting using the monoclonal anti-LetA antibody (clone #72).

### AlphaFold2 predictions

To identify the alternative conformations of LetA, we used AlphaFold2_multimer_version3 via Colabfold^84^. The sequence of LetA was retrieved from the MG1655 reference genome in the NCBI database. Program outputs yielded five ranked models, each with 32 samples or “seeds”, resulting in 160 predictions. Finding ambiguity in the coevolutionary signal was achieved by reducing the depth of the input multiple sequence alignments (16:32), enabling “dropout”, and setting “recycling”, which is the number of times the structure is fed into the neural network, to 0^84,85^. While many predictions showed ZnR^N^ and ZnR^C^ interacting with each other, 37 models showed different degrees of separation between the two ZnRs. In addition, five predictions exhibited severe clashes in ZnR^C^. These observations made it difficult to interpret the cytoplasmic region, which also includes the unstructured N- and C-terminal regions that are not observed in our cryo-EM density, and we therefore focused on the TMD region. The RMSD heatmap was built as follows: First, each of the 161 PDBs (LetA cryo-EM structure and 160 models generated by AlphaFold) was aligned with all others PDBs using “align” function from PyMol Molecular Graphics System (Version 3.0 Schrödinger, LLC), restricting the alignment to the carbon atoms and number of cycles to 0. Then, a matrix with the 161 PDBS in x and y was filled with the RMSD. Finally, the dendrogram was computed using the fastcluster python package^86^ (using the Ward method and Euclidean metric). For each cluster, the representative model was selected as the one having the lowest average RMSD within that cluster.

### System preparation for MD simulations

The LetAB complex used in the MD simulations was constructed by integrating the cryo-EM resolved structure with the AlphaFold2 multimer predicted model using Chimera and Coot. Missing residues in the C-terminal region of LetA (residues 419-427) were reconstructed using AlphaFold2, while the N-terminal disordered region (residues 1-26) was excluded due to its low pLDDT score. Similarly, for LetB, the N-terminal absent residues (residues 1-13) were omitted for the same reason. We retained the TM helices and the first MCE Ring of LetB (residues <= 160) to preserve the native environment surrounding LetA, while the remaining portions of LetB were excluded to minimize the system size. Additionally, the two absent TM helices (residues 14-45) of LetB were modeled using AlphaFold2. The N-termini of LetA and LetB were capped with an acetylated N-terminus (ACE), while the C-termini of LetA and LetB were capped with a standard C-terminus (CTER) and a methylamidated C-terminus (CT3), respectively. Protonation states of titratable residues were determined using PropKa3^87,88^. The orientation of the protein complex relative to the membrane was established using the Positioning of Proteins in Membranes (PPM) 3.0 web server^89^, and the resultant oriented protein complex was embedded into a native Gram-negative bacterial inner membrane (IM) using Membrane Builder module in CHARMM-GUI^90,91^. Each membrane leaflet consisted of 1-palmitoyl-2-(cis-9,10-methylene-hexadecanoyl)-phosphatidylethanolamine (PMPE, 16:0/cy17:0), 1-palmitoyl-2-oleoyl-phosphatidylethanolamine (POPE, 16:0/18:1(9Z)), 1-pentadecanoyl-2-(cis-9,10-methylene-hexadecanoyl)-phosphatidylethanolamine (QMPE, 15:0/cy17:0), 1-oleoyl-2-(9Z-hexadecenoyl)-phosphatidylethanolamine (OYPE, 18:1(9Z)/16:1(9Z)), 1-palmitoyl-2-(cis-9,10-methylene-hexadecanoyl)-phosphatidylglycerol (PMPG, 16:0/cy17:0), 1-palmitoyl-2-(9Z-hexadecenoyl)-phosphatidylglycerol (PYPG, 16:0/16:1(9Z)), and 1,1′-palmitoyl-2,2′-(11Z-vacenoyl)-cardiolipin (PVCL2, 1’-[16:0/18:1(11Z)],3’-[16:0/18:1(11Z)]) in ratios of 46, 13, 12, 8, 10, 9, and 2, respectively. The cryo-EM resolved lipid was elongated to PMPE (in simulations performed with Lipid 1), as it represents the most abundant phospholipid in the Gram-negative bacterial IM. To assess the functional role of the cryo-EM resolved lipid, we constructed systems both with and without Lipid 1. For each scenario, lipid positions within the membrane were randomly shuffled using the Membrane Mixer plugin^92^ in VMD to minimize any biases from initial lipid placement, resulting in three replicas for each condition, totaling six replicas overall. Finally, the resulting protein-membrane systems were solvated and neutralized with 0.15 M NaCl.

### MD simulation protocols

All the MD simulations were executed using the NAMD^93^ program. CHARMM36m^94^ and CHARMM36^95^ force fields were employed for the proteins and lipids, respectively. The TIP3P model was used for water molecules^96^. Temperature was maintained at 310 K via a Langevin thermostat with a damping coefficient of γ = 1 ps^-1^, and pressure was held at 1 bar through the Nosé-Hoover piston^97,98^. The Particle-mesh Ewald (PME)^99^ method was used for calculating long-range electrostatic interactions within periodic boundary conditions at every time step. Non-bonded interactions were calculated with a cutoff of 12 Å, and a switching distance set at 10 Å. To accommodate volumetric changes in the system, a flexible cell was employed, allowing independent fluctuations in three dimensions while preserving a constant x/y ratio for the membrane. The SHAKE^100^ and SETTLE^101^ algorithms were used to constrain bonds involving hydrogen atoms. For the initial equilibration and production runs, a 4-fs timestep was used, facilitated by hydrogen mass repartition (HMR)^102,103^ to accelerate the simulations. A 2-fs timestep was applied without HMR for non-equilibrium simulations involving collective variables (colvars), as well as any subsequent equilibrium simulations.

To equilibrate the membrane-protein systems, each system underwent an initial 10,000 steps of energy minimization using the steepest descent algorithm, followed by equilibrations with gradually reduced harmonic restraints^104^. Initially, only the phospholipid tails were allowed to move without any constraints for 1 ns in an NVT ensemble (constant Number of particles, Volume, and Temperature), followed by a 10-ns phase where all components excluding the protein were unrestrained in an NPT ensemble (constant Number of particles, Pressure, and Temperature). Subsequently an additional 10-ns simulation was performed to allow all components except the protein backbone to move freely under NPT. A force constant of 10 kcal/mol/Å^2^ was applied to the restrained atoms. Finally, all restraints were removed, and each system was subjected to a 2-µs production run. In each of the six replicas (3 with Lipid 1 present at the start and 3 without), a lipid moved spontaneously upwards into the Lipid 2 site in the central cleft. The identity of Lipid 2 varied, and was PMPE in four replicas, PYPG in one replica and cardiolipin in one replica.

### Exploring the potential pathway of phospholipid transport by steered MD (SMD) simulations

To elucidate the potential mechanisms and pathways for phospholipid transport and evaluate the feasibility of accommodating a phospholipid within the periplasmic pocket in TMD^C^, a series of SMD simulations were conducted. These simulations employed collective variables (COLVARS)^105^ to direct the upward movement of Lipid 2 from the central cleft, starting from the poses identified in prior MD simulations. Forces were applied to the COM of three specific regions of Lipid 2: the headgroup, the terminal six carbons of the tail closest to the bottom of the periplasmic pocket, and the terminal six carbons of both tails. Each pulling scenario used a stepwise protocol with the distanceZ COLVARS to avoid unintended pathways, ensuring that the pulled atom group traversed the periplasmic pocket in TMD^C^. The lipid was initially steered towards the bottom of the periplasmic pocket in TMD^C^, followed by movement towards the middle of the pocket, and ultimately to the top of the pocket. Initial configurations for these SMD simulations were derived from the final frame of the 2-µs production run of replica 2 for the system with Lipid 1 and replica 3 for the system without Lipid 1, as they represented the lipids with the most elevated positions for each condition. This results in six distinct SMD setups (2 initial configurations × 3 pulling scenarios). To ensure optimal interactions between the pulled lipid and its surrounding environment, the pulling velocity was set to 0.2 Å/ns with a force constant of 10.0 kcal/mol/Å^2^. The duration of each simulation is provided in Supplementary Table 3. Throughout the SMD simulations, the centerToReference and rotateToReference options in NAMD were enabled to align LetA with its initial conformation before calculating distances and forces at each timestep, which avoided the effects of protein translation and rotation on the applied force. Additionally, the *z*-center of LetA was harmonically restrained using the harmonicWalls function in COLVARS with a force constant of 10.0 kcal/mol/Å^2^ and the lower and upper wall thresholds set at −2 Å and 2 Å, respectively. This restraint prevented the global upward movement of LetA induced by the applied forces on Lipid 2, which could otherwise distort the local membrane structure.

Following the completion of SMD simulations, the systems underwent an additional 10-ns equilibration phase, during which the protein backbone and the heavy atoms of the pulled lipid were harmonically restrained with a force constant of 10 kcal/mol/Å^2^. All restraints were subsequently removed, and a 300 ns production run was conducted for each system.

### Water bridge network analysis

To investigate the potential role of polar residues (D181, K178, S321, K328, S364, D367, and T402) in TMD^c^ in a proton shuttle pathway, hydrogen bonds were analyzed on a frame-by-frame basis across simulation replicas. This included hydrogen bonds formed directly between the residues, between each residue and adjacent water molecules, and among the water molecules themselves. Hydrogen bonds were defined using the geometric criteria: a donor (D)-hydrogen (H) distance cutoff of 1.2 Å, a donor-acceptor (A) distance cutoff of 3.0 Å, and a minimum D-H-A angle of 120°, ensuring the inclusion of only well-structured hydrogen bonds.

The occupancy of hydrogen bonds is defined as the fraction of total simulation frames in which a given hydrogen bond is observed. Water bridges were classified by their order: direct residue–residue hydrogen bonds with no intervening water molecules were designated as zero-water (0-W) bridges, while those involving one, two, or three intervening water molecules (1-W, 2-W, and 3-W bridges) were identified by systematically linking residue-to-water and water-to-water hydrogen bonds, thereby constructing higher-order networks. To visually represent the water bridge networks with the highest occupancy, principal component analysis (PCA) was applied to project the spatial arrangement of residues into two dimensions. Each residue was depicted as a node, with edges connecting the nodes to represent the highest-occupancy water bridge between the residues, and the thickness of the edges indicating the relative occupancy of the corresponding water bridge.

To calculate the predicted pKa of ionizable residues in LetA, the crosslinked LetA structure was uploaded into the PropKa online server: https://www.ddl.unimi.it/vegaol/propka.html^106,107^

### Sample Preparation for LC-MS Lipidomics

Three replicates for each protein were analyzed, with each replicate containing a purified protein in detergent, a detergent buffer negative control, and an isolated *E. coli* membrane positive control. The lipids from each sample were extracted via Folch extraction. Briefly, varying amounts of sample, chloroform, methanol, EquiSPLASH LIPIDOMIX (Avanti Polar Lipids) internal standards, and water were combined as described in Supplementary Table 4. The bottom layer was extracted, dried using N_2_ gas, and resuspended in 100 µL of LC-MS grade methanol. Each sample was analyzed using data-dependent acquisition (DDA) LC-MS/MS. After DDA data collection, 30 µL of each sample was diluted with 50 µL LC-MS grade methanol to ensure sufficient volume for triplicate MS1-only injections for quantitation.

### Collection of Data-Dependent LC-MS/MS Lipidomics Data

Lipids were separated prior to MS analysis using a 21-minute trap-and-elute method as previously described^108,109^. A Waters XBridge Direct Connect HP C_8_ column (10 µm, 2.1 × 30 mm) was used as the trap column, and a Waters Premier Acquity UPLC CSH C_18_ column (1.7 µm, 2.1 × 100 mm) was used for analytical separations. Gradients for the C_8_ (trap) and C_18_ (analytical) columns were controlled by the Alpha and Beta pumps respectively and are provided in Table 2. The column selection valve position was changed at 0.50 minutes, allowing the Beta pump to flow through both the C_8_ and C_18_ columns. At 17.50 minutes the valve reverted to the initial position to allow for washing and re-equilibration of the C_8_ and C_18_ columns by the Alpha and Beta pumps, respectively. A 10 µL sample injection volume was used, and the column compartment was held at 60 °C.

Samples were ionized via ESI in negative mode and introduced into a Synapt XS mass spectrometer operated in sensitivity mode. The capillary and sampling cone voltages were set to 2.45 kV and 49 V, respectively. The source offset was 80 V, the source temperature was 120°C, and the desolvation temperature was 250°C. A top-5 DDA method was applied, with an accumulated TIC threshold of 100000 and a maximum acquisition time of 0.25 seconds. MS1 and MS2 spectra were collected from 50 – 2000 Th in continuum mode at a resolution of 10,000 with a scan speed of 0.1 seconds, and fragmentation was performed using collision-induced dissociation (CID) with a collision energy ramp from 20–40 V in the trap cell. Dynamic exclusion was used with an exclusion time of 15 seconds and an exclusion width of 0.5 Da. A fixed exclusion range of 50–450 Th was used to minimize selection of non-lipid precursors. Blank injections of isopropyl alcohol (IPA) were performed every three samples using the same instrumental methods.

### Collection of MS1-Only LC-MS Lipidomics Data

After using DDA methods to identify the lipids, we used MS1-only scans to perform quantitation with accurate mass and retention time alignment, as described previously^108,109^. 10 µL of each diluted sample was loaded and separated as described above. MS1 scan parameters were identical to DDA MS1 scans. Triplicate injections of each sample were performed in a randomized order. 10 µL IPA injections were performed after each sample run to mitigate potential carryover between samples.

### Lipid Library Construction using DDA Lipidomics Data

All DDA files were centroided using MSConvert, and mass calibration was performed using an in-house Python script using the known masses and retention times of the EquiSPLASH lipids. Lipid identification based on the calibrated DDA data files was performed using MS-DIAL (version 5.3)^110^. A minimum peak height of 1000 and mass slice width of 0.1 Da were used for peak detection. Only CL, PE, and PG lipids were searched, as these are the most prevalent lipids in *E. coli*. MS1 and MS2 accurate mass tolerances of 0.025 Da were used, and both [M-H]- and [M+CH_3_COO^-^]- adducts were allowed. For alignment, a retention time tolerance of 0.5 minutes and mass tolerance of 0.015 Da were used. A set of high-quality lipid identifications was then manually checked to produce a lipid library to be used for MS1-based lipid quantification.

### Processing of MS1-Only Lipidomics Data

MS1-only files were centroided and calibrated in the same way as DDA files, with the addition of retention time calibration using the known retention times of EquiSPLASH-spiked lipids. Calibrated MS1 files were loaded into Skyline (v23.1.0) and searched against the DDA-constructed lipid library^111^. An ion match tolerance of 0.05 Th and mass accuracy of 10 ppm were used. To ensure accurate quantification, each extracted-ion-chromatogram integration was manually checked and adjusted as necessary. Raw peak areas were then standardized to sample volume and total identified lipid area.

Finally, we compared the enrichment of specific lipid classes between the isolated protein samples and the starting membranes. First, within each individual sample, the total intensity of each lipid class (CL, PE, and PG) was calculated by summing the standardized areas of each individual lipid in each class. The average total peak area for each class was then calculated using the summed areas from each replicate. To compare the lipid composition in the protein samples relative to the *E. coli* membranes, the average fold change was then calculated for each class between the protein and membrane samples. Standard deviations were propagated through the averaging, and a 95% confidence interval for the fold-change was calculated. A two-sample t-test comparing the means of each lipid class area between the protein and membrane samples was performed, and p-values were corrected for multiple testing using the Benjamini-Hochberg method. A significance level of 0.05 was used to determine statistically significant differences.

## REFERENCES

1. Drew, D. & Boudker, O. Ion and lipid orchestration of secondary active transport. Nature 626, 963–974 (2024).

2. Gouaux, E. & Mackinnon, R. Principles of selective ion transport in channels and pumps. Science 310, 1461–1465 (2005).

3. Sugano, K. et al. Coexistence of passive and carrier-mediated processes in drug transport. Nat. Rev. Drug Discov. 9, 597–614 (2010).

4. Thomas, C. & Tampé, R. Structural and mechanistic principles of ABC transporters. Annu. Rev. Biochem. 89, 605–636 (2020).

5. Shi, Y. Common folds and transport mechanisms of secondary active transporters. Annu. Rev. Biophys. 42, 51–72 (2013).

6. Drew, D. & Boudker, O. Shared molecular mechanisms of membrane transporters. Annu. Rev. Biochem. 85, 543–572 (2016).

7. Palmgren, M. P-type ATPases: Many more enigmas left to solve. J. Biol. Chem. 299, 105352 (2023).

8. Mulkidjanian, A. Y., Makarova, K. S., Galperin, M. Y. & Koonin, E. V. Inventing the dynamo machine: the evolution of the F-type and V-type ATPases. Nat. Rev. Microbiol. 5, 892–899 (2007).

9. Ekiert, D. C. et al. Architectures of Lipid Transport Systems for the Bacterial Outer Membrane. Cell 169, 273–285.e17 (2017).

10. Malinverni, J. C. & Silhavy, T. J. An ABC transport system that maintains lipid asymmetry in the Gram-negative outer membrane. Proc. Natl. Acad. Sci. U. S. A. 106, 8009–8014 (2009).

11. Pandey, A. K. & Sassetti, C. M. Mycobacterial persistence requires the utilization of host cholesterol. Proc Natl Acad Sci U S A 105, 4376–4380 (2008).

12. Yang, Y., Zienkiewicz, A., Lavell, A. & Benning, C. Coevolution of domain interactions in the chloroplast TGD1, 2, 3 lipid transfer complex specific to Brassicaceae and Poaceae plants. Plant Cell 29, 1500–1515 (2017).

13. Awai, K., Xu, C., Tamot, B. & Benning, C. A phosphatidic acid-binding protein of the chloroplast inner envelope membrane involved in lipid trafficking. Proc. Natl. Acad. Sci. U. S. A. 103, 10817–10822 (2006).

14. Isom, G. L. et al. MCE domain proteins: conserved inner membrane lipid-binding proteins required for outer membrane homeostasis. Sci. Rep. 7, 8608 (2017).

15. Chen, J. et al. Structure of an endogenous mycobacterial MCE lipid transporter. Nature 620, 445–452 (2023).

16. Coudray, N. et al. Structure of bacterial phospholipid transporter MlaFEDB with substrate bound. Elife 9, (2020).

17. Tang, X. et al. Structural insights into outer membrane asymmetry maintenance in Gram-negative bacteria by MlaFEDB. Nat. Struct. Mol. Biol. 28, 81–91 (2021).

18. Thong, S. et al. Defining key roles for auxiliary proteins in an ABC transporter that maintains bacterial outer membrane lipid asymmetry. Elife 5, (2016).

19. Powers, M. J., Simpson, B. W. & Trent, M. S. The Mla pathway in Acinetobacter baumannii has no demonstrable role in anterograde lipid transport. Elife 9, (2020).

20. Zhou, C. et al. Structural insight into phospholipid transport by the MlaFEBD complex from P. aeruginosa. J. Mol. Biol. 433, 166986 (2021).

21. Mann, D. et al. Structure and lipid dynamics in the maintenance of lipid asymmetry inner membrane complex of A. baumannii. Commun. Biol. 4, 817 (2021).

22. Chi, X. et al. Structural mechanism of phospholipids translocation by MlaFEDB complex. Cell Res. 30, 1127–1135 (2020).

23. Zhang, Y., Fan, Q., Chi, X., Zhou, Q. & Li, Y. Cryo-EM structures of Acinetobacter baumannii glycerophospholipid transporter. Cell Discov. 6, 86 (2020).

24. Warakanont, J. et al. Chloroplast lipid transfer processes in Chlamydomonas reinhardtii involving a TRIGALACTOSYLDIACYLGLYCEROL 2 (TGD2) orthologue. Plant J. 84, 1005–1020 (2015).

25. Nazarova, E. V. et al. Rv3723/LucA coordinates fatty acid and cholesterol uptake in Mycobacterium tuberculosis. Elife 6, (2017).

26. Nazarova, E. V. et al. The genetic requirements of fatty acid import by Mycobacterium tuberculosis within macrophages. Elife 8, (2019).

27. Chen, Y., Wang, Y. & Chng, S.-S. A conserved membrane protein negatively regulates Mce1 complexes in mycobacteria. Nat. Commun. 14, 5897 (2023).

28. Forrellad, M. A. et al. Role of the Mce1 transporter in the lipid homeostasis of Mycobacterium tuberculosis. Tuberculosis (Edinb.) 94, 170–177 (2014).

29. Rank, L., Herring, L. E. & Braunstein, M. Evidence for the Mycobacterial Mce4 transporter being a multiprotein complex. J. Bacteriol. 203, (2021).

30. García-Fernández, J., Papavinasasundaram, K., Galán, B., Sassetti, C. M. & García, J. L. Molecular and functional analysis of the mce4 operon in Mycobacterium smegmatis. Environ. Microbiol. 19, 3689–3699 (2017).

31. Isom, G. L. et al. LetB Structure Reveals a Tunnel for Lipid Transport across the Bacterial Envelope. Cell 181, 653–664.e19 (2020).

32. Liu, C., Ma, J., Wang, J., Wang, H. & Zhang, L. Cryo-EM structure of a bacterial lipid transporter YebT. J. Mol. Biol. 432, 1008–1019 (2020).

33. Nakayama, T. & Zhang-Akiyama, Q.-M. pqiABC and yebST, Putative mce Operons of Escherichia coli, Encode Transport Pathways and Contribute to Membrane Integrity. J. Bacteriol. 199, (2017).

34. Vieni, C., Coudray, N., Isom, G. L., Bhabha, G. & Ekiert, D. C. Role of Ring6 in the Function of the E. coli MCE Protein LetB. J. Mol. Biol. 434, 167463 (2022).

35. Krishna, S. S., Majumdar, I. & Grishin, N. V. Structural classification of zinc fingers: survey and summary. Nucleic Acids Res. 31, 532–550 (2003).

36. van Kempen, M. et al. Fast and accurate protein structure search with Foldseek. Nat. Biotechnol. 42, 243–246 (2024).

37. Susa, K. J., Kruse, A. C. & Blacklow, S. C. Tetraspanins: structure, dynamics, and principles of partner-protein recognition. Trends Cell Biol. 34, 509–522 (2024).

38. Ben-Yaacov, A. et al. Molecular mechanism of AMPA receptor modulation by TARP/stargazin. Neuron 93, 1126–1137.e4 (2017).

39. Suzuki, H., Tani, K. & Fujiyoshi, Y. Crystal structures of claudins: insights into their intermolecular interactions. Ann. N. Y. Acad. Sci. 1397, 25–34 (2017).

40. Zimmerman, B. et al. Crystal structure of a full-length human tetraspanin reveals a cholesterol-binding pocket. Cell 167, 1041–1051.e11 (2016).

41. Liu, S., Sukumar, N. & Li, W. Takifugu rubripes VKOR-like C138S mutant with vitamin K1. Preprint at 10.2210/pdb6wv8/pdb (2020).

42. Li, P., Lees, J. A., Lusk, C. P. & Reinisch, K. M. Cryo-EM reconstruction of a VPS13 fragment reveals a long groove to channel lipids between membranes. J Cell Biol 219, (2020).

43. Cooper, B. F. et al. Phospholipid transport across the bacterial periplasm through the envelope-spanning bridge YhdP. J. Mol. Biol. 168891 (2024).

44. Suits, M. D. L., Sperandeo, P., Dehò, G., Polissi, A. & Jia, Z. Novel structure of the conserved gram-negative lipopolysaccharide transport protein A and mutagenesis analysis. J Mol Biol 380, 476–488 (2008).

45. Kaur, D., Khaniya, U., Zhang, Y. & Gunner, M. R. Protein motifs for proton transfers that build the transmembrane proton gradient. Front. Chem. 9, 660954 (2021).

46. Perales-Calvo, J., Lezamiz, A. & Garcia-Manyes, S. The Mechanochemistry of a Structural Zinc Finger. J. Phys. Chem. Lett. 6, 3335–3340 (2015).

47. Jumper, J. et al. Highly accurate protein structure prediction with AlphaFold. Nature 596, 583–589 (2021).

48. Wayment-Steele, H. K. et al. Predicting multiple conformations via sequence clustering and AlphaFold2. Nature 625, 832–839 (2024).

49. Del Alamo, D., Sala, D., Mchaourab, H. S. & Meiler, J. Sampling alternative conformational states of transporters and receptors with AlphaFold2. Elife 11, (2022).

50. Lambert, E., Mehdipour, A. R., Schmidt, A., Hummer, G. & Perez, C. Evidence for a trap-and-flip mechanism in a proton-dependent lipid transporter. Nat. Commun. 13, 1022 (2022).

51. Li, Y., Orlando, B. J. & Liao, M. Structural basis of lipopolysaccharide extraction by the LptBFGC complex. Nature 567, 486–490 (2019).

52. Klesse, G., Rao, S., Sansom, M. S. P. & Tucker, S. J. CHAP: A Versatile Tool for the Structural and Functional Annotation of Ion Channel Pores. J. Mol. Biol. 431, 3353–3365 (2019).

53. Edgar, R. C. MUSCLE: multiple sequence alignment with high accuracy and high throughput. Nucleic Acids Res. 32, 1792–1797 (2004).

54. Crooks, G. E., Hon, G., Chandonia, J.-M. & Brenner, S. E. WebLogo: a sequence logo generator. Genome Res. 14, 1188–1190 (2004).

55. Klykov, O., Gangwar, S. P., Yelshanskaya, M. V., Yen, L. & Sobolevsky, A. I. Structure and desensitization of AMPA receptor complexes with type II TARP γ5 and GSG1L. Mol. Cell 81, 4771–4783.e7 (2021).

56. Meng, E. C. et al. UCSF ChimeraX: Tools for structure building and analysis. Protein Sci. 32, e4792 (2023).

57. Punjani, A., Rubinstein, J. L., Fleet, D. J. & Brubaker, M. A. cryoSPARC: algorithms for rapid unsupervised cryo-EM structure determination. Nat. Methods 14, 290–296 (2017).

58. Mastronarde, D. N. SerialEM: A program for automated tilt series acquisition on Tecnai microscopes using prediction of specimen position. Microsc. Microanal. 9, 1182–1183 (2003).

59. Carragher, B. et al. Leginon: an automated system for acquisition of images from vitreous ice specimens. J. Struct. Biol. 132, 33–45 (2000).

60. Scheres, S. H. W. RELION: implementation of a Bayesian approach to cryo-EM structure determination. J. Struct. Biol. 180, 519–530 (2012).

61. Baek, M. et al. Accurate prediction of protein structures and interactions using a three-track neural network. Science 373, 871–876 (2021).

62. Emsley, P., Lohkamp, B., Scott, W. G. & Cowtan, K. Features and development of Coot. Acta Crystallogr. D Biol. Crystallogr. 66, 486–501 (2010).

63. Liebschner, D. et al. Macromolecular structure determination using X-rays, neutrons and electrons: recent developments in Phenix. Acta Crystallogr D Struct Biol 75, 861–877 (2019).

64. Pettersen, E. F. et al. UCSF Chimera--a visualization system for exploratory research and analysis. J. Comput. Chem. 25, 1605–1612 (2004).

65. Davis, I. W. et al. MolProbity: all-atom contacts and structure validation for proteins and nucleic acids. Nucleic Acids Res. 35, W375–83 (2007).

66. Barad, B. A. et al. EMRinger: side chain–directed model and map validation for 3D cryo-electron microscopy. Nat. Methods 12, 943–946 (2015).

67. Prisant, M. G., Williams, C. J., Chen, V. B., Richardson, J. S. & Richardson, D. C. New tools in MolProbity validation: CaBLAM for CryoEM backbone, UnDowser to rethink ‘waters,’ and NGL Viewer to recapture online 3D graphics. Protein Sci. 29, 315–329 (2020).

68. Zheng, S. Q. et al. MotionCor2: anisotropic correction of beam-induced motion for improved cryo-electron microscopy. Nat. Methods 14, 331–332 (2017).

69. Waterhouse, A. M., Procter, J. B., Martin, D. M. A., Clamp, M. & Barton, G. J. Jalview Version 2—a multiple sequence alignment editor and analysis workbench. Bioinformatics 25, 1189–1191 (2009).

70. Langmead, B. & Salzberg, S. L. Fast gapped-read alignment with Bowtie 2. Nat. Methods 9, 357–359 (2012).

71. Li, H. et al. The Sequence Alignment/Map format and SAMtools. Bioinformatics 25, 2078–2079 (2009).

72. Masella, A. P., Bartram, A. K., Truszkowski, J. M., Brown, D. G. & Neufeld, J. D. PANDAseq: paired-end assembler for illumina sequences. BMC Bioinformatics 13, 31 (2012).

73. Martin, M. Cutadapt removes adapter sequences from high-throughput sequencing reads. EMBnet J. 17, 10 (2011).

74. MacRae, M. R. et al. Protein–protein interactions in the Mla lipid transport system probed by computational structure prediction and deep mutational scanning. J. Biol. Chem. 299, (2023).

75. Iglewicz, B. & Hoaglin, D. The ASQC Basic References in Quality Control: Statistical Techniques. vol. 16 9–17 (ASQC Quality Press, Milwaukee, 1993).

76. Zvelebil, M. J., Barton, G. J., Taylor, W. R. & Sternberg, M. J. Prediction of protein secondary structure and active sites using the alignment of homologous sequences. J. Mol. Biol. 195, 957–961 (1987).

77. Dewachter, L. et al. Deep mutational scanning of essential bacterial proteins can guide antibiotic development. Nat. Commun. 14, 241 (2023).

78. Baba, T. et al. Construction of Escherichia coli K-12 in-frame, single-gene knockout mutants: the Keio collection. Mol. Syst. Biol. 2, 2006.0008 (2006).

79. Cherepanov, P. P. & Wackernagel, W. Gene disruption in Escherichia coli: TcR and KmR cassettes with the option of Flp-catalyzed excision of the antibiotic-resistance determinant. Gene 158, 9–14 (1995).

80. Greenfield, E. A. Antibodies: A Laboratory Manual. (Cold Spring Harbor Laboratory Press, 2014).

81. Prince, C. & Jia, Z. An Unexpected Duo: Rubredoxin Binds Nine TPR Motifs to Form LapB, an Essential Regulator of Lipopolysaccharide Synthesis. Structure 23, 1500–1506 (2015).

82. Chin, J. W., Martin, A. B., King, D. S., Wang, L. & Schultz, P. G. Addition of a photocrosslinking amino acid to the genetic code of Escherichiacoli. Proc. Natl. Acad. Sci. U. S. A. 99, 11020–11024 (2002).

83. Studier, F. W. Protein production by auto-induction in high density shaking cultures. Protein Expr. Purif. 41, 207–234 (2005).

84. Mirdita, M. et al. ColabFold: making protein folding accessible to all. Nat. Methods 19, 679–682 (2022).

85. Del Alamo, D., Sala, D., Mchaourab, H. S. & Meiler, J. Sampling alternative conformational states of transporters and receptors with AlphaFold2. Elife 11, (2022).

86. Müllner, D. fastcluster: Fast Hierarchical, Agglomerative Clustering Routines for R and Python. Journal of Statistical Software 53, 1–18 (2013).

87. Olsson, M. H. M., Søndergaard, C. R., Rostkowski, M. & Jensen, J. H. PROPKA3: Consistent Treatment of Internal and Surface Residues in Empirical pKa Predictions. J. Chem. Theory Comput. 7, 525–537 (2011).

88. Søndergaard, C. R., Olsson, M. H. M., Rostkowski, M. & Jensen, J. H. Improved Treatment of Ligands and Coupling Effects in Empirical Calculation and Rationalization of pKa Values. J. Chem. Theory Comput. 7, 2284–2295 (2011).

89. Lomize, A. L., Todd, S. C. & Pogozheva, I. D. Spatial arrangement of proteins in planar and curved membranes by PPM 3.0. Protein Sci. 31, 209–220 (2022).

90. Jo, S., Kim, T., Iyer, V. G. & Im, W. CHARMM-GUI: a web-based graphical user interface for CHARMM. J. Comput. Chem. 29, 1859–1865 (2008).

91. Wu, E. L. et al. CHARMM-GUI Membrane Builder toward realistic biological membrane simulations. J. Comput. Chem. 35, 1997–2004 (2014).

92. Licari, G., Dehghani-Ghahnaviyeh, S. & Tajkhorshid, E. Membrane Mixer: A Toolkit for Efficient Shuffling of Lipids in Heterogeneous Biological Membranes. J. Chem. Inf. Model. 62, 986–996 (2022).

93. Phillips, J. C. et al. Scalable molecular dynamics on CPU and GPU architectures with NAMD. J. Chem. Phys. 153, 044130 (2020).

94. Huang, J. et al. CHARMM36m: an improved force field for folded and intrinsically disordered proteins. Nat. Methods 14, 71–73 (2017).

95. Klauda, J. B. et al. Update of the CHARMM all-atom additive force field for lipids: validation on six lipid types. J. Phys. Chem. B 114, 7830–7843 (2010).

96. Jorgensen, W. L., Chandrasekhar, J., Madura, J. D., Impey, R. W. & Klein, M. L. Comparison of simple potential functions for simulating liquid water. J. Chem. Phys. 79, 926–935 (1983).

97. Martyna, G. J., Tobias, D. J. & Klein, M. L. Constant pressure molecular dynamics algorithms. J. Chem. Phys. 101, 4177–4189 (1994).

98. Feller, S. E., Zhang, Y., Pastor, R. W. & Brooks, B. R. Constant pressure molecular dynamics simulation: The Langevin piston method. J. Chem. Phys. 103, 4613–4621 (1995).

99. Essmann, U. et al. A smooth particle mesh Ewald method. J. Chem. Phys. 103, 8577–8593 (1995).

100. Ryckaert, J.-P., Ciccotti, G. & Berendsen, H. J. C. Numerical integration of the cartesian equations of motion of a system with constraints: molecular dynamics of n-alkanes. J. Comput. Phys. 23, 327–341 (1977).

101. Miyamoto, S. & Kollman, P. A. Settle: An analytical version of the SHAKE and RATTLE algorithm for rigid water models. J. Comput. Chem. 13, 952–962 (1992).

102. Hopkins, C. W., Le Grand, S., Walker, R. C. & Roitberg, A. E. Long-Time-Step Molecular Dynamics through Hydrogen Mass Repartitioning. J. Chem. Theory Comput. 11, 1864–1874 (2015).

103. Balusek, C. et al. Accelerating Membrane Simulations with Hydrogen Mass Repartitioning. J. Chem. Theory Comput. 15, 4673–4686 (2019).

104. Li, Y., Liu, J. & Gumbart, J. C. Preparing Membrane Proteins for Simulation Using CHARMM-GUI. Methods Mol. Biol. 2302, 237–251 (2021).

105. Fiorin, G., Klein, M. L. & Hénin, J. Using collective variables to drive molecular dynamics simulations. Mol. Phys. 111, 3345–3362 (2013).

106. Powers, N. & Jensen, J. H. Chemically accurate protein structures: validation of protein NMR structures by comparison of measured and predicted pKa values. J. Biomol. NMR 35, 39–51 (2006).

107. Li, H., Robertson, A. D. & Jensen, J. H. Very fast empirical prediction and rationalization of protein pKa values. Proteins 61, 704–721 (2005).

108. Odenkirk, M. T., Zhang, G. & Marty, M. T. Do nanodisc assembly conditions affect natural lipid uptake? J. Am. Soc. Mass Spectrom. 34, 2006–2015 (2023).

109. Zhang, G. et al. Identifying membrane protein-lipid interactions with lipidomic lipid exchange-mass spectrometry. J. Am. Chem. Soc. 145, 20859–20867 (2023).

110. Tsugawa, H. et al. MS-DIAL: data-independent MS/MS deconvolution for comprehensive metabolome analysis. Nat. Methods 12, 523–526 (2015).

111. MacLean, B. et al. Skyline: an open source document editor for creating and analyzing targeted proteomics experiments. Bioinformatics 26, 966–968 (2010).

