## Extended Data for "LetA defines a structurally distinct transporter family involved in lipid trafficking"

### Extended Data Fig. 1

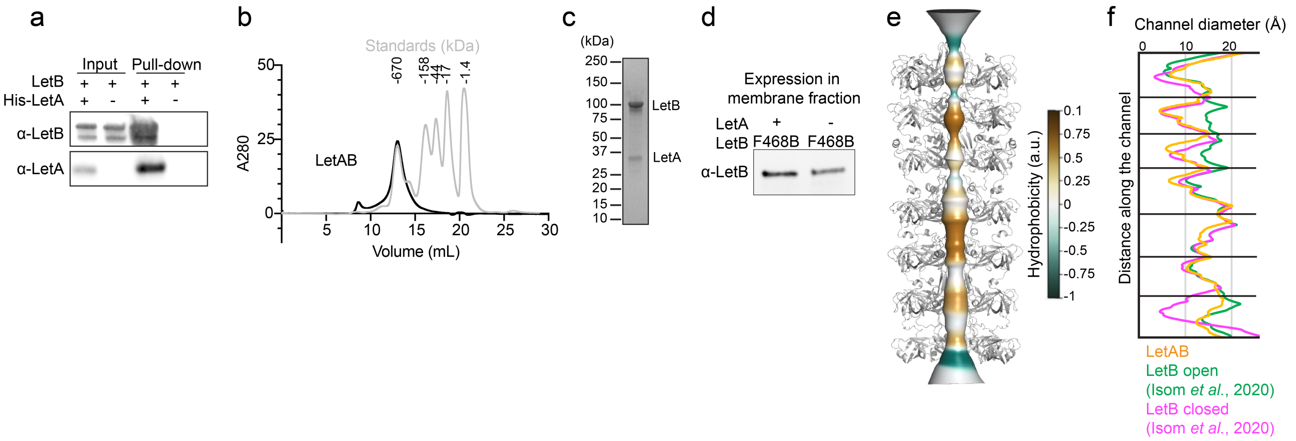

**Extended Data Fig. 1**. **Functional, biochemical and structural analysis of LetA, related Fig. 1. a,** Western blot showing results of pull-down assay, used to assess the interaction between LetA and LetB. His-LetA was used as the bait for pull-down, and interaction with untagged LetB was assessed using α-LetA (clone 72) and α-LetB antibodies (See Methods). **b,** Size exclusion chromatogram for LetAB (black) overlaid with standards (gray). The LetAB complex elutes around the same volume as the 670 kDa protein standard. **c,** Coomassie gel of purified LetAB from the peak fraction shown in (**b**). **d,** Western blot assessing expression and localization to the membrane fractions for LetB with BPA crosslinker incorporated at position 468, with or without co-expression of LetA. LetB presence in the membrane fraction was probed using α-LetB antibody (clone 72). **e,** Composite model of crosslinked LetAB (LetA omitted for clarity) showing the tunnel running through LetB. The tunnel is depicted as a smooth surface colored by the hydrophobicity of pore-facing residues, calculated using CHAP[52]. **f,** Tunnel radius of the LetAB and LetB in the open (PDB 6V0D) and closed (PDB 6V0C) states, measured using CHAP[52].

Extended Data Fig. 2

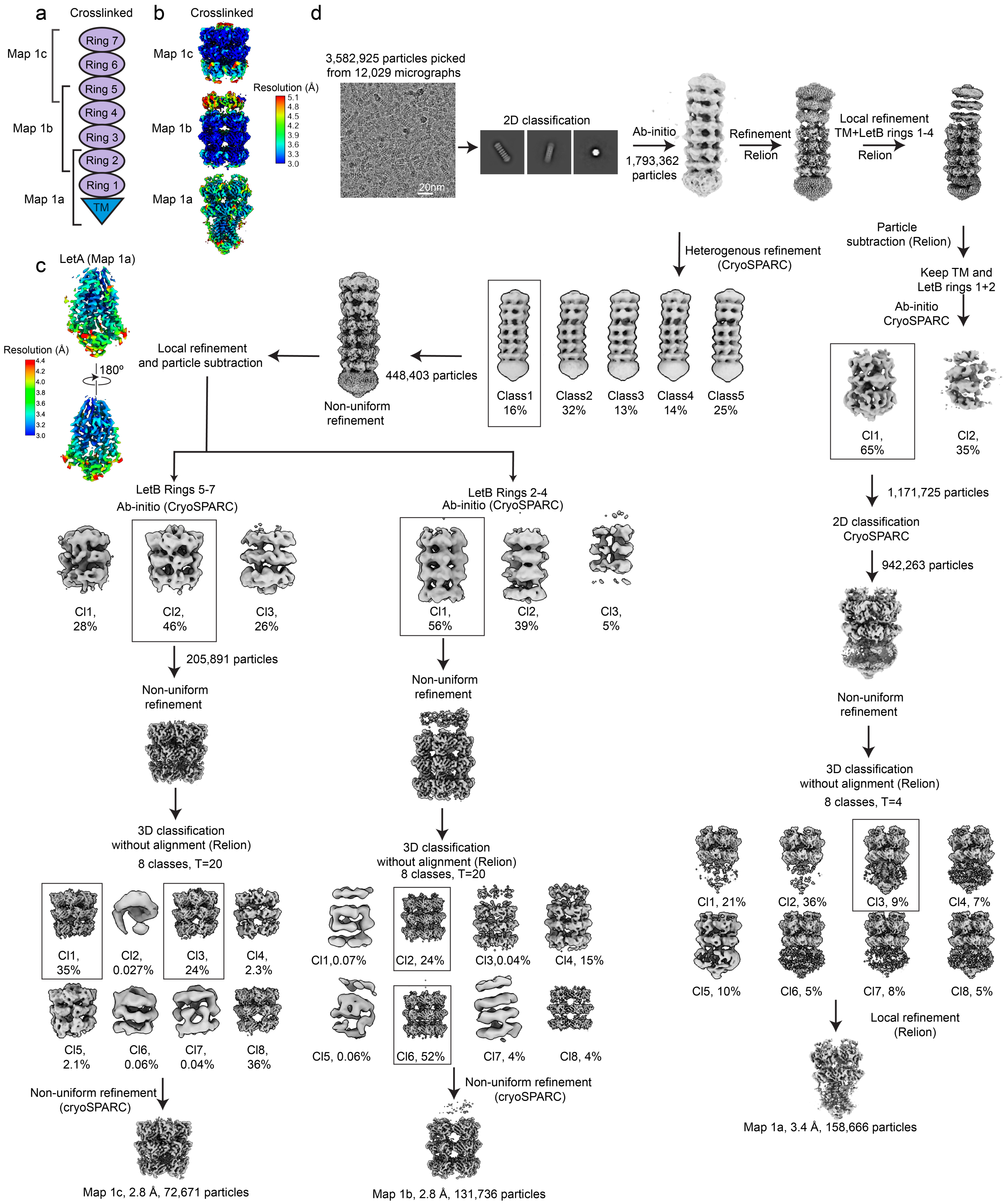

Extended Data Fig. 2

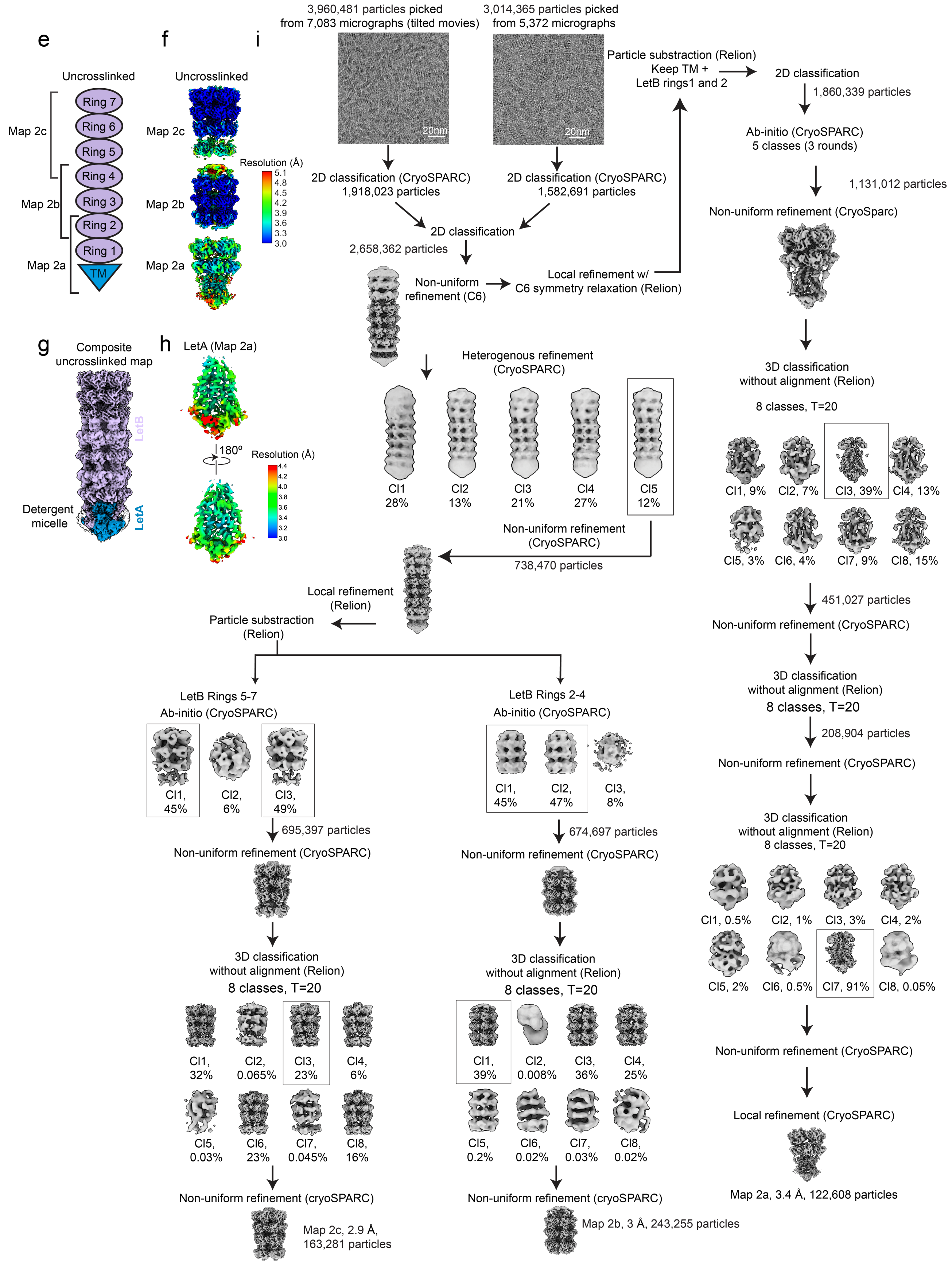

**Extended Data Fig. 2**. **Cryo-EM data processing workflow for the crosslinked and uncrosslinked datasets, related to Fig. 2.** Key steps of data processing workflows are shown for the crosslinked **a-d** and uncrosslinked (**e-i**) datasets. **a,** Schematic showing LetB (purple) in complex with LetA (blue, TM). The regions corresponding to each signal-subtracted map (crosslinked dataset) are annotated. **b,** Map 1a, 1b, and 1c, colored by local resolution as estimated using CryoSPARC. **c,** Map of crosslinked LetA (Map 1a), colored by local resolution as estimated using CryoSPARC. To calculate the local resolution of LetA, density for LetB Rings 1 and 2 are omitted. **d,** Cryo-EM data processing workflow for the crosslinked LetAB dataset; black boxes indicate the classes that were chosen for further processing. See Methods for details. **e,** Schematic showing LetB (purple) in complex with LetA (blue, TM). The regions corresponding to each signal-subtracted map (crosslinked dataset) are annotated. **f,** Map 2a, 2b, and 2c, colored by local resolution as estimated using CryoSPARC. **g,** Composite map (Map 2) of uncrosslinked LetAB. LetA (blue), LetB (purple) and the detergent micelle (white) are shown. The map is shown at a contour level of 2.24. **h,** Map of crosslinked LetA (Map 2a), colored by local resolution as estimated using CryoSPARC. To calculate the local resolution of LetA, density for LetB Rings 1 and 2 are omitted. **i,** Cryo-EM data processing workflow for the uncrosslinked LetAB dataset; black boxes indicate the classes that were chosen for further processing. See Methods for details.

### Extended Data Fig. 3

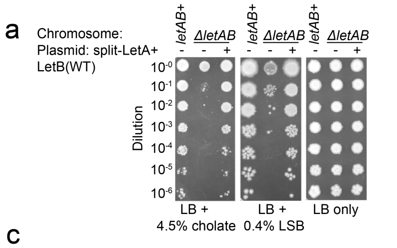


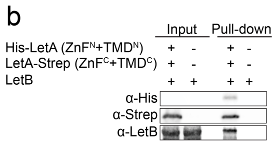


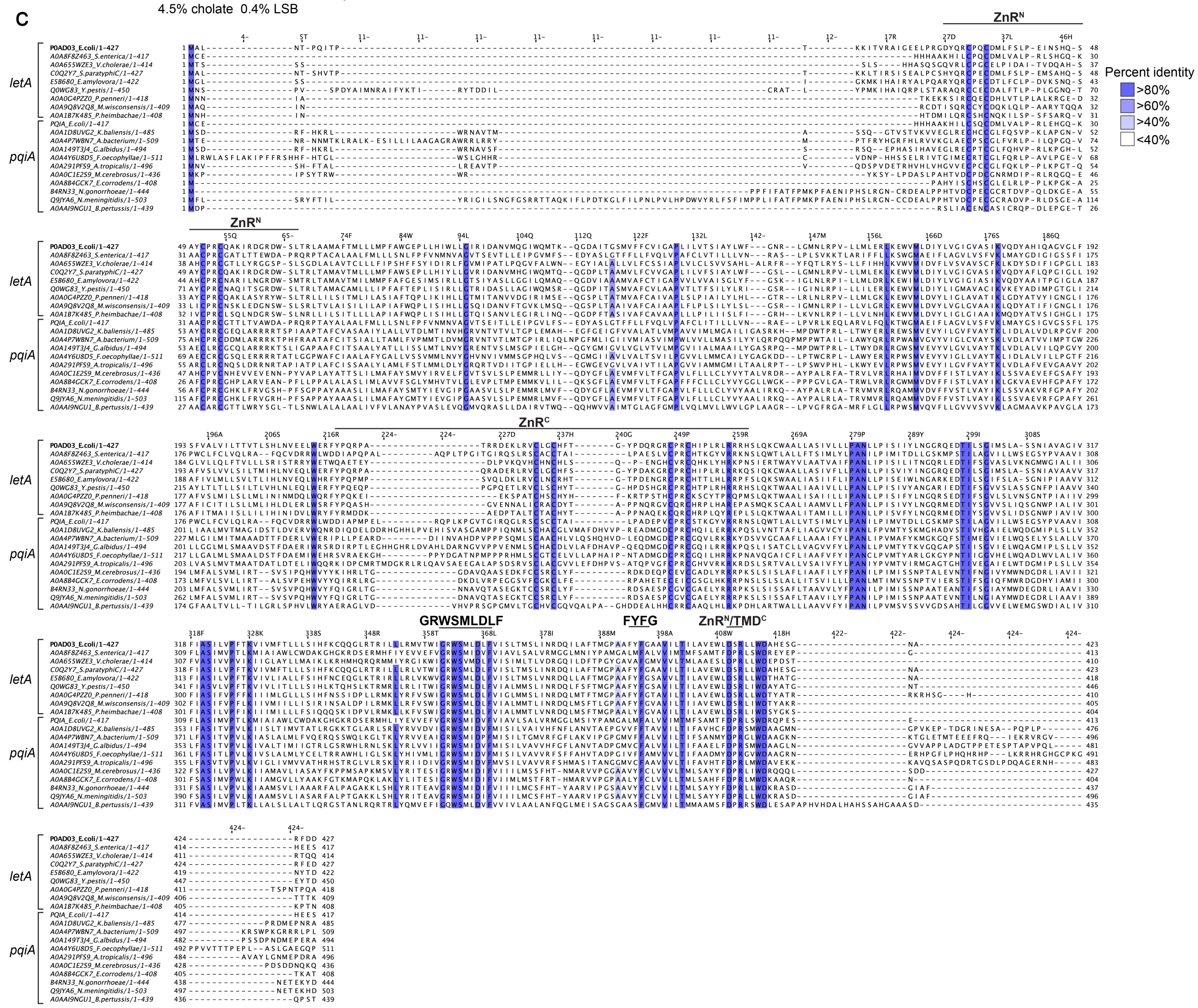


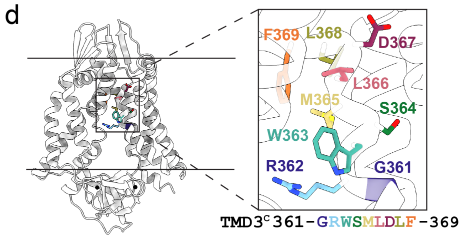


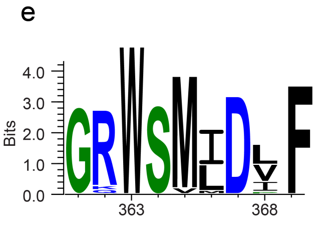


f

| Analyte | LetA (ppb) | Buffer (ppb) | Fold enrichment |
|---|---|---|---|
| Zn | 20 | 1 | 20 |
| Cu | 4 | 3 | 1 |
| Ni | 3 | 2 | 2 |
| Fe | 30 | 30 | 1 |
| Mn | 0.3 | 0.3 | 1 |
| Co | 0.2 | 0.1 | 2 |
| Cr | 200 | 100 | 2 |
| V | 10 | 10 | 1 |
| Ti | 100 | 100 | 1 |
| Ag | 0.0 | 0.04 | 0 |

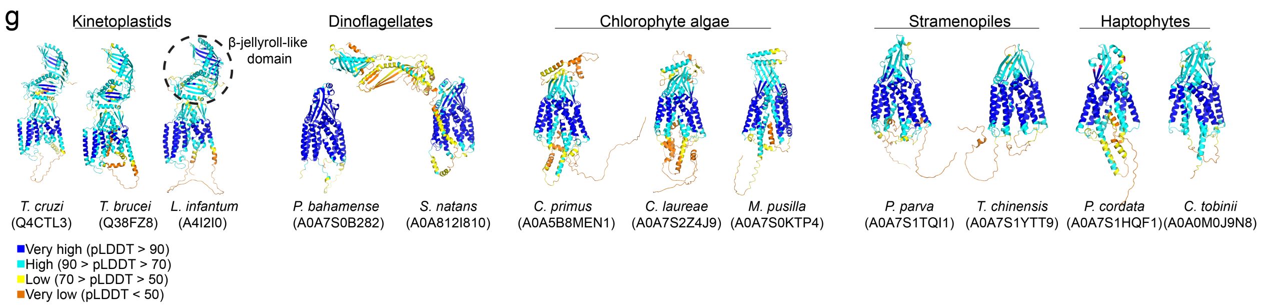

**Extended Data Fig. 3**. **Comparison of LetA with structural homologs, related to Fig. 3. a,** Cellular assay to assess the function of the LetA variant where N- and C-terminal molecules are co-expressed as separate open reading frames (split-LetA construct). 10-fold serial dilutions of strains indicated were spotted on LB agar with or without LSB or cholate, as noted. All strains are constructed in a *ΔpqiAB* background. **b,** Western blot showing results of pull-down assay, used to assess the interaction between the N-terminal LetA module and the C-terminal LetA module, when co-expressed as separate open reading frames (split-LetA construct). His-LetA($ZnR^N$+$TMD^N$) was used as the bait for pull-down, and interaction with LetA-Strep ($ZnR^C$+$TMD^C$) and untagged LetB was assessed using α-Strep and α-LetB antibodies. **c,** 20 sequences of LetA and PqiA proteins representing Alpha-, Beta, and Gammaproteobacteria, aligned using MUSCLE[53]; Uniprot IDs are provided. Positions are colored based on sequence identity (see color key). Regions of interest are indicated. **d,** Cartoon representation of LetA; inset shows residues that are part of the $^{361}$GRWSM-Ψ-D-Ψ-F$^{369}$ motif in colored sticks. **e,** Sequence logo of the $^{361}$GRWSM-Ψ-D-Ψ-F$^{369}$ motif generated using WebLogo 3[54]. Residues are colored based on hydrophilic (blue), neutral (green), or hydrophobic (black) properties. **f,** ICP-MS results showing semi-quantitative elemental scanning analysis for 10 transition metals. Average parts per billion (ppb) values of two independent experiments are shown for LetA and buffer control samples. **g,** AlphaFold models of LetA-like proteins from the various species as indicated. Uniprot IDs are provided in parentheses. Models are colored by the predicted local distance difference test (pLDDT) scores.

Extended Data Fig. 4

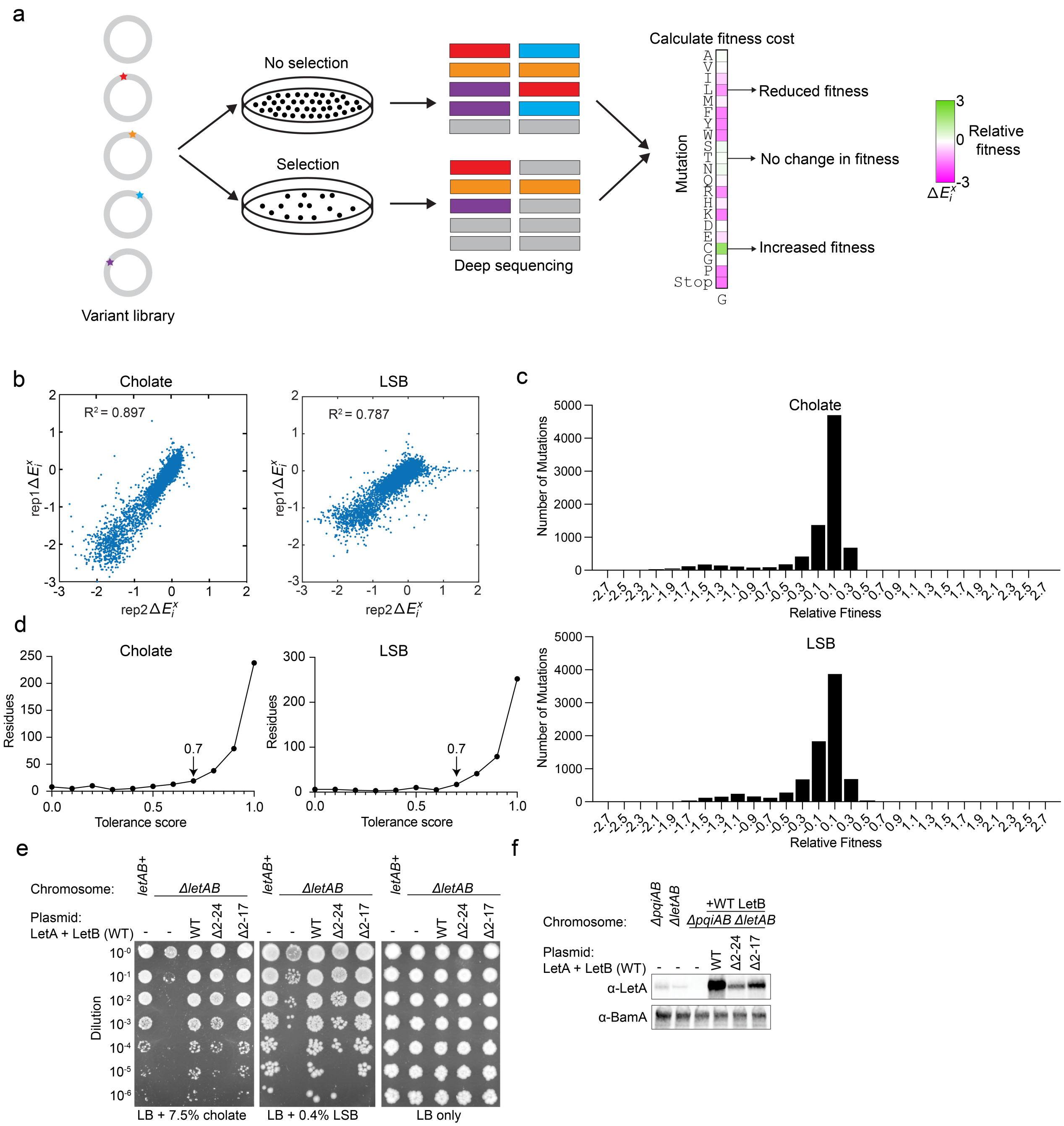

**Extended Data Fig. 4. DMS workflow and identification of functionally important residues, related to Fig. 4. a,** Schematic showing the workflow for the DMS experiment and data analysis. A variant library is generated and transformed into *ΔpqiAB ΔletAB* cells. Plasmids are shown as gray circles; star symbols indicate mutations. Cells are grown on LB plates in the absence or presence of detergent. Selection results in decreased frequency of mutations, which can be assessed by deep sequencing. The schematic of a vertical strip shows the relative fitness of each mutation for a given position, where mutations that decrease fitness relative to the WT are shown in shades of magenta, while mutations that increase fitness are in shades of green. *x*-axis: residue position; *y*-axis: mutation. **b,** Relative fitness values of replicate 1 (*y*-axis) and replicate 2 (*x*-axis). $R^2$ = 0.897 for cholate and $R^2$ = 0.787 for LSB, validating the reproducibility of the experiments. **c,** Histograms showing the frequency of the relative fitness scores for the cholate and LSB datasets. **d,** Distribution of tolerance scores for cholate and LSB datasets. The tolerance score of 0.7 was selected as the cut-off for determining functionally important residues in both datasets. **e,** Cellular assay to assess the function of LetA N-terminal truncation constructs. WT LetB is co-expressed with each of the LetA mutants. 10-fold serial dilutions of the strains indicated were spotted on LB agar with or without cholate or LSB. All strains are constructed in a *ΔpqiAB* background. **f,** Western blot analysis of cell lysates of the strains indicated, to compare cellular levels of WT LetA and N-terminal deletion mutants. α-LetA (clone 45) was used to probe LetA. BamA levels were probed using an α-BamA antibody as a loading control.

Extended Data Fig. 5

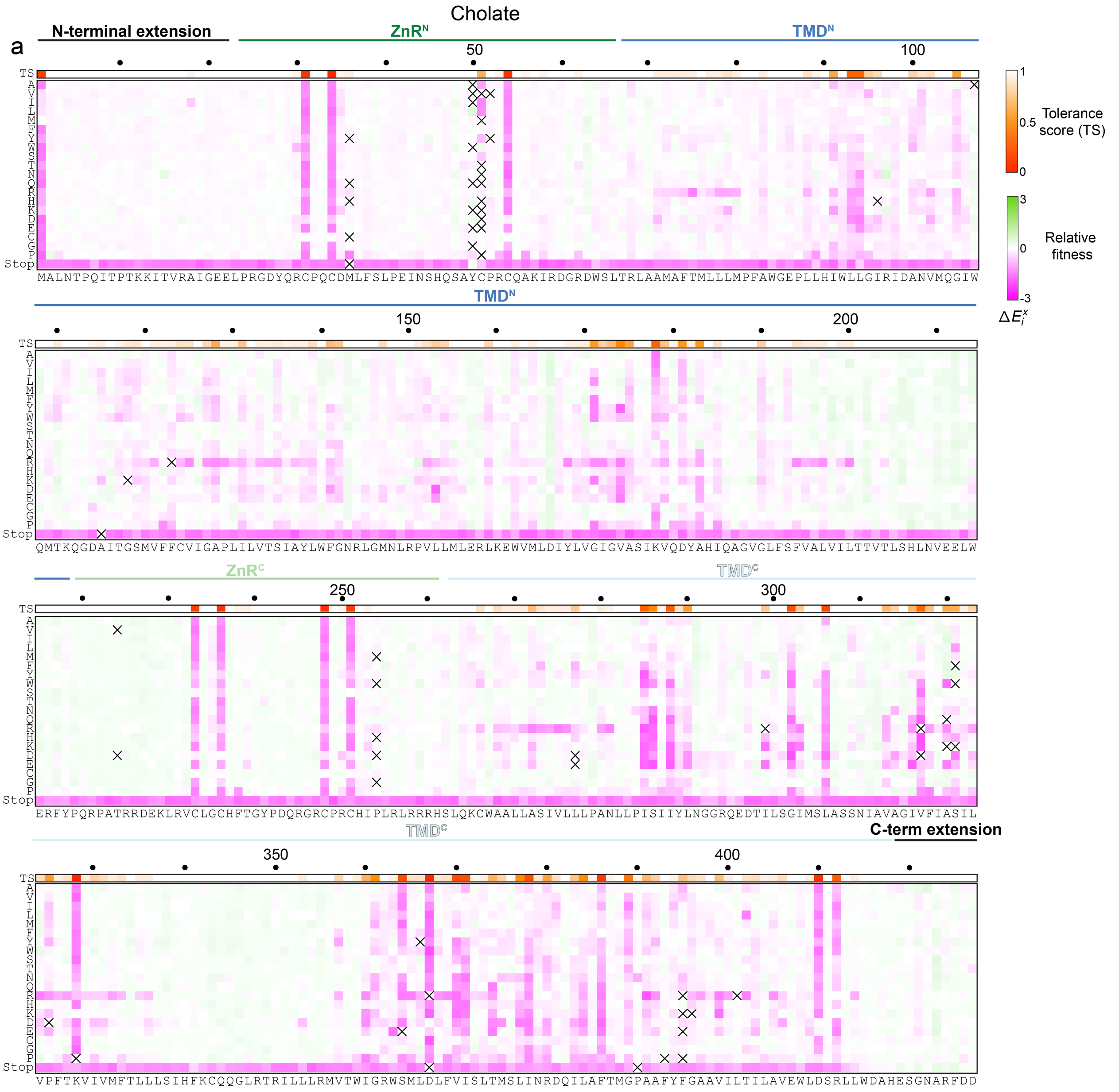

Extended Data Fig. 5

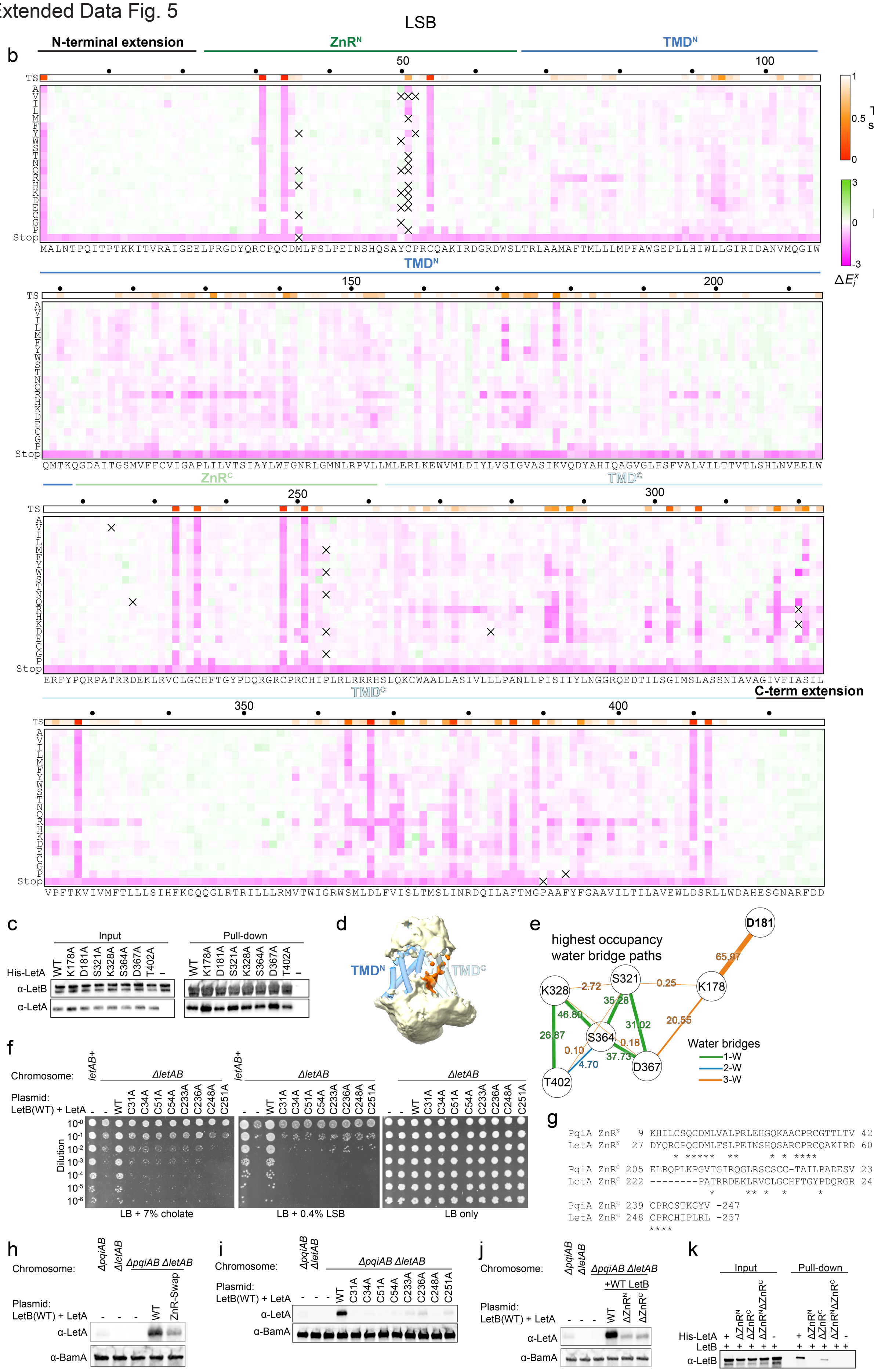

**Extended Data Fig. 5. Results of the DMS experiments, related to Fig. 4. a,b,** Heat maps summarizing results of deep mutational scanning of LetA (average of two biological replicates), where either cholate (**a**) or LSB (**b**) was used for selection. X-axis: the sequence of WT LetA from N-terminus to C-terminus; Y-axis: all possible amino acid substitutions, including the STOP codon. Each square represents the fitness cost of an individual mutation relative to WT. Mutations that decrease in fitness relative to the WT are shown in shades of magenta, while mutations that increase fitness are in shades of green, and white represents neutral mutations, as shown in the key. The tolerance score (TS) at each position is shown in the horizontal strip above the LetA sequence. Squares containing an "X" indicate incomplete coverage. **c,** Western blot analysis showing results of pull-down assay, used to assess the interaction between LetA polar network mutants and LetB. His-LetA WT and mutants were used as the bait for pull-down, and interaction with untagged LetB was assessed using α-LetA (clone 72) and α-LetB antibodies. LetA (WT or mutants) and WT LetB are co-expressed from the same plasmid. **d,** Water density map from equilibrium MD simulations for LetA. The map was generated using water oxygens within 3.5 Å of LetA and averaged over all the frames in the equilibrium MD simulations (-Lipid 1). LetA is shown in cartoon representation, where the helices are depicted as cylinders. $TMD^N$ and $TMD^C$ are colored according to Fig. 3a. The water density is colored in light yellow, except for the densities inside $TMD^C$, which are highlighted in orange. **e,** Hydrogen-bond network between proposed proton shuttle residues and water. The most frequently occurring water bridges in the equilibrium MD simulations (-Lipid 1, three replicas) are shown. Each line represents the water bridge that most consistently appears between the two corresponding nodes throughout the simulation. Each polar residue is shown as a node, with the lines connecting the nodes representing water bridges. The thickness of the lines corresponds to the relative occupancy of the water bridge throughout the simulation. The color coding of the edges indicates the number of water molecules involved in forming the bridge: 1-W (green) represents a single water molecule bridge, 2-W (blue) indicates a bridge formed through two water molecules, and 3-W (orange) shows a water bridge through three water molecules. The percentage occupancy of each water bridge is annotated alongside the edges. **f,** Cellular assay to examine the function of LetA ZnR cysteine mutants. 10-fold serial dilutions of the indicated strains were spotted on LB agar with or without cholate or LSB. All strains are constructed in a *ΔpqiAB* background. **g,** Sequence alignment of the LetA and PqiA ZnR domains. **h,** Western blot analysis of lysates of the strains indicated, to compare cellular levels of WT LetA and a ZnR mutant in which the ZnR domains in LetA are swapped with those of the *E. coli* PqiA protein. α-LetA (clone 72) was used to probe LetA. BamA levels were probed using an α-BamA antibody as a loading control. **i,** Western blot analysis of lysates of the strains indicated, to compare cellular levels of WT LetA and ZnR cysteine point mutants. α-LetA (clone 72) was used to probe LetA. BamA levels were probed using an α-BamA antibody as a loading control. **j,** Western blot analysis of lysates of the strains indicated, to compare cellular levels of WT LetA and ZnR deletion mutants. Anti-LetA (clone 72) was used to probe LetA. BamA levels were probed using an anti-BamA antibody as a loading control. **k,** Western blot analysis showing results of pull-down assay, used to assess the interaction between LetA ZnR deletion mutants and LetB. His-LetA WT and mutants were used as the bait for pull-down, and interaction with untagged LetB was assessed using α-LetA (clone 72) and α-LetB antibodies.

### Extended Data Fig. 6

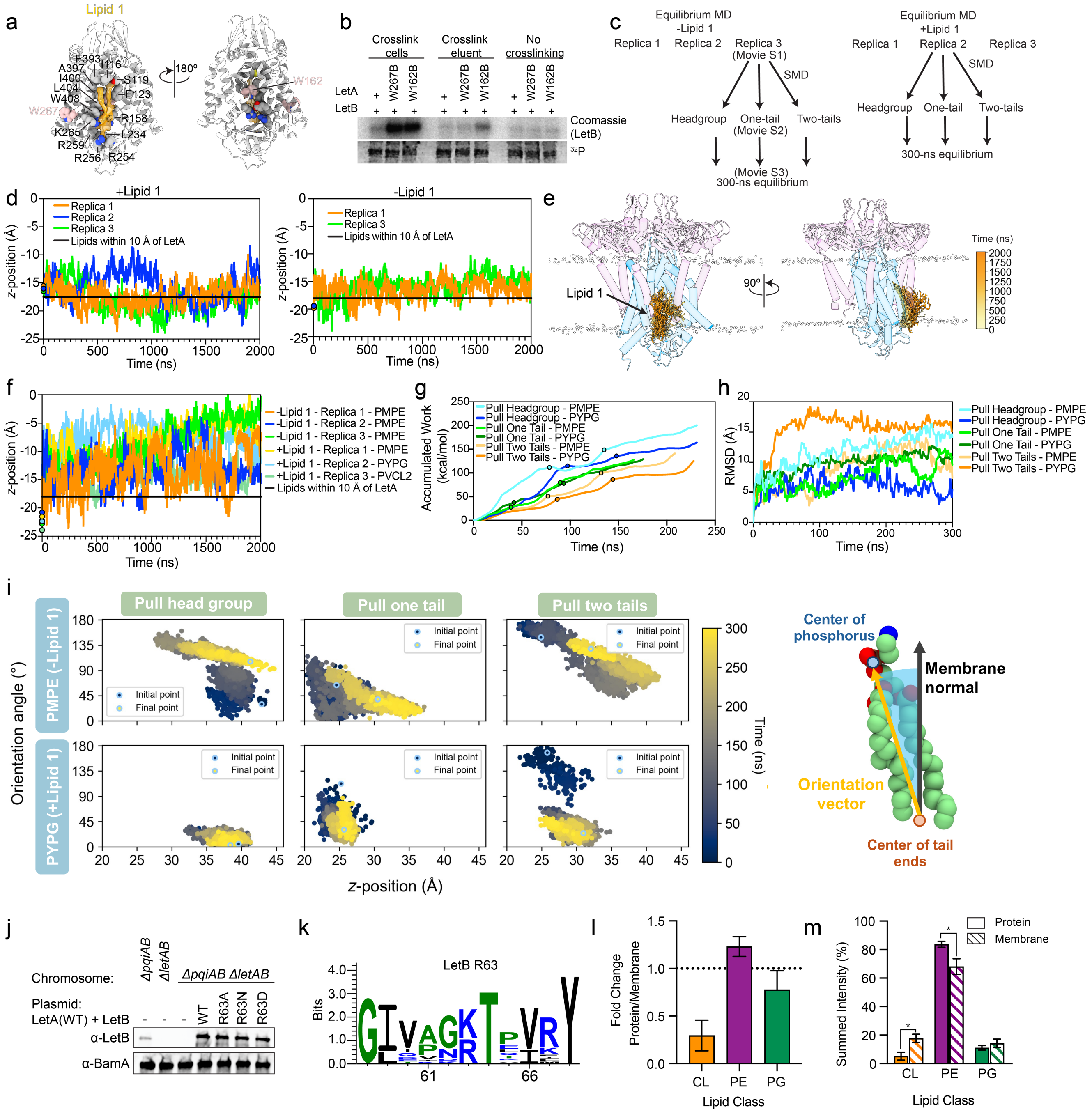

**Extended Data Fig. 6. MD simulations to examine putative lipid translocation pathways through LetA, related to Fig. 5. a,** Cartoon representation of LetA (gray) with EM density (orange) corresponding to Lipid 1 (yellow). Fig. was prepared in ChimeraX[56] using the zone function with an applied radius of 3.5 Å and map contour of 3.24 (Map 2a). Gray spheres indicate the residues that are interacting with Lipid 1. Nitrogen and oxygen atoms are highlighted in blue and red, respectively. Residues replaced with BPA in crosslinking experiment presented in (b) are shown as pink spheres (W162 and W267). W267 is exposed to bulk lipids and serves a positive control. W127 is predicted to interact with one of the Lipid 1 tails. **b,** SDS-PAGE analysis of purified LetAB and its BPA mutants, either crosslinked or uncrosslinked and stained by Coomassie (LetB) or phosphor-imaged ($^{32}$P signal). **c,** Diagram showing the MD simulations performed and the movies associated with each step. **d,** Time series of the *z*-position of the phosphorus atom of Lipid 1 in each replica. Lipid 1 was either included (+Lipid 1) or excluded (-Lipid 1) at the start of the simulation. Except in Replica 2 (-Lipid 1), a phospholipid binds to the Lipid 1 binding site regardless of whether a lipid was modeled in it to begin with or not (there is no trace for Replica 2 (-Lipid 1) in the right panel, since no lipid is stably bound). The starting *z*-positions of the phosphorus atom are marked by circles (see color key). The average phosphorous atom position from the lipids within 10 Å of LetA is shown as a black line. **e,** Representative snapshots of Lipid 1 in Replica 1 (+Lipid 1) of the equilibrium MD simulation, showing its stable binding throughout the simulation. Each Lipid 1 snapshot is colored according to the simulation timestep, transitioning from light yellow at the start to orange at the end. LetA is shown in blue, MCE Ring 1 in purple, and phosphorus atoms of the bulk lipids are in white spheres to indicate the position of the membrane. Helices are depicted as cylinders. **f,** Time series of the *z*-position of the phosphorus atom from the most elevated lipid in the central cavity. The lipid type is indicated in the color key. The starting *z*-position of the phosphorus atoms are indicated by a circle. The average phosphorus atom position of lipids within 10 Å of LetA is depicted as a black line. **g,** Accumulated non-equilibrium work profiles for different pulling protocols and the pulled lipid types (PMPE and PYPG). The first and second circles on each line indicate the time when the pulled lipid reached the bottom and middle of the periplasmic pocket, respectively. At the end of each line, the lipid has reached the top of the periplasmic pocket. **h,** RMSD profiles of steered lipids. RMSD plots for each lipid type (PMPE and PYPG) and different pulling methods (head group, one tail, or two tails) showing lipid movement and flexibility within the periplasmic pocket during the 300-ns equilibrium simulations after SMD. **i,** The conformational dynamics of the steered lipids are illustrated in 2D scatter plots showing the orientation angle versus the *z*-position of the phosphate group. Time progression is indicated by a color gradient from dark blue to yellow, with the initial and final frame highlighted by a circle (see color key). The orientation angle, as shown in the schematic on the right, is defined by the angle between the orientation vector (yellow) and the membrane normal (black). **j,** Western blot analysis of cell lysates of the strains indicated, to compare cellular levels of WT LetB and LetB R63 mutants. LetB was probed using a α-LetB antibody. As a loading control, BamA levels were detected using an α-BamA antibody. WT LetA was co-expressed with each LetB mutant. **k,** Sequence logo of positions 58 to 68 of LetB by WebLogo 3 (see Methods for list of LetB sequences). **l,** Bar plot showing the fold change differences between purified LetAB and the membrane fraction. Black dotted line indicates no enrichment or depletion of the indicated lipid class. Data are mean ± standard deviation from three independent LetAB purifications. **m,** Bar plot showing the relative abundance of phospholipids from the membrane fraction of *E. coli* cells overexpressing LetAB and purified LetAB. The summed intensity is used to estimate abundance. Data are mean ± standard deviation from three independent LetAB purifications. Asterisk indicates the p-value is less than 0.05.

### Extended Data Fig. 7

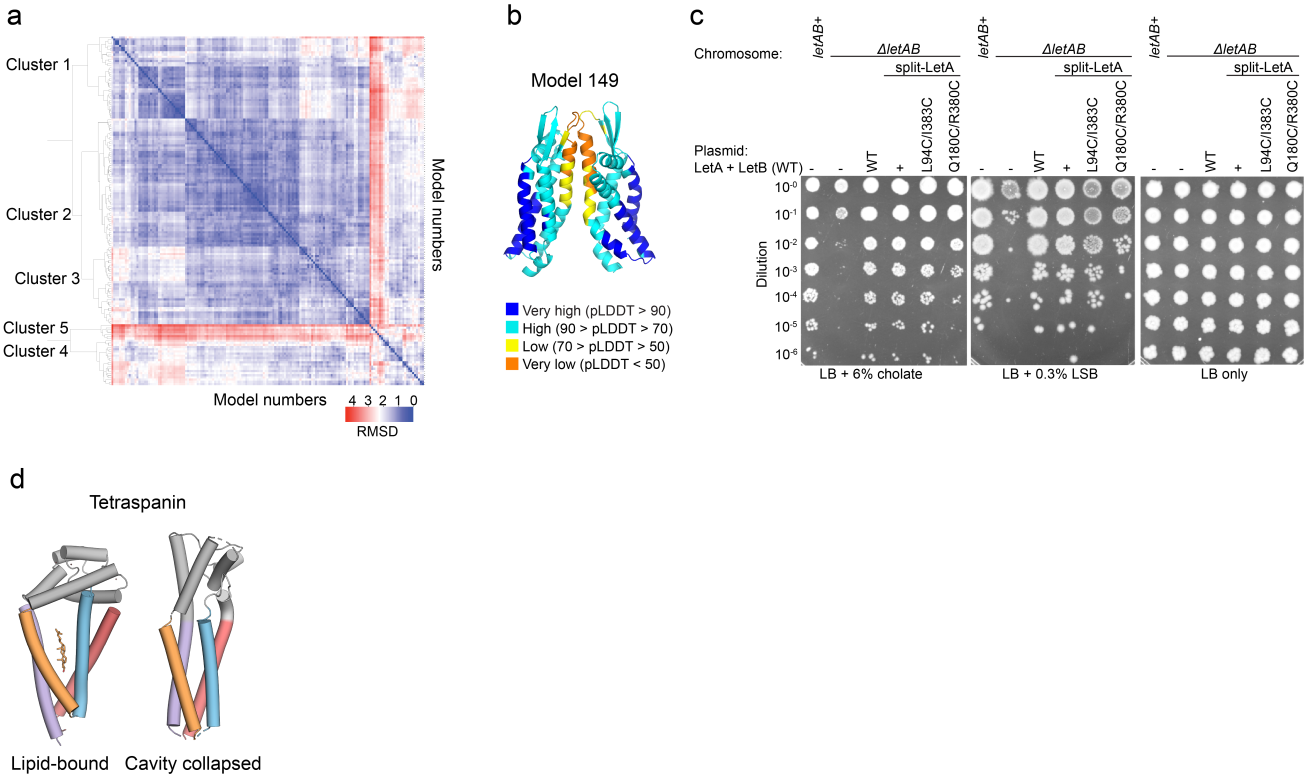

**Extended Data Fig. 7. LetA alternative conformations and evolutionary comparisons, related to Fig. 5. a,** Heatmap of pairwise RMSD values for AlphaFold2 LetA predictions (see Methods). Hierarchical clustering analysis was performed, resulting in five clusters. **b,** Cartoon representation of Model 149 (Cluster 5) colored by pLDDT score. ZnR domains are not shown due to unreliable predictions (see Methods). **c,** Cellular assay to examine the function of split-LetA mutants. 10-fold serial dilutions of strains indicated were spotted on LB agar with or without cholate or LSB. All strains are constructed in a *ΔpqiAB* background. **d,** Cartoon representation of tetraspanin in the absence (PDB: 7JIC) or presence of cholesterol (PDB: 5TCX). In the absence of cholesterol, the lipid-binding cavity collapses. Structural elements that are similar to those in the LetA TMD are colored according to Fig. 3c.
