## Supplementary Figure 1 for "LetA defines a structurally distinct transporter family involved in lipid trafficking"

#### a) Related to Extended Data Fig.1a

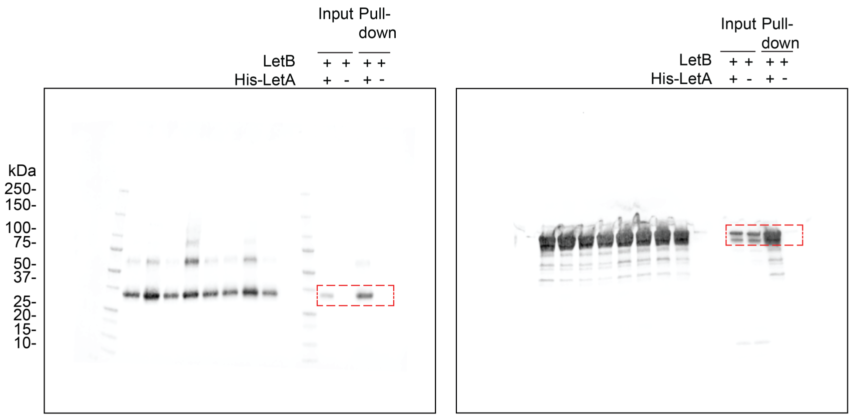


α-LetA (clone 72, 700 channel) α-LetB (800 channel)

#### b) Related to Extended Data Fig.1c

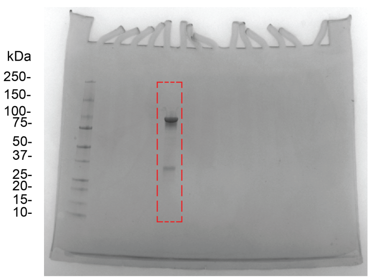


#### c) Related to Fig. 1e

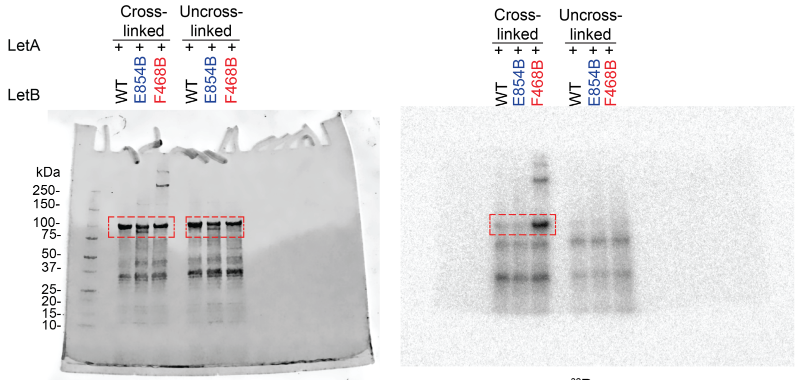


Coomassie $^{32}P$

#### d) Related to Fig. 1f

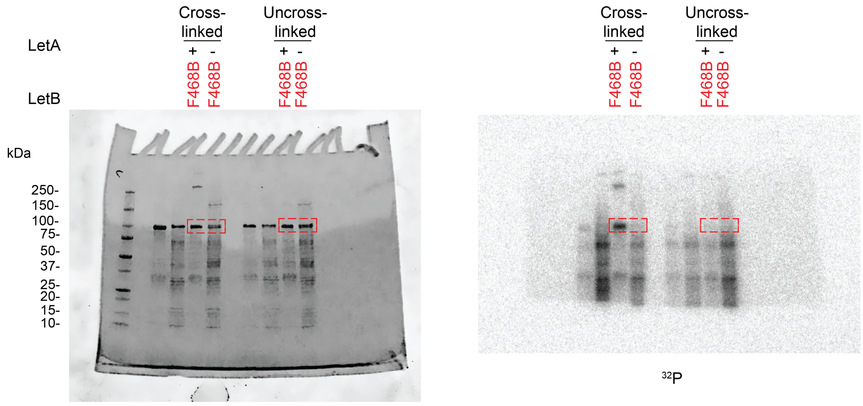


Coomassie $^{32}P$

#### Supplementary Figure 1

##### e) Related to Extended Data Fig. 1d

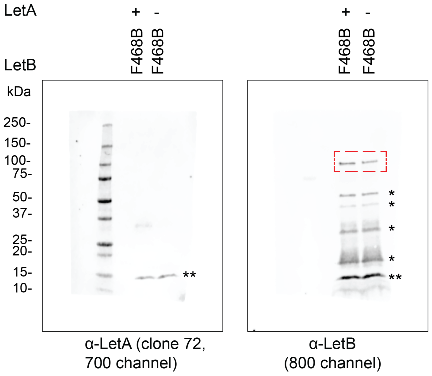


##### f) Related to Extended Data Fig. 3b

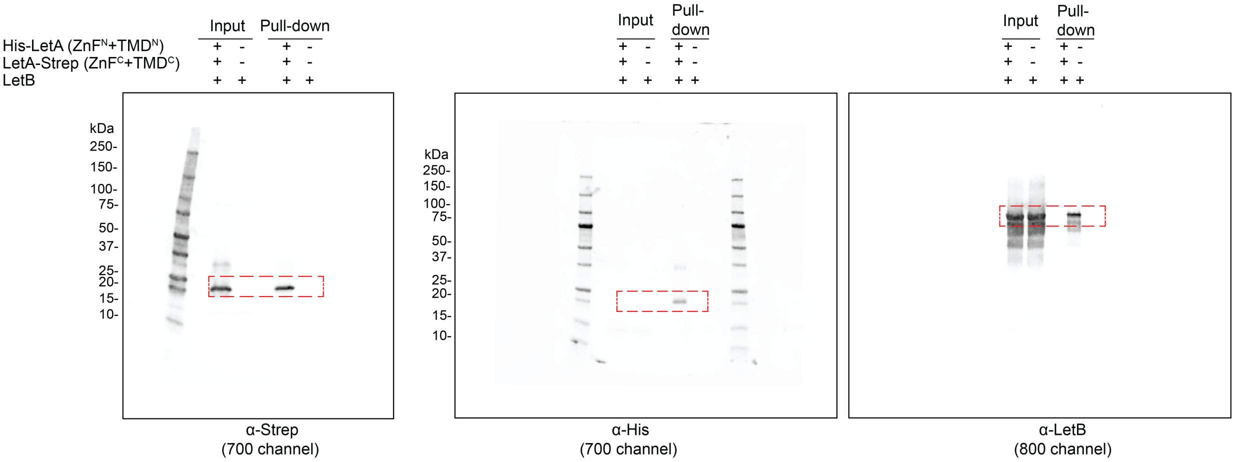


##### g) Related to Extended Data Fig. 4f

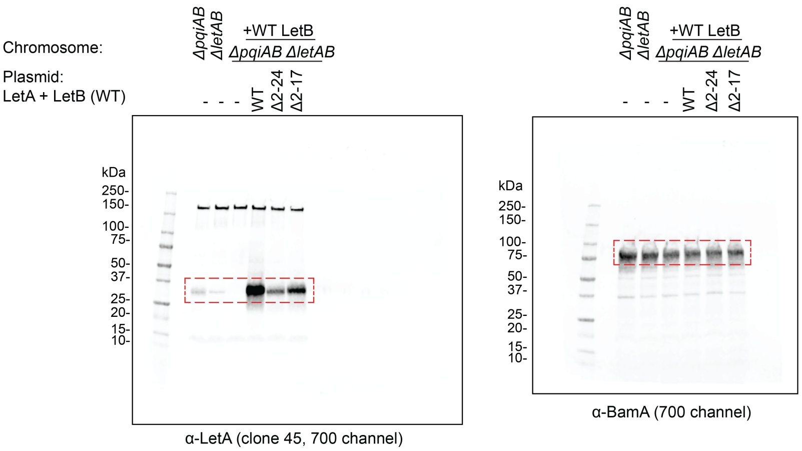

#### Supplementary Figure 1

##### h) Related to Extended Data Fig. 5c

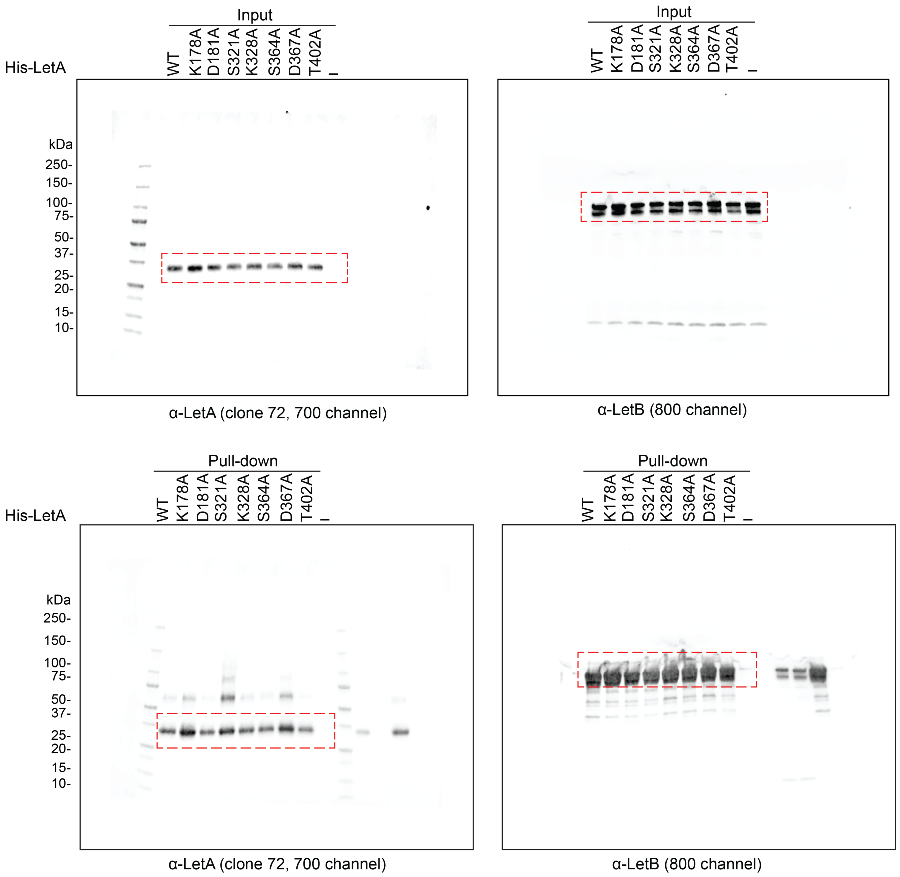


##### i) Related to Extended Data Fig. 5h

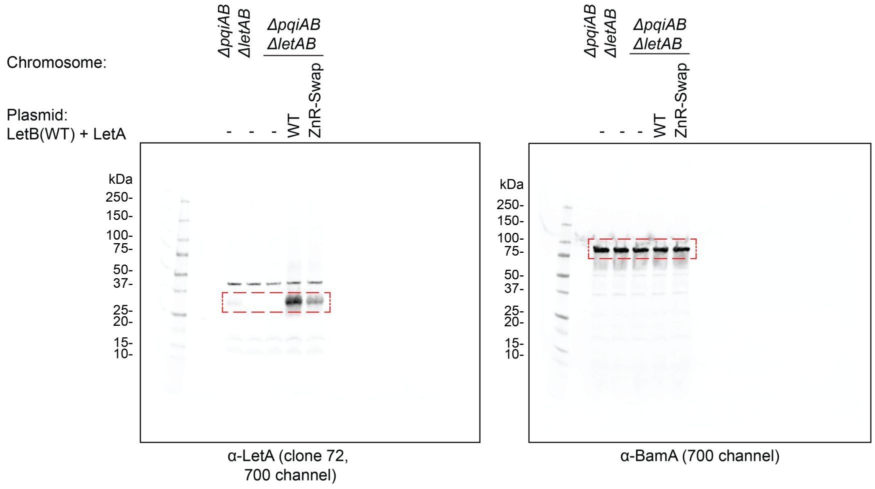

### Supplementary Figure 1

#### j) Related to Extended Data Fig. 5i

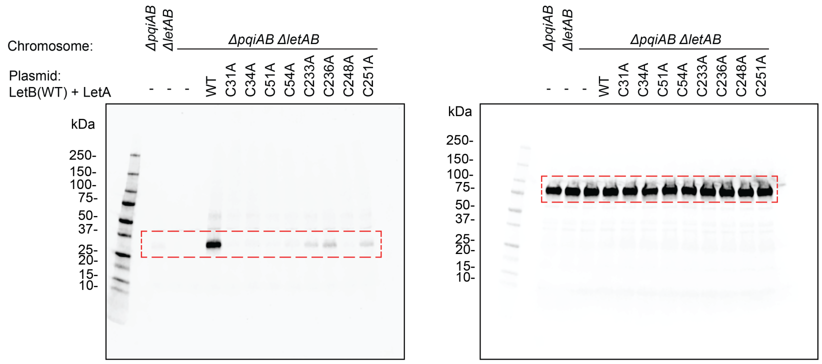


α-LetA (clone 72, 700 channel)

α-BamA (700 channel)

#### k) Related to Extended Data Fig. 5j

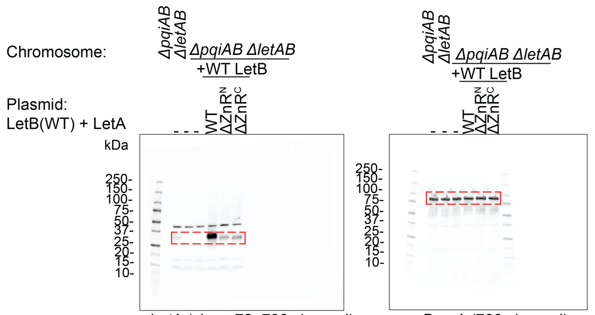


α-LetA (clone 72, 700 channel)

α-BamA (700 channel)

#### l) Related to Extended Data Fig. 5k

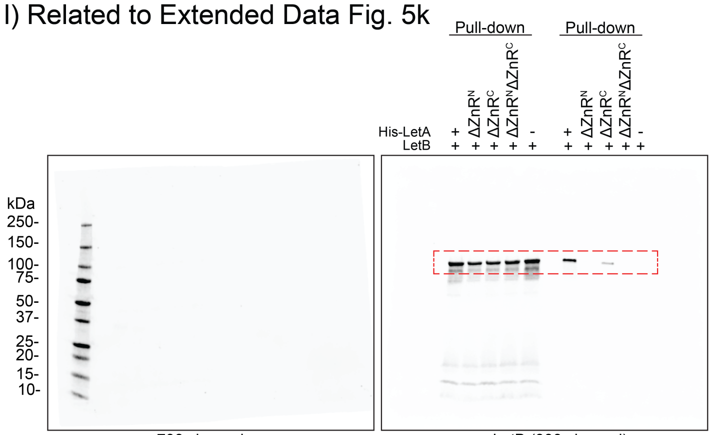


700 channel

α-LetB (800 channel)

#### m) Related to Extended Data Fig. 6b

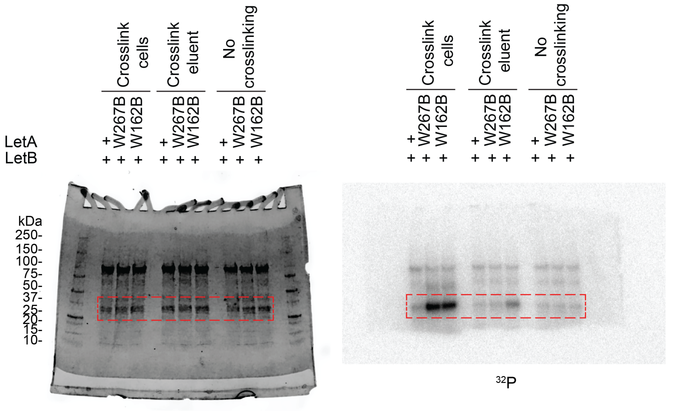


$^{32}P$

Coomassie

#### Supplementary Figure 1

##### n) Related to Extended Data Fig. 6j

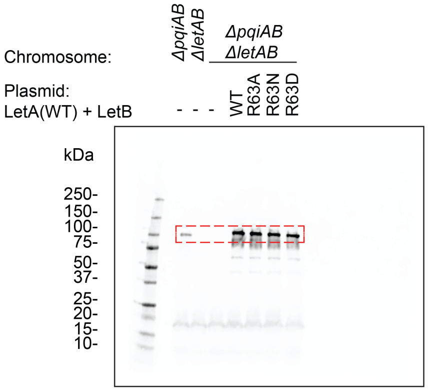


α-LetB (700 channel)

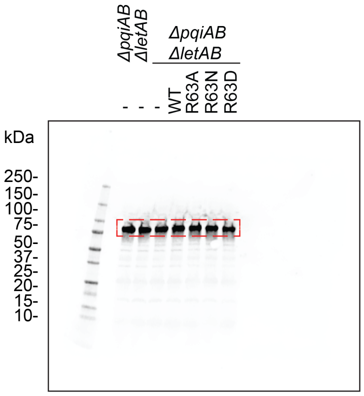


α-BamA (700 channel)

##### o) Related to Fig. 5k

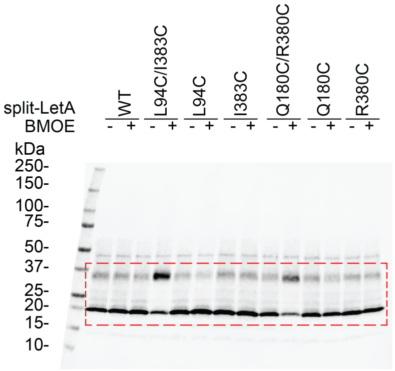


α-LetA (clone 72, 700 channel)

**Supplementary Figure 1. Uncropped gels and blots, shown in Main and Extended Data Figs.** Each panel shows the uncropped version of gels or western blots corresponding to those shown in the main figures or Extended Data Figs., as indicated. Dashed boxes (red) indicate where the images were cropped. Molecular weight protein standards are shown, and the antibodies used in the experiment are indicated, along with the LI-COR channels. **a,** Uncropped western blots shown in Extended Data Fig. 1a. **b,** Uncropped Coomasie gel shown in Extended Data Fig. 1c. **c,** Uncropped Coomassie gel (left) and phosphor screen (right) shown in Fig. 1e. Before exposing the gel to the phosphor screen, the gel was cut to remove the section below the 20 kDa marker, which contains free phospholipids that result in a high $^{32}P$ background. **d,** Uncropped Coomassie gel (left) and phosphor screen (right) shown in Fig. 1f. Before exposing the gel to the phosphor screen, the gel was cut to remove the section below the 20 kDa marker, which contains free phospholipids that result in a high $^{32}P$ background. **e,** Uncropped Western blots shown in Extended Data Fig. 1d. We have previously shown specificity for the custom antibody used for LetB[31,34]. As BPA incorporation is not 100% efficient, in addition to full length LetB, LetB truncation products may also be generated, indicated by asterisks. The bands annotated with double asterisks are due to non-specific antibody reactivity with lysozyme, which was included in the bacterial lysis buffer. **f,** Uncropped Western blots shown in Extended Data Fig. 3f. The α-His and α-LetB images are from the same blot and the α-Strep image is from a separate blot analyzing the exact same samples. **g,** Uncropped Western blot corresponding to Extended Data Fig. 4f. The α-LetA (clone 45) and α-BamA images are from separate blots analyzing the exact same samples. **h,** Uncropped Western blots shown in Extended Data Fig. 5c. **i,** Uncropped Western blots shown in Extended Data Fig. 5h. The α-LetA (clone 72) and α-BamA are from separate blots analyzing the exact same samples. **j,** Uncropped Western blot shown in Extended Data Fig. 5i. The α-LetA (clone 72) and α-BamA images are from separate blots analyzing the exact same samples. **k,** Western blot related to Extended Data Fig. 5j. The α-LetA (clone 72) and α-BamA are from separate blots analyzing the exact same samples. **l,** Uncropped Western blot shown in Extended Data Fig. 5k. The image from the 700 channel is provided to show the protein ladder. **m,** Uncropped Coomassie gel (left) and phosphor screen (right) shown in Extended Data Fig. 6b. Before exposing the gel to the phosphor screen, the gel was cut to remove the section below the 20 kDa marker, which contains free phospholipids that result in a high $^{32}P$ background. **n,** Uncropped Western blot shown in Extended Data Fig. 6j. The α-LetB and α-BamA images are from separate blots analyzing the exact same samples. **o,** Uncropped Western blot shown in Fig. 5k. The monomer (~20 kDa) and dimer (~35 kDa) products were probed using an α-LetA antibody (clone 72).
