## Supplementary Table 1 for "LetA defines a structurally distinct transporter family involved in lipid trafficking"

**Table S1A. Cryo-EM data collection, refinement, and validation statistics.**

| **Data collection and processing** | Map 1a (LetA+rings1-2) |
|---|---|
| EMD ID | EMD-49148 |
| Microscope | Krios G3 (PNCC Krios-1) |
| Voltage (kEV) | 300 |
| Camera | K3 |
| Energy filter | BioContinuum |
| Magnification | 81000x |
| Nominal Pixel size (Å/pixel) | 1.029 |
| Total electron exposure (e-/Å$^2$) | 50 |
| Exposure time (s) | 2.8 |
| Number of frames (no.) | 50 |
| Defocus range (μm) | -0.8 to -2.1 |
| Estimated defocus range (μm) | -0.1 to -4.0 |
| Automation software | Serial-EM |
| No. of micrographs at 0° tilt | 12029 |
| Number of particles | 158666 |
| Resolution (Å, FSC=0.143) | 3.38 |
| Map sharpening B-factors (Å$^2$) | -20 |
| Sphericity from 3D FSC | 0.966 |
| Symmetry | C1 |
| Box size (px) | 256 |
| **Model Composition** | Composite Model in Map 1 |
| EMD ID | EMD-49148 |
| PDB ID | 9N8W |
| Initial model (PDB code) | RoseTTAFold, 6V0J, 6V0F, 6V0E |
| Model composition | |
| Chains | 8 |
| Non-hydrogen atoms | 41785 |
| Protein residues | 5478 |
| Ligands | 2 zinc |
| Mean B-factor (Å$^2$) | |
| Protein | 81 |
| Ligand | 88 |
| R.M.S. deviations | |
| rmsd (bonds) | 0.003 |
| rmsd (angles) | 0.571 |
| Validation | |
| EMRinger score | 2.36 |
| MolProbity score | 1.45 |
| Clashscore, all atoms | 4.65 |
| Rotamer outliers (%) | 0.84 |
| CaBLAM outliers (%) | 3.06 |
| Cβ outliers (%) | 0 |
| FSC (Model-map) (Å, FSC=0.5) | 3.3 |
| Ramachandran plot (%): | |
| Favored | 96.58 |
| Allowed | 3.37 |
| Outliers | 0.05 |
| Rama-Z | |
| whole | -0.52 |
| helix | -0.60 |
| sheet | 0.50 |
| loop | -0.78 |
| Map CC (mask) | 0.80 |
| Map CC (box) | 0.75 |
| Map CC (peaks) | 0.66 |
| Map CC (volume) | 0.79 |
| Map CC for ligands | 0.81 |

**Table S1B. Cryo-EM data collection, refinement, and validation statistics.**

| **Data collection and processing** | Map 2a (LetA+rings1-2) |
|---|---|
| EMD ID | |
| Microscope | Krios G3 (NYSBC Krios #1) |
| Voltage (kEV) | 300 |
| Camera | K3 |
| Energy filter | BioContinuum |
| Magnification | 81000x |
| Nominal Pixel size (Å/pixel) | 1.083 |
| Total electron exposure (e-/Å$^2$) | 51 |
| Exposure time (s) | 2 |
| Number of frames (no.) | 40 |
| Defocus range (μm) | -2 to -5 |
| Estimated defocus range (μm) | -1.3 to -3.6 |
| Automation software | Leginon |
| No. of micrographs at 0° tilt | 5372 |
| No. of micrographs at -30° tilt | 7083 |
| Number of particles | 190823 |
| Resolution (Å, FSC=0.143) | 3.37 |
| Map sharpening B-factors (Å2) | -10 |
| Sphericity from 3D FSC | 0.981 |
| Symmetry | C1 |
| Box size (px) | 256 |
| **Model Composition** | Composite Model in Map 2 |
| EMD ID | EMD-49152 |
| PDB ID | 9N8X |
| Initial model | Crosslinked LetAB model |
| Model composition | |
| Chains | 9 |
| Non-hydrogen atoms | 41543 |
| Protein residues | 5441 |
| Ligands | 1 PEF/2 zinc |
| Mean B-factor (Å$^2$) | |
| Protein | 105 |
| Ligand | 97 |
| R.M.S. deviations | |
| rmsd (bonds) | 0.002 |
| rmsd (angles) | 0.543 |
| Validation | |
| EMRinger score | 2.32 |
| MolProbity score | 1.46 |
| Clashscore, all atoms | 3.86 |
| Rotamer outliers (%) | 1.18 |
| CaBLAM outliers (%) | 3.32 |
| Cβ outliers (%) | 0 |
| FSC (Model-map) (Å, FSC=0.5) | 3.2 |
| Ramachandran plot (%): | |
| Favored | 96.49 |
| Allowed | 3.43 |
| Outliers | 0.07 |
| Rama-Z | |
| whole | -0.93 |
| helix | -0.85 |
| sheet | 0.19 |
| loop | -1.02 |
| Map CC (mask) | 0.71 |
| Map CC (box) | 0.78 |
| Map CC (peaks) | 0.62 |
| Map CC (volume) | 0.70 |
| Map CC for ligands | 0.75 |
