## Supplementary Table 2 for "LetA defines a structurally distinct transporter family involved in lipid trafficking"

| Reagent/Resource | Description | Reference | Addgene ID |
|---|---|---|---|
| OverExpress™ C43(DE3) | *E. coli* competent cells for protein expression (#CMC0019) | Sigma-Aldrich | |
| Rosetta(DE3) | *E. coli* competent cells for protein expression (#70954) | Novagen | |
| TOP10 | *E. coli* competent cells for plasmid propagation (#C404010) | ThermoFisher | |
| bBEL466 | *E. coli* K-12 BW25113 *ΔletAB* | Isom *et al*., 2020 | |
| bBEL385 | *E. coli* K-12 BW25113 *ΔpqiAB* | Isom *et al*., 2020 | |
| bBEL384 | *E. coli* K-12 BW25113 *ΔpqiAB ΔletAB* kanR | Isom *et al*., 2020 | |
| bBEL620 | *E. coli* K-12 BW25113 *ΔpqiAB ΔletA* | This study | |
| bBEL621 | *E. coli* K-12 BW25113 *ΔpqiAB ΔletB* | This study | |
| bBEL609 | *E. coli* K-12 BW25113 *ΔpqiAB ΔletAB* | This study | |
| pET17b-letAB | LetAB for complementation | Isom *et al*., 2017 | 175804 |
| pBEL1284 | His-LetA-LetB for expression | Vieni *et al*., 2022 | 175960 |
| pBEL2071 | LetA-LetB (deoverlapped) complementation vector | Isom *et al*., 2020 | 139928 |
| pBEL2214 | [LetA-QHQ-7xHis] expression vector | This study | 232486 |
| pBEL2399 | [LetA(C31A)]-LetB complementation vector | This study | 232487 |
| pBEL2400 | [LetA(C34A)]-LetB complementation vector | This study | 232488 |
| pBEL2421 | [LetA(C233A)]-LetB complementation vector | This study | 232489 |
| pBEL2422 | [LetA(C236A)]-LetB complementation vector | This study | 232490 |
| pBEL2472 | [LetA(Δ2-24)]-LetB complementation vector | This study | 232491 |
| pBEL2577 | [6xHis2xQH-TEV-LetA(K328A)]-LetB expression vector | This study | 232492 |
| pBEL2516 | [6xHis2xQH-TEV-LetA(D367A)]-LetB expression vector | This study | 232493 |
| pBEL2518 | [6xHis2xQH-TEV-LetA(K178A)]-LetB expression vector | This study | 232494 |
| pBEL2606 | [6xHis2xQH-TEV-LetA(W267Bpa)]-LetB expression vector | This study | 232495 |
| pBEL2630 | [LetA(C248A)]-LetB complementation vector | This study | 232496 |
| pBEL2631 | [LetA(C251A)]-LetB complementation vector | This study | 232497 |
| pBEL2634 | [LetA1(1-223)]-[LetA2(224-427)]-LetB complementation vector | This study | 232498 |
| pBEL2635 | [LetA(Δ2-17)]-LetB complementation vector | This study | 232499 |
| pBEL2636 | [6xHis2xQH-TEV-LetA]-LetB(F468Bpa) expression vector | This study | 232500 |
| pBEL2638 | [6xHis2xQH-TEV-LetA]-LetB(E854Bpa) expression vector | This study | 232501 |
| pBEL2658 | [LetA(C51A)]-LetB expression vector | This study | 232502 |
| pBEL2659 | [LetA(C54A)]-LetB expression vector | This study | 232503 |
| pBEL2745 | [6xHis2xQH-TEV-LetA(W162Bpa)]-LetB expression vector | This study | 232504 |
| pBEL2772 | [LetA(ΔZnRN)]-LetB complementation vector | This study | 232505 |
| pBEL2782 | [LetB-6xHis2xQH] expression vector | This study | 232506 |
| pBEL2792 | [LetA(ΔZnR$^{N}$ΔZnR$^{C}$)]-LetB complementation vector | This study | 232507 |
| pBEL2800 | [LetB(F468Bpa)-6xHis2xQH] expression vector | This study | 232508 |
| pBEL2802 | [6xHis2xQH-TEV-LetA1(1-223,C124S)]-[LetA2(224-427,C266S,C343S)]-LetB | This study | 232509 |
| pBEL2809 | [6xHis2xQH-TEV-LetA1(1-223,L94C,C124S)]-[LetA2(224-427,C266S,C343S,I383C)]-LetB | This study | 232510 |
| pBEL2819 | [6xHis2xQH-LetA(ΔZnR$^{N}$ΔZnR$^{C}$)]-LetB expression vector | This study | 232511 |
| pBEL2820 | [6xHis2xQH-TEV-LetA1(1-223,Q180C,C124S)]-[LetA2(224-427,C266S,C343S,R380C)]-LetB | This study | 232512 |
| pBEL2864 | [LetA1(1-223, C124S)]-[LetA2(224-427,C266S, C343D)]-LetB complementation vector | This study | 232513 |
| pBEL2865 | [6xHis2xQH-TEV-LetA1(1-223)]-[LetA2(224-427)-StrepTagII]- LetB expression vector | This study | 232514 |
| pBEL2870 | [LetA1(1-223,L94C,C124S)]-[LetA2(224-427,I383C,C266S, C343S)]-LetB complementation vector. | This study | 232515 |

| | | | |
|---|---|---|---|
| pBEL2871 | [LetA1(1-223,Q180C,C124S)]-[LetA2(224-427,R380C,C266S,C343S)]-LetB complementation vector. | This study | 232516 |
| pBEL2874 | [6xHis2xQH-TEV-LetA1(1-223,L94C,C124S)]-[LetA2(224-427,C266S,C343S)]-LetB | This study | 232517 |
| pBEL2875 | [6xHis2xQH-TEV-LetA1(1-223,Q180C,C124S)]-[LetA2(224-427,C266S,C343S)]-LetB | This study | 232518 |
| pBEL2879 | [LetA(ΔZnR$^{C}$)]-LetB complementation vector | This study | 232519 |
| pBEL2886 | [LetB] expression vector | This study | 232520 |
| pBEL2889 | [6xHis2xQH-TEV-LetA1(1-223,C124S)]-[LetA2(224-427,C266S,C343S,I383C)]-LetB | This study | 232521 |
| pBEL2890 | [6xHis2xQH-TEV-LetA1(1-223,C124S)]-[LetA2(224-427,C266S,C343S,R380C)]-LetB | This study | 232522 |
| pBEL2913 | [6xHis2xQH-LetA(ΔZnR$^{N}$)]-LetB expression vector | This study | 232523 |
| pBEL2914 | [6xHis2xQH-LetA(ΔZnR$^{C}$)]-LetB expression vector | This study | 232524 |
| pBEL2921 | [6xHis2xQH-TEV-LetA(S364A)]-LetB expression vector | This study | 232525 |
| pBEL3014 | [6xHis2xQH-TEV-LetA(D181A)]-LetB expression vector | This study | 232526 |
| pBEL3015 | [6xHis2xQH-TEV-LetA(S321A)]-LetB expression vector | This study | 232527 |
| pBEL3016 | [6xHis2xQH-TEV-LetA(T402A)]-LetB expression vector | This study | 232528 |
| pBEL3017 | [LetA(PqiA-ZnR$^{N}$/PqiA-ZnR$^{C}$)]-LetB complementation vector | This study | 232529 |
| pBEL3158 | LetA-[LetB(R63A)] complementation vector | This study | 232530 |
| pBEL3159 | LetA-[LetB(R63N)] complementation vector | This study | 232531 |
| pBEL3160 | LetA-[LetB(R63D)] complementation vector | This study | 232532 |
