## Supplementary Table 3 for "LetA defines a structurally distinct transporter family involved in lipid trafficking"

**Supplementary Table 3.** Duration of each SMD simulation step

| | | SMD simulation steps | | | |
|---|---|---|---|---|---|
| | | Towards the bottom of the periplasmic pocket | Towards the middle of the periplasmic pocket | Towards the top of the periplasmic pocket | Total |
| Pulling the headgroup | +Lipid 1 | 96.9 ns | 53.1 ns | 85.8 ns | 235.8 ns |
| | -Lipid 1 | 81.0 ns | 54.7 ns | 95.0 ns | 230.7 ns |
| Pulling the terminal six carbons of the tail closest to the $TMD^C$ amphipathic groove | +Lipid 1 | 41.9 ns | 49.6 ns | 84.1 ns | 175.6 ns |
| | -Lipid 1 | 39.6 ns | 49.2 ns | 77.1 ns | 165.9 ns |
| Pulling the terminal six carbons of both tails | +Lipid 1 | 86.6 ns | 57.2 ns | 83.7 ns | 227.5 ns |
| | -Lipid 1 | 76.6 ns | 53.3 ns | 75.9 ns | 205.8 ns |
