## Supplementary Table 4 for "LetA defines a structurally distinct transporter family involved in lipid trafficking"

**Supplementary Table 4A.** Replicate 1

| Sample | Sample (µL) | Chloroform (µL) | Methanol (µL) | $H_2O$ (µL) | EquiSPLASH (µL) |
|---|---|---|---|---|---|
| Buffer | 85 | 170 | 45 | 68 | 45 |
| Protein | 40 | 80 | 0 | 32 | 40 |
| Membrane | 100 | 200 | 50 | 80 | 50 |

**Supplementary Table 4B.** Replicate 2

| Sample | Sample (µL) | Chloroform (µL) | Methanol (µL) | $H_2O$ (µL) | EquiSPLASH (µL) |
|---|---|---|---|---|---|
| Buffer | 50 | 100 | 25 | 40 | 25 |
| Protein | 50 | 100 | 25 | 40 | 25 |
| Membrane | 50 | 100 | 25 | 40 | 25 |

**Supplementary Table 4C.** Replicate 3

| Sample | Sample (µL) | Chloroform (µL) | Methanol (µL) | $H_2O$ (µL) | EquiSPLASH (µL) |
|---|---|---|---|---|---|
| Buffer | 30 | 60 | 15 | 24 | 15 |
| Protein | 30 | 60 | 15 | 24 | 15 |
| Membrane | 30 | 60 | 15 | 24 | 15 |

**Supplementary Table 4D.** Trap-and-elute LC gradient employed for DDA and MS1 data acquisition. Time points where no change occurs for a specific pump are marked with an asterisk (*).

| | **Alpha Pump** | | | | **Beta Pump** | | | |
|---|---|---|---|---|---|---|---|---|
| **Time (min)** | **Flow (mL/min)** | **%A** | **%B** | **Curve** | **Flow (mL/min)** | **%A** | **%B** | **Curve** |
| 0 | 1.000 | 70.0 | 30.0 | Initial | 0.250 | 60.0 | 40.0 | Initial |
| 0.5 | 1.000 | 60.0 | 40.0 | 6 | 0.250 | 60.0 | 40.0 | 6 |
| 1.5 | * | * | * | * | 0.250 | 50.0 | 50.0 | 6 |
| 2.5 | * | * | * | * | 0.250 | 40.0 | 60.0 | 6 |
| 2.9 | * | * | * | * | 0.250 | 34.0 | 66.0 | 6 |
| 4.5 | * | * | * | * | 0.250 | 32.3 | 67.7 | 6 |
| 4.51 | * | * | * | * | 0.250 | 32.2 | 67.8 | 11 |
| 12.00 | * | * | * | * | 0.250 | 3.0 | 70.0 | 6 |
| 13.00 | * | * | * | * | 0.250 | 10.0 | 90.0 | 6 |
| 15.00 | * | * | * | * | 0.250 | 10.0 | 90.0 | 6 |
| 15.50 | * | * | * | * | 0.250 | 1.0 | 99.0 | 6 |
| 17.00 | 0.300 | 1.0 | 99.0 | 11 | * | * | * | * |
| 17.50 | * | * | * | * | 0.250 | 1.0 | 99.0 | 11 |
| 20.00 | 1.000 | 70.0 | 30.0 | 11 | * | * | * | * |
